# Combinatorial perturbation analysis reveals divergent regulations of mesenchymal genes during epithelial-to-mesenchymal transition

**DOI:** 10.1101/627372

**Authors:** Kazuhide Watanabe, Nicholas Panchy, Shuhei Noguchi, Harukazu Suzuki, Tian Hong

**Affiliations:** RIKEN Center for Integrative Medical Sciences, 1-7-22 Suehiro-cho, Tsurumi-ku, Yokohama, Kanagawa 230-0045, Japan; Department of Biochemistry & Cellular and Molecular Biology. The University of Tennessee, Knoxville. Knoxville, Tennessee, 37996, United States of America; National Institute for Mathematical and Biological Synthesis. Knoxville, Tennessee, 37996, United States of America

## Abstract

Epithelial-to-mesenchymal transition (EMT), a fundamental transdifferentiation process in development, produces diverse phenotypes in different physiological or pathological conditions. Many genes involved in EMT have been identified to date, but mechanisms contributing to the phenotypic diversity and those governing the coupling between the dynamics of epithelial (E) genes and that of the mesenchymal (M) genes are unclear. In this study, we employed combinatorial perturbations to mammary epithelial cells to induce a series of EMT phenotypes by manipulating two essential EMT-inducing elements, namely TGF-β and ZEB1. By measuring transcriptional changes in more than 700 E- and M-genes, we discovered that the M-genes exhibit a significant diversity in their dependency to these regulatory elements and identified three groups of M-genes that are controlled by different regulatory circuits. Notably, functional differences were detected among the M-gene clusters in motility regulation and in survival of breast cancer patients. We computationally predicted and experimentally confirmed that the reciprocity and reversibility of EMT are jointly regulated by ZEB1. Our integrative analysis reveals the key roles of ZEB1 in coordinating the dynamics of a large number of genes during EMT, and it provides new insights into the mechanisms for the diversity of EMT phenotypes.

## Introduction

In epithelial-to-mesenchymal transition (EMT), a developmental program essential for morphogenic processes in embryogenesis and crucial in pathogenesis of malignant tumors, the cell-state transition occurs between two major states, epithelial (E) and mesenchymal (M), with well-characterized morphological features ^1–3^. In the classical EMT process during development, transition from E to M state is unidirectional and two phenotypes are mutually exclusive. However, in cancer cells, phenotypes induced by EMT can be diverse with multiple intermediate or metastable states ^3–8^ and the transition between E and M states is often bidirectional ^2,4^. Thus, E and M phenotypes are regulated primarily in a reciprocal fashion during classical EMT, but context-dependent regulations of E/M phenotypes were observed in nonconical or partial EMT. It is therefore challenging to obtain a comprehensive picture of molecular regulation, especially of reciprocity of E and M phenotypes.

EMT is characterized by a number of effecter genes each of which contributes to defining E or M phenotypes. Transcriptional profiling has been used to systematically measure such molecular phenotypes of EMT in a quantitative manner ^9,10^. In these studies, several hundreds of EMT-related genes were selected through meta-analysis and manual curation using the expressional as well as functional characterization. For example, Tan et al. applied machine learning to obtain a list of signature genes which can precisely predict aggressiveness of cancer cells ^9^. Such molecular approaches contributed to understanding correlation of EMT and disease phenotypes ^9,10^. Notably, EMT phenotypes with diverse transcriptional profiles has been observed in various pathological conditions ^9,10^. However, understanding regulatory mechanisms of wide variety of EMT signature genes, particularly the coordination of these genes during EMT, requires mechanistic studies in appropriate model systems.

Among a myriad of EMT-regulating factors discovered to date, TGF-β has been shown to be a potent EMT promoting signal ^11^, and ZEB1 is an EMT-inducing transcription factor that not only functions as a regulator for EMT program but is also involved in tumorigenesis ^12^. Although TGF-β induces ZEB1 expression ^13^, it is not clear whether ZEB1 can serve as an indispensable master regulator for TGF-β-induced EMT among other master EMT-TFs including SNAIL/SLUG and TWIST families ^2^, and whether TGF-β and ZEB1 are in a linear axis that controls the entire EMT program. In addition, TGF-β has been shown to play paradoxical (tumor-initiating and -suppressing) roles in cancer progression ^14,15^. Similarly, a poised chromatin configuration of ZEB1 promoter was shown to be tumorigenic, and ZEB1 can be both tumor-promoting and pro-apoptotic factors ^15,16^. These observations suggest that there are complex transcriptional programs activated by these two factors. However, the regulatory networks connecting these two factors to diverse transcriptional activities are not clear at the transcriptomic level.

In this study, we employed combinatorial perturbations to TGF-β and ZEB1 and created a series of EMT states in mammary epithelial cells. Using cells at these states of EMT, we applied transcriptomics, machine learning, mathematical modeling and live-cell imaging analyses to examine how the coordinated transition between E and M states are regulated. We identified three groups of M-genes that can be distinguished by their responsiveness to TGF-β and ZEB1 pathways and demonstrated the distinct biological impacts of the three M clusters in breast cancer patient survival and cell motility regulation. Surprisingly, high expressions of a cluster of M-genes that are strictly dependent on ZEB1 have significant association with good prognosis in breast cancer patients. Furthermore, using a mathematical model, we show that the reciprocity of EMT is synergistically controlled by TGF-β and ZEB1, and that the loss of this reciprocity transitions leads to partial EMT state with increased reversibility, which reduces the robustness of the destination state. Our results provide a holistic view of regulations of diverse mesenchymal genes during EMT, and they elucidate the mechanisms by which the cells ensure the coupling between E- and M-gene expressions in the transition. The classification of M-genes that we developed can be useful for the understanding of the diversity of EMT that is observed in various physiological and pathological conditions.

## Results

### Divergence of mesenchymal genes upon perturbations of TGF-β and ZEB1

To dissect the molecular events involved in switching E and M phenotypes during TGF-β-induced EMT, we first generated ZEB1 knockout (KO) clones of MCF10A cells using CRISPR/Cas9 genome-editing technology (**Supplementary Figure 1**, see **Methods** for details). The KO cells did not show any detectable phenotypes in the basal culture condition (**Supplementary Figure 2A**). TGF-β treatment induced suppression of a representative E marker E-cadherin (E-cad, encoded by *CDH1* gene) and activation of a representative M marker Vimentin (VIM) in WT cells (**Supplementary Figure 2B-C**), confirming that the E- and M-genes are reciprocally regulated during EMT. However, TGF-β failed to downregulate E-cad in KO cells while VIM was still upregulated to the similar extent with the WT cells (**Supplementary Figure 2B-C**). These results suggest that while ZEB1 is a potent EMT inducing transcription factor, its expression is dispensable for the induction of some M-genes.

To obtain a comprehensive view of the relative contribution of ZEB1 and TGF-β to EMT expression, we compared the MCF10A cells under 4 treatment conditions (TGF-β treated, TGF-β treated and ZEB1 KO, ZEB1 overexpressed, ZEB1 overexpressed and TGF-β inhibited) and their respective control conditions (8 conditions in total, **Figure 1A** and **Table 1**). ZEB1 overexpression and TGF-β signaling inhibition were performed by using doxycycline (DOX)-inducible system and TGF-β type1 receptor kinase inhibitor SB-431542, respectively. We examined the transcriptomes of the cells under these 8 conditions with Cap Analysis of Gene Expression (CAGE), a highly sensitive and quantitative transcriptome assay which detects activities of transcription start site (TSS) ^17,18^. We also defined 8 contrast conditions (described in **Table 2**) for the purpose of calculating log fold-change (logFC) to quantify differential expression under different regulatory regimes. We used a list of EMT genes curated from two sources: a set of 416 E- and M-genes annotated by Tan et al. ^9^ (see Methods) and additional 319 EMT genes without explicit E- or M-genes annotation (Zhao et al. ^10^). Overall, 60.6% of annotated EMT genes, exhibited significant differential expression (*q* < 0.05. See Methods for details) under at least one of the 8 contrast conditions, compared to the rest of the genome where only 10.0% were differentially expressed. The difference represents a significant enrichment of differently expressed genes in the annotated EMT set (Fisher’s Exact test, *p* < 2.2e-16), indicating that our set of EMT genes is more responsive to manipulation of TGF-β and ZEB1 than the genome in general.

**Figure 1.**
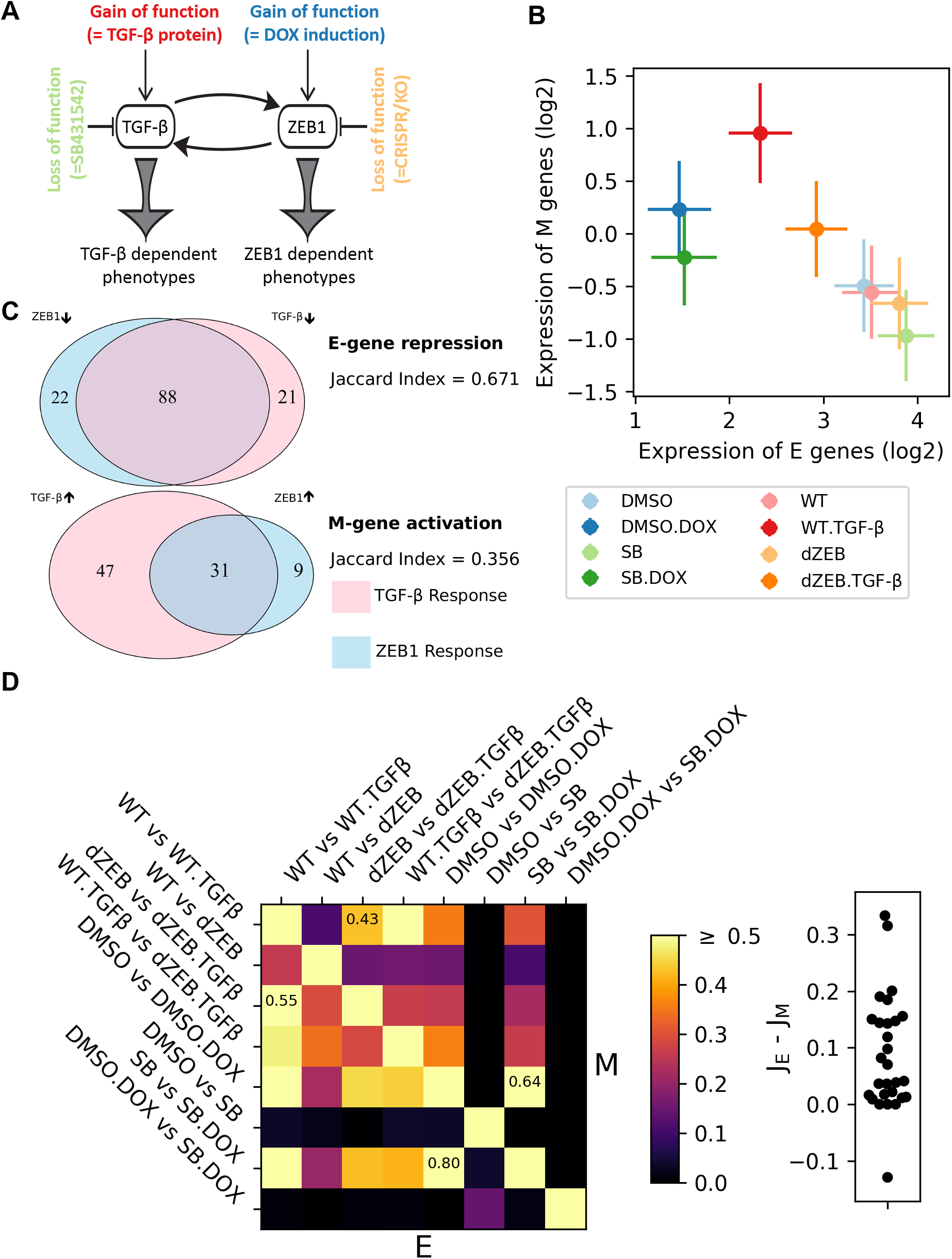
Quantification of EMT gene expression in response to TGF-β and ZEB1. **(A)** Illustration of perturbations of TGF-β and ZEB1 conditions in this study. Each colored perturbation has a control condition. **(B)** Mean expression levels of E- and M-genes of 8 conditions. Vertical and horizontal bars show standard error of the means for all annotated E or M genes. The colors are matched to (A). **(C)** Venn diagrams showing the overlap in E (top) and M (bottom) genes which show significant expression differences in response to TGF-β or ZEB1 treatment relative to their respective control conditions. TGF-β response is in pink while ZEB1 response is in light blue. The number E and M-genes responding to TGF-β or ZEB1 uniquely as well as those responding to both are listed in the respective part of the Venn diagram. The type of response, activation or repression, is indicated by directional arrows (up-arrow: activation, down-arrow: repression). The overlap was quantified using the Jaccard index, which is the number of genes differentially expressed by both TGF-β and ZEB1 divided by the total number of differentially expressed genes. **(D)** Jaccard indices of E- and M-genes for each pair of conditions. Heatmap shows all the Jaccard indices. Lower triangular entries: E-genes. Upper triangular entries: M genes. The Jaccard index for the conditions which differ most between E and M-genes are shown in those cells. Swarm plot shows the differences between the Jaccard indices of E-genes and those of the M-genes for each of the 28 pairs of conditions (*p* < 0.001 for single value t-test with a null distribution centered at 0).

**Table 1.** Combinatorial perturbation conditions

| Expression Condition | Description |
| --- | --- |
| WT | control |
| TGF-β | TGF-β treatment |
| dZEB | CRISPR-mediated ZEB1 knockout |
| TGF-β + dZEB | TGF-β treatment and ZEB1 knockout |
| DMSO | control for doxycycline-induction of ZEB1 and SB431542 |
| DOX | doxycycline-induction of ZEB1 |
| SB | TGF-β inhibition by TGF-β receptor kinase inhibitor SB431542 |
| DOX + SB | ZEB1 over-expression and TGF-β inhibition |

We next explored expression differences between genes associated with epithelial phenotypes (E-genes) and those associated with mesenchymal phenotypes (M-genes). As expected, E-genes have lower expression under TGF-β treatment or ZEB1 overexpression conditions, and higher expression when these factors are inhibited or knocked out, while M-genes show the inverse pattern (**Figure 1B and Supplementary Figure 3**), though there is obvious variation in the strength and level of response of each class of EMT genes to TGF-β or ZEB1 gain and loss. If we only consider cases of significant differential expression, 89.1% of E-genes were down-regulated under by TGF-β treatment (TGF-β vs. WT) or ZEB1 overexpression (DOX vs. DMSO), while 89.7% of M-genes were up-regulated in response to TGF-β treatment or ZEB1 overexpression. This is consistent with TGF-β/ZEB1 induction being responsible for a shift from epithelial phenotype to mesenchymal phenotype. As expected, the mean expression levels of E- and M-genes showed a negative correlation among the 8 conditions. However, the overexpression of ZEB1 had much stronger effect on E-genes than on M-genes (**Figure 1B**, the blue dots compared to the red dots), suggesting the primary role of ZEB1 in inhibiting E genes.

We next considered the response to TGF-β and ZEB1 independently. E- and M-genes exhibit different patterns of differential expression to TGF-β and ZEB1. Among E-genes, there is a large degree of overlap between genes which are down-regulated in response to both TGF-β and ZEB1 (67.1%), while almost twice as many M-genes are differentially expressed in response to TGF-β compared ZEB1, such that differential expression of most M-genes (64.4%) is specific to one factor or the other (**Figure 1C**). This difference in expression between E and M genes is robust to variations in the q-value and log fold-change thresholds that we used to define differential expression (**Supplementary Figure 4**). The overlap for E and that for M-genes become similar only with very large fold change thresholds (LogFC > 2) and even with this high threshold, E and M genes remain distinct as more E genes are differentially expressed in response to ZEB1 while more M genes are differentially expressed in response to TGF-β. In addition, we observed a difference between E-genes and M-genes in terms of the dependence of one EMT factor on the other one: there is an 80% overlap in E-genes that were both differentially expressed in response to ZEB1 induction (DMSO + DOX vs DMSO) and in response to ZEB1 induction in the absence of TGF-β (DOX + SB vs SB) and a 55% overlap between E-genes differentially expressed in response to TGF-β (WT + TGF-β vs. WT) and in response to TGF-β in the absence of ZEB1 (TGF-β + dZEB vs dZEB) (**Figure 1D**). This suggests that most E-genes are differentially expressed in response to both TGF-β and ZEB1, and the response to ZEB1 alone was stronger than that to TGF-β. Comparably, in M-genes these overlaps are only 64% for ZEB1-induced differential expressions, and 43% for TGF-β-induced differential expressions (**Figure 1D**). We also observed a distinction between E- and M-genes across EMT inducing factors: among E-genes that were differentially expressed in response to ZEB1 without TGF-β, 64% were differentially expressed in response to TGF-β with ZEB1 and 42% were differentially expressed in response to TGF-β without ZEB1. The corresponding values for M-genes were only 36% and 22% respectively. While the overlap between TGF-β and ZEB1 expression, both with and without the other factor, represents the largest differences in the amount of overlapping differential expression, E-genes are differentially expressed more uniformly than M-genes across all but a few comparisons (**Figure 1D**), an observation which is also robust to stricter definitions of differential expression (**Supplementary Figure 4**). Therefore, we conclude that M-genes exhibiting a divergent response to different combinations of ZEB1 and TGF-β input while E-genes are more uniformly responsive to the TGF-β-ZEB1 axis (**Figure 1D**). Furthermore, given both the divergence between TGF-β- and ZEB1-induced differential expression among M-genes as well as the fact that M-genes are less frequently differentially expressed by either TGF-β treatment or ZEB1 induction independently, we hypothesized the existence of multiple M-gene regulatory modules under the control of TGF-β and ZEB1.

### Three distinct types of regulatory circuits connect TGF-β and ZEB1 to M-genes

In an attempt to integrate all our expression profiles into single analysis, we first applied hierarchical clustering to logFC expression data (**Supplementary Figure 5**). While the resulting tree largely separates annotated E- and M-genes in two main clusters respectively, 19.5% of annotated EMT genes were incorrectly classified (19.2% E-genes, and 19.8% M-genes). Although some of this error may be the result of misannotated EMT genes, the difficulty in correctly separating E- and M-genes may in part be due to the high-dimensionality of our data set as well as the assumption that all conditions are equally important to the distinction between E- and M-genes.

To address this issue, we used a semi-supervised approach to classify the E- and M-genes, and to identify major classes of M-genes subsequently. We first applied a self-organizing maps (SOM) algorithm (see **Methods**) to map EMT genes onto a 10×10 grid based on their logFC of expression under the 8 contrast conditions. This dimensionality reduction clearly separated E-genes and M-genes (**Figure 2A**), with 87 of the 100 nodes dominated by one type of EMT gene (E:M ≥ 2 or M:E ≥ 2), and 65 consisting exclusively of E-genes or M-genes. Of the remaining 13 nodes, 5 contain a mixture of annotated E- and M-genes, while the remaining nodes lack EMT genes with E or M annotations (black nodes, **Figure 2A**), though this accounts for only 15 of the 319 (4.8%) EMT genes without an E or M annotation. Overall, 95.5% of EMT genes fall into a node with more than a 2:1 ratio of E/M or M/E genes. Using this cutoff, 7.9% of annotated EMT genes were incorrectly classified, and 87.6% of annotated EMT genes were correctly classified. Based on this classification, we further propagated E and M annotations to non-annotated EMT genes in nodes with predominant E- or M-gene annotations. Importantly, after updating our definition of E- and M-genes, we observe the same overall pattern of response to different combinations of TGF-β and ZEB1 expression, though there is slight reduction in responsiveness overall (**Supplementary Table 1**).

**Figure 2.**
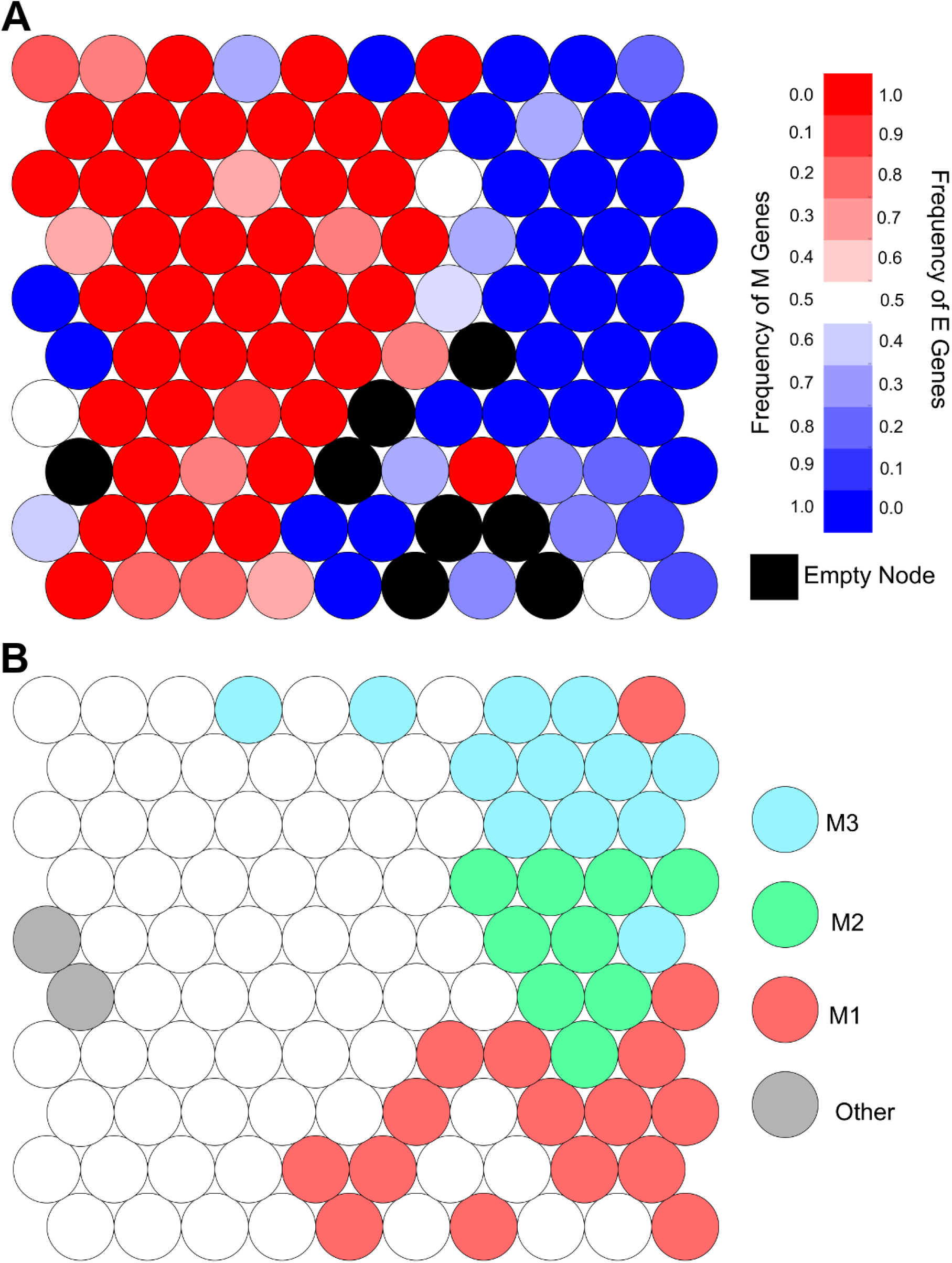
Self-organizing Maps (SOMs) for E-and M-genes and for clustering M-genes. **(A)** SOM nodes by frequency of E- and M-genes using a visual representation of the final map of EMT genes onto a 10-by-10 grid by SOM. The color of each node indicates the frequency of E-genes (darker-red) and M-genes (darker-blue) in each node. Nodes that are colored black are empty. **(B)** Clustering of SOM nodes with predominantly M-gene membership using a visual representation of the final map of EMT genes onto a 10-by-10 grid by SOM. Each of the 38 genes which have a 2:1 or greater ratio of M-genes to E-genes is colored according to the cluster of M-genes it was assigned to by hierarchical clustering (red = M1, green = M2, blue = M3, grey =other).

Based on the diversity in M gene regulation, we used nodes with predominant M-gene annotation for the further exploration of M-gene regulatory modules. We selected 38 nodes from our SOM grid with predominantly M-genes and applied hierarchical clustering and cut the resulting tree to generate four clusters (**Figures 2B** and **Supplementary Figure 6**. See **Methods**). Of the resulting clusters, three (‘M1’, ‘M2’, and ‘M3’ in Figure 2B) contain at least 70 genes overall and more than 30 annotated M-genes, but the smallest (‘Other’ in Figure2B) contained only 11 genes total and was therefore deemed too small for further analysis. The M1, M2, and M3 clusters together cover 86.2% of annotated M-genes while including only 9.4% of E-genes (see **Table 3**). To visualize expression of our M-gene clusters, we performed principal component analysis (PCA) with the logFC expression data (**Supplementary Figure 7**). We found that each M-gene cluster has a distinct pattern of expression, and we inferred the key perturbations that gave rise to the differences among the M-gene clusters by examining the loading (R) of individual principal components (PCs). In particular, PC1, which separates M1 and M3, is correlated with TGF-β response (R = 0.363 for WT + TGF-β vs. WT and 0.264 for dZEB + TGF-β vs. dZEB) and anticorrelated with ZEB1 response (R = −0.518 DMSO + DOX vs. DMSO and −0.541 for SB + DOX vs. SB), suggesting that M1 genes are more responsive to TGF-β and M3 genes are more responsive to ZEB1. Similarly, PC2 is correlated with ZEB1-independent TGF-β response (R = 0.653 dZEB + TGF-β vs. dZEB), suggesting M2 genes have a stronger response to TGF-β independent of ZEB1 than M1 and M3 genes do. While visualizing expression data in key directions is useful ^19^, we were unable to distinguish our three clusters of M-genes with PCA alone (compare the labeled plots to the unlabeled plots (**Supplementary Figure 7**, upper triangle).

We next performed further characterization of each group of M-genes. The M1 cluster covers almost half (48.9%) of M-genes, including the regulator SNAI1, fibroblast growth factor (FGF2) and fibroblast growth factor receptors (FGFR2), heat-shock proteins CRYAB and HSBP2, as well as several genes associated with tumor growth and migration including MMP9 ^20^, CDH11 ^21^, FOXC1 ^22,23^, and RAC1 ^24,25^. However, 25 of the M-genes in this cluster (13.3% of total M-genes) belonged to a single node comprised of genes that were not expressed across all data sets, and so were excluded from subsequent analysis (doing this also excluded 23% of the E-genes initially included in our M-gene clusters). These M1 genes were up-regulated in response to TGF-β treatment more than ZEB1 induction (Mann-Whitney U test, *p* < 2.2e-16), but the response of M1 genes to TGF-β was significantly reduced when ZEB1 was absent (median logFC changed from 0.83 to 0.14, *p* = 1.10e-10, Mann-Whitney U test) (**Figure 3A**). Therefore, M1 genes are upregulated by TGF-β, and the responses depend on ZEB1. However, neither of the factors has a strong influence on these genes alone. The next largest cluster M2 (20.7% of M-genes) includes the structural proteins FN1 and VIM, which are both associated with the mesenchymal cell phenotype ^26,27^ as well as the tumor suppressor genes CDKN1B (P27, ^28^), CDKN2A (P16, ^29^) and PTEN ^30^. Like M1 genes, M2 genes had greater response to TGF-β than to ZEB1 (Mann-Whitney U test, *p* = 0.02) (**Figure 3B**), but the difference in median response was much smaller for M2 genes (TGF-β = 1.01, ZEB1 = 0.57) than for M1 genes (TGF-β = 0.83, ZEB1 = 0.0, *p* = −2.56-e15). Furthermore, the response of M2 genes to TGF-β was not affected by the absence of ZEB1, (Mann-Whitney U test, *p* = 0.69) nor was its response to ZEB1 affected by the inhibition of TGF-β (Mann-Whitney U test, *p* = 0.33) (**Figure 3B**). As such, while TGF-β had a larger impact on M2 expression, both TGF-β and ZEB1 regulated M2 genes independent of one another. The M3 cluster (16.5% of M-genes) contains the transcription factors TWIST1 and TWIST2, growth factors HFG and FGFR1, two Mitogen-activated protein kinases, MAP3K3 and MAPK7, which are over-expressed in tumors ^31–34^, and the Insulin-like growth factor-binding IFGBP3 which as complication relationship with cancer progression depending on cancer type ^35–37^ M3 genes responded to ZEB1 more than TGF-β (Mann-Whitney U test, *p* = 2.32e-7) with no significant difference in ZEB1 response in the absence of TGF-β (Mann-Whitney U test, *p* = 0.09) (**Figure 3C**). M3 genes responded to TGF-β induction (median logFC = 0.35). This response was lost when ZEB1 was absent (median logFC = 0) and this change in response was significantly different (Mann-Whitney U-test, *p* = 7.18e-6). Therefore, we conclude that M3 genes are regulated by ZEB1 independent of TGF-β. To further confirm the differential expression patterns among the M-gene clusters, we performed RT-PCR for representative genes in each cluster and the results were consistent with our CAGE experiments and the clustering analysis (**Supplementary Figure 8**). The clustering information for all 735 EMT genes that we analyzed is listed in **Supplementary Table 2**. In general, the three largest clusters obtained from our analysis show distinct patterns of regulation by TGF-β and ZEB1, in contrast to E-genes which primarily respond to ZEB1 directly (**Supplementary Figure 9**). We summarized our findings in an illustrative model shown in **Figure 3D** which is largely reflective of the differences in expression suggested by our PCA analysis. In this model, M1 genes are regulated by TGF-β and ZEB1 via an AND logic gate. This AND-gate can only be turned on by TGF-β but not ZEB1, possibly because TGF-β can activate ZEB1 completely, but ZEB1 can only partially activate TGF-β signaling (**Figure 3D**, dashed arrow). The activation of TGF-β signaling by ZEB1 is supported by previous findings that show the mechanisms and importance of the mutual activation between TGF-β and ZEB1 ^38–40^. In particular, ZEB1 activates SMAD proteins which serve as key mediators of TGF-β signaling ^39,40^. In contrast to the M1 gene cluster, M2 genes are regulated by the two factors via an OR-gate, M3 genes are regulated by ZEB1 but not ZEB1-independent TGF-β pathway, and E-genes are assumed to be controlled by ZEB1. The latter assumption is based on the observation that 82% of E-genes were downregulated by the expression of ZEB1 alone. Note that the arrows in this simplified network diagram (**Figure 3D**) do not represent direct molecular interactions, but the diagram establishes causal, rather than correlative, relationships between the two core EMT factors and other M-genes because of the controlled perturbations that we performed.

**Figure 3.**
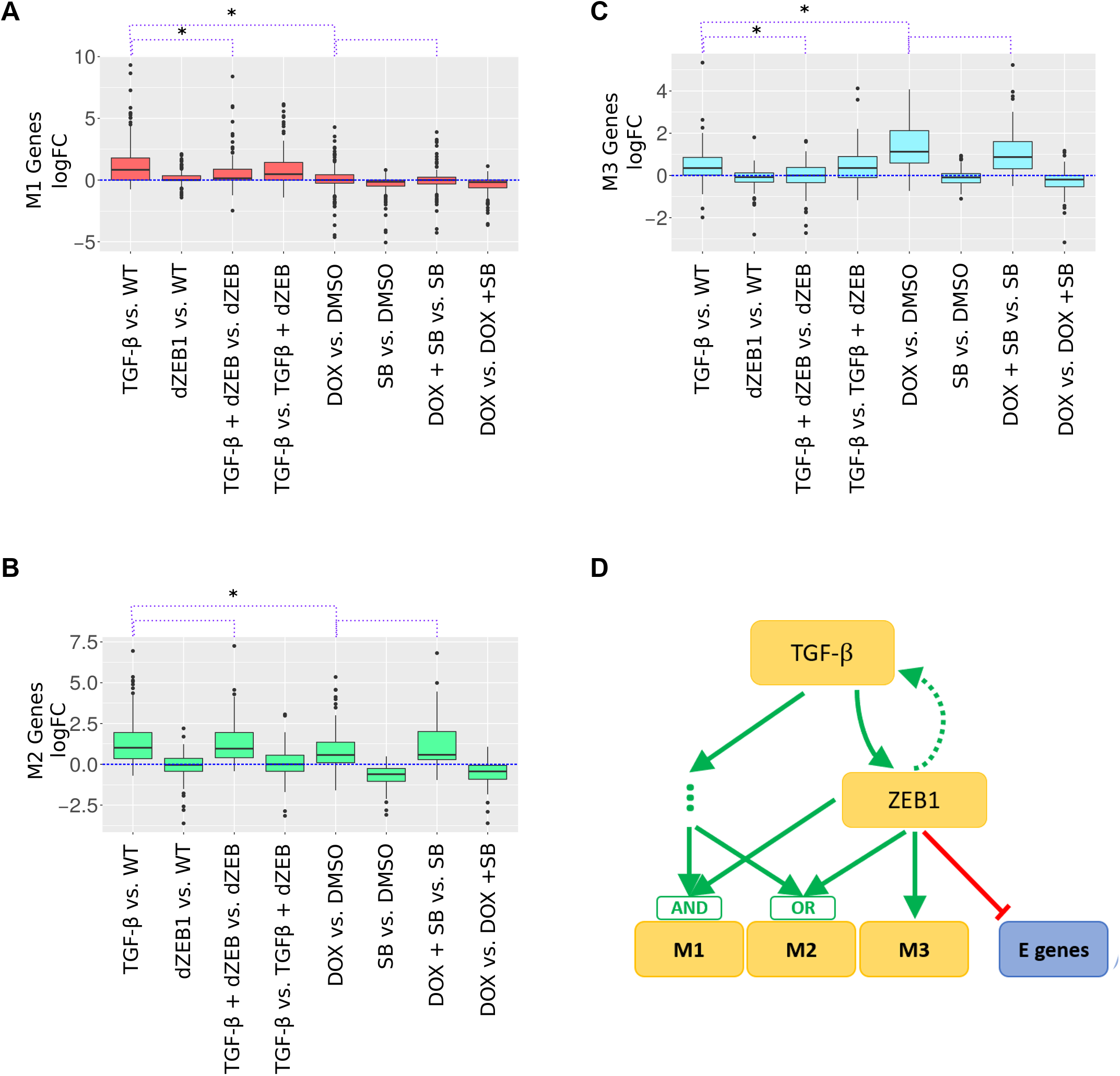
Expression of M-gene clusters under TGF-β and ZEB1 regulation. **(A)** Boxplot of log fold-change of expression of M1 genes in the 8 contrast conditions (see **Table 2**). The colored region (red) indicates the inter-quartile range of expression while whiskers extend 1.5 times this range on either side. Outliers are indicated by black dots. The purple dotted lines above the plot indicate comparisons between the expression under conditions, specifically, TGF-β vs. WT to DOX vs. DMSO, TGF-β vs. WT to TGF-β + dZEB vs. dZEB and DOX vs. DMSO to DOX + SB vs. SB. A * indicates that distribution of expression is significantly different based on the Mann-Whitney U-test at an alpha of 0.05. **(B)** Similar to **(A)** but for M2 genes, with a green colored region. **(C)** Similar to **(A)**, but for M3 genes, with a blue colored region. **(D)** A model of EMT-gene regulation based on the expression patterns of M-genes clusters in **(A-C)**. Green arrows indicate activation while red arrows indicate repression.

We tested if the M-gene clusters are differentially regulated by other EMT inducers using a set of previously published microarray data ^41^ (see **Supplementary Text 1, Supplementary Table S3**). We found that M2 and M3 genes have a significantly larger response to EMT inducing factor Gsc than M1 genes do (**Supplementary Figures 10 and 11**). Future work involving systematic and controlled perturbations of EMT factors will be required to elucidate the regulation of M-genes by other core EMT transcription factors.

### M-gene clusters have distinct biological functions

To investigate the functional difference between the three M-gene groups, we retrieved human GO annotation from Gene Ontology Consortium and identified significantly enriched terms within each group of M-genes compared to the genome overall using Fisher’s Exact test (*p* < 0.05, see **Methods**). M1 and M2 genes both had 85 significantly enriched terms while M3 genes had only 29. However, the majority of these terms are uniquely enriched to one group of M-genes, with only six terms found in all groups and 67, 66, 18 appearing exclusively in M1, M2 and M3 genes respectively (**Figure 4A**). In comparison, there are 169 GO terms are uniquely enriched among E-genes (**Supplementary Table 4**). However, when we categorized E genes into three groups by the responsiveness to ZEB1 and TGF-β (as in Figure 1C), neither E genes responsive to ZEB1 or to TGF-β alone were enriched for any GO terms, while those responsive to both were enriched for 20 terms, primarily related to cell-cell junctions and adhesion (**Supplementary Table 5**). Among the set of unique genes in each group, we identified several sets of GO terms related to the same overarching function, which we have exhibited in **Figure 4A**. In particular, we highlight the contrast between M1 and M2 genes: although both groups are enriched for term related to apoptosis, M1 genes are enriched for several terms related to cell motility, adhesion and proliferation while M2 genes are enriched for terms relating to the negative regulation of the same processes. Additionally, M2 are enriched for a wide variety development-associated GO terms not found in M1, including circulatory system, nervous system, embryonic and other organs. This indicates a distinction between classes of M-genes: M1 genes are associated with the mobility and proliferative qualities associated with EMT, and M2 genes are related to development functions.

**Figure 4.**
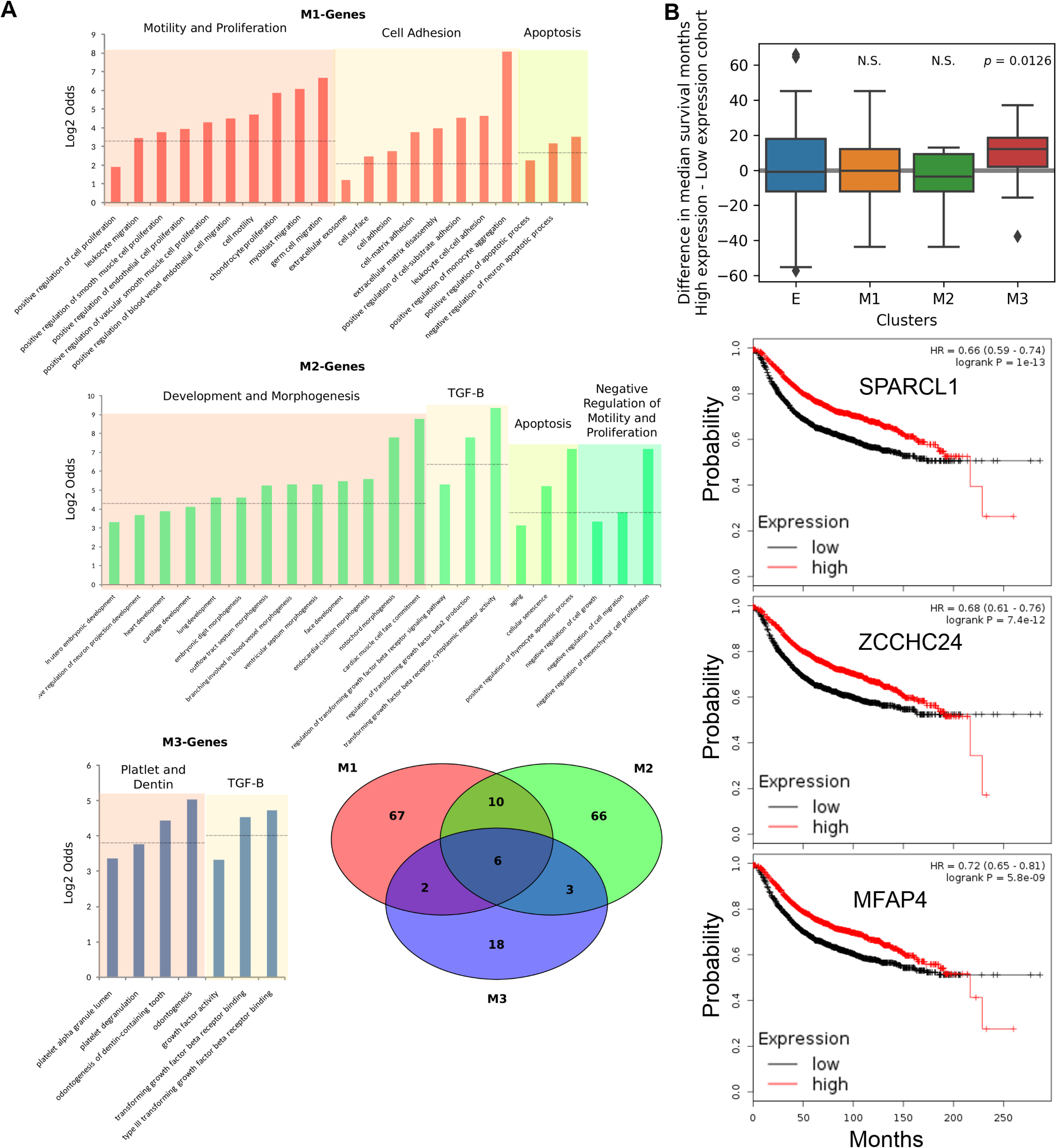
Functional annotations and survival analysis of M clusters. **(A)** Bar graphs show groups of related GO terms associated with each cluster of M-genes. Venn diagram shows the overlapped GO terms (*p* < 0.05 for selection of GO terms) among the three M clusters. **(B)** Top: Differences between high expression and low expression cohorts in terms of median survival months for breast cancer patients. Each group represents one type (cluster) of genes. Single-value *t*-test was performed with each group of median survival months. Bottom: Kaplan-Meier plots for three representative M3 genes. Survival months and Kaplan-Meier plots were obtained from KM-Plotter ^46^.

In addition to normal cellular functions, EMT is also strongly implicated in cancer progression, especially in breast cancers ^26,42–45^. To check whether the three clusters of M-genes have differential roles in prognosis of breast cancer patients, we performed survival analysis for all the EMT genes annotated in this and earlier studies ^46^ (see **Methods** for details). We found that there is no significant difference between patients with low and high expressions of E, M1 and M2 genes in terms of the median survival months (**Figure 4B**), indicating the overall complexity of EMT’s role in cancer progression. However, high expression of M3 genes is significantly associated with better prognosis compared to low expression of the corresponding genes in the breast cancer patients (**Figure 4B**. See **Supplementary Table 2** for the full list of median survival months for the EMT genes). The Kaplan-Meier plots for three representative genes are shown in **Figure 4B**. Among these genes, SPARCL1 was identified as a tumor suppressor gene ^47^. ZCCHC24 is strongly correlated with sensitivity to drug treatment ^48^. MFAP4 is downregulated in several types of cancer and it was recently suggested to be a marker for developing therapies ^49^ These results suggest that the high expression of the genes that are strictly dependent on ZEB1 pathway may play protective roles in cancer progression, or they can be served as markers for improved prognosis. To exclude the possibility that the association between better prognosis and high expression of M3 genes was driven by a few outlier genes, we calculated the percentages of genes of which higher expressions are significantly associated better prognosis for each cluster. Consistent with the boxplot and the t-test (**Figure 4B**), M3 cluster has higher percentage of such genes (45.7%) than any other cluster does (M1: 33.6%, M2: 22.5%, E: 30.7%). These results are consistent with several recent findings that challenge the simple association between mesenchymal state and the invasiveness of cancer: higher expression of certain mesenchymal genes is associated with better prognosis, or inversely, higher expression of certain epithelial genes is associated with worse survival ^9,50,51^. For instance, the E-gene GRHL2 (**Supplementary Table 2**) was shown to correlate with poor survival across all subtypes of breast cancer ^52^.

We next performed gene set enrichment analysis to further explore the significance of the M-gene clusters in cancer settings. We found that these clusters are differentially expressed across a variety of cancer types (see **Supplementary Text 2**), suggesting that co-expression of M-gene cluster occurs in cancer. M3 genes specifically are uniquely enriched among genes differentially expressed in luminal A breast cancer and papillary thyroid cancer (**Supplementary Table 6**). However, the set of differentially expressed genes tends to be small (between 4% and 28% of the cluster), so such relationship between M-gene clusters and cancer is likely driven by sub-clusters of cancer related genes. We performed further analysis on the expression of the M-gene clusters in mesenchymal-like cancer cell lines ^53^ (**Supplementary Text 3**). We found that the mean expressions of the three M-gene clusters are significantly different in these cancer cell lines: M2 has the highest expression level and M1 has the lowest expression level (**Supplementary Figure 12**). In addition, M3 genes have stronger within-cluster correlations (M3 genes vs. M3 genes) than between-cluster correlations (M3 genes vs. M1/M2 genes) in these cells (**Supplementary Figures 12** and **13**). These results suggest the significance of the M-gene clusters in cancer cells.

### Differential cell movement patterns regulated by TGF-β and ZEB1

To explore the relation between gene clusters and cellular phenotypes controlled by TGF-β and ZEB1, we collected imaging data for MCF10A cells under the 8 conditions listed in **Table 1** and analyzed the movement patterns of the cells for each condition (**Figure 5A**). We used four metrics to quantify the cell movement for each trajectory: the mean instantaneous velocity, the displacement scaled by duration, the straightness index and the lifetime-averaged number of nearest neighbors (see **Methods**). Overexpression of ZEB1 significantly increased the velocity of the cells, whereas the influence of TGF-β treatment on velocity is less significant (**Figure 5B**, top panel). Nonetheless, inhibition of TGF-β signaling had the most prominent negative effect on the velocity of the movement. In contrast, increasing TGF-β or ZEB1 signal had significant positive effect on displacement, which quantifies the overall migration efficiency of the cells. ZEB1 overexpression was the most potent condition to increase displacement, while the presence of TGF-β signaling pathway was also essential for the increase (**Figure 5B**, panel 2). ZEB1 alone positively influenced the straightness of the cell movement, even in the absence of the TGF-β signaling pathway, whereas the influence of TGF-β on straightness depends on the presence of ZEB1 (**Figure 5B**, panel 3). ZEB1 also had a predominant role in reducing the number of neighboring cells during the lifetime of the trajectories (**Figure 5B**, bottom panel), suggesting that its positive regulation of the straightness of the movement is partially via the reduction of cell-to-cell contact. These results demonstrate the key role of ZEB1 in regulating the straightness of the cell movement, which is correlated with the overall efficiency of cell migration.

**Figure 5.**
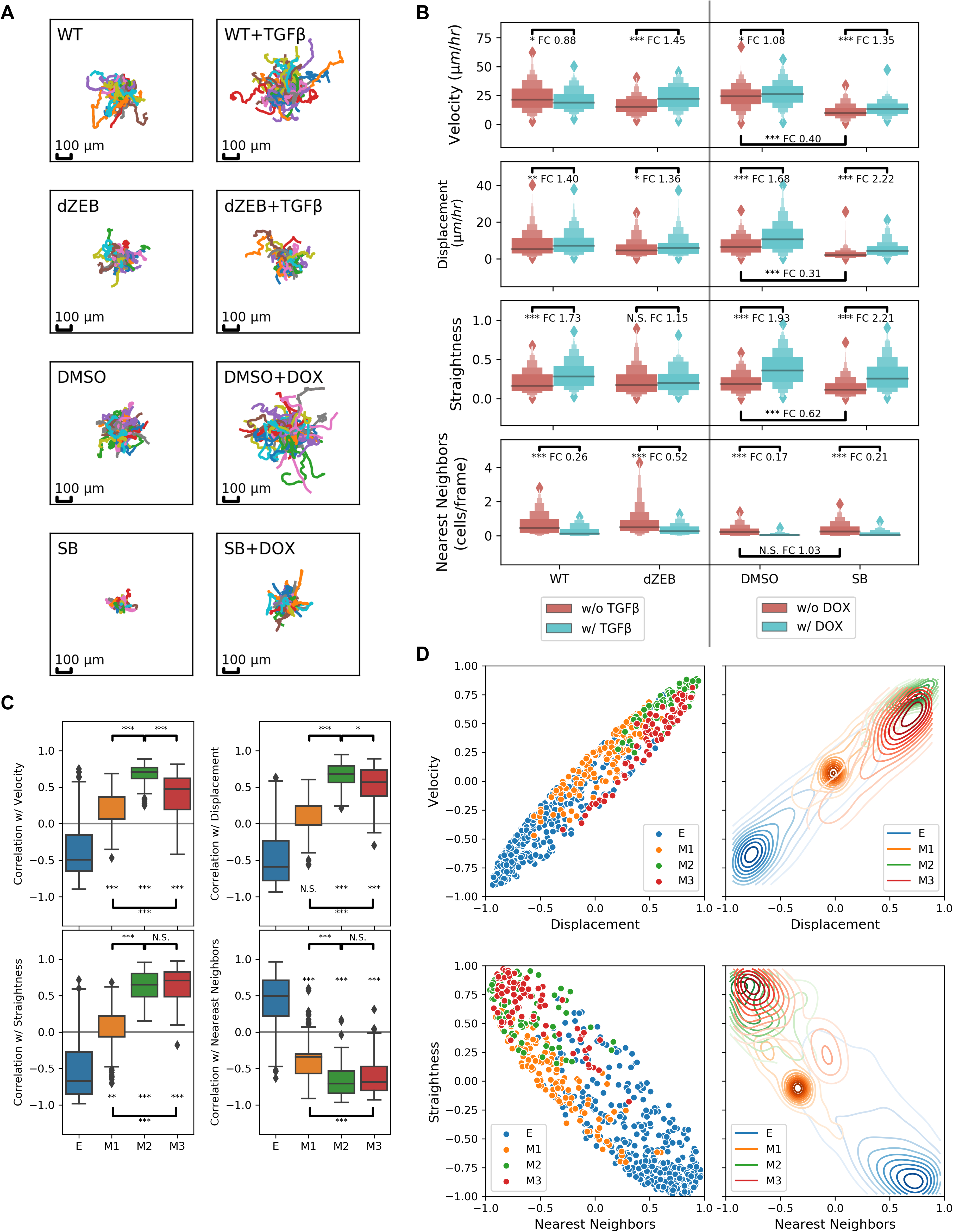
Differential cell movement patterns regulated by ZEB1 dependent and independent pathways. **(A)** Cell movement trajectories when TGF-β signaling and/or ZEB1 expression is perturbed under 8 conditions. 100 cells were randomly selected for each condition. Each trajectory was centered at its starting position. **(B)** Distributions of four metrics (instantaneous velocity, mean displacement normalized by duration of trajectory, straightness index of the movement and number of nearest neighbors) for cell trajectories under 8 conditions. Statistical significance was obtained using Mann-Whitney U test. FC: fold-change. **(C)** Distributions of Spearman correlations coefficients (as a distance measurement) between gene expression and movement metrics for four type of genes (E, M1, M2 and M3) across 8 conditions. **(D)** Scatter plots showing pairwise relationships between correlation coefficients of gene expression and different movement metrics. ***:*p* < 0.001, **:*p* < 0.01, *:*p* < 0.05, N.S.: not significant (*p* > 0.05). Significant mark at each box indicates the *p* value for testing if the median of the group is significantly different from 0. Significant mark at each horizontal bar indicates the *p* value for testing if two groups of values are significantly different.

We next asked how the expression of M-gene clusters is correlated with the movement patterns. We calculated the Spearman correlation coefficients between the gene expression levels across the 8 conditions and values of each of the movement metrics described above under the same conditions. These correlation coefficients serve as distance measurements between the gene expression pattern of each gene and movement pattern. For example, a positive coefficient between the expression of a gene and displacement means that the higher expression of that gene is correlated with the higher displacement. Among the three M-gene clusters, the expression of M2 genes has the strongest correlation with velocity, displacement, straightness and nearest neighbors (all four metrics of movements). Compared with M2 genes, the expression of M3 genes has weaker, but still significantly positive correlation with velocity, and comparable correlations with all other three metrics. This suggests that M2 and M3 genes have similar contributions to the overall movement patterns. We asked under which specific conditions M2 gene expression shows better correlation with the velocity than the expression of other M-genes does, and we found that when cells were treated with TGF-β in the absence of ZEB1, velocity was significantly increased, and this is the condition under which M3 genes, but not the other two groups of genes, were significantly (>2-fold) upregulated (**Supplementary Figure 14**). In contrast to M2 and M3 genes, the expression of M1 genes is not significantly correlated with the displacement, and its correlations with other movement metrics are much weaker than that of the expression of M2 and M3 genes. These weak correlations are consistent with the differential sensitivities of cell movement patterns and M1 gene expression to EMT signals: the movement of the cells is sensitive to perturbations to either TGF-β or ZEB1, whereas M1 genes can only be upregulated when both signals are present. Nonetheless, the significant correlation between the expression of M1 genes and some movement patterns is consistent with the GO analysis (**Figure 4A**).

### A mathematical model for ZEB1-TGF-β transcriptional network reveals the role of ZEB1 in controlling reciprocity and reversibility of EMT

To gain more insights into the roles of ZEB1 in controlling EMT, we built a mathematical model to describe the gene regulatory network for EMT in response to TGF-β signal based on the three groups of M-genes and one E-gene group (**Figure 6A**). In particular, M1 genes are activated by TGF-β and ZEB1 via an AND logic gate, M3 genes are directly activated through ZEB1-dependent pathway but not ZEB1-independent pathway, and M2 genes are influenced by both ZEB1-dependent and ZEB1-independent pathway via an OR logic gate. We assumed that the E-genes are primarily controlled by ZEB1-dependent pathway, as suggested by the expression analysis of E genes in earlier sections (**Supplementary Figure 9**). In addition, we included some known feedback loops involving ZEB1 that were established in earlier studies, including a positive feedback loop between ZEB1 and TGF-β ^38–40^, and a representative mutual inhibition loop formed by ZEB1 and another factor (e.g. miR200, OVOL2 and GRHL2, ^4,54,55^. Note that all of these typical E-genes were correctly classified as E-genes with our algorithm. See **Supplementary Table 2**.).

**Figure 6.**
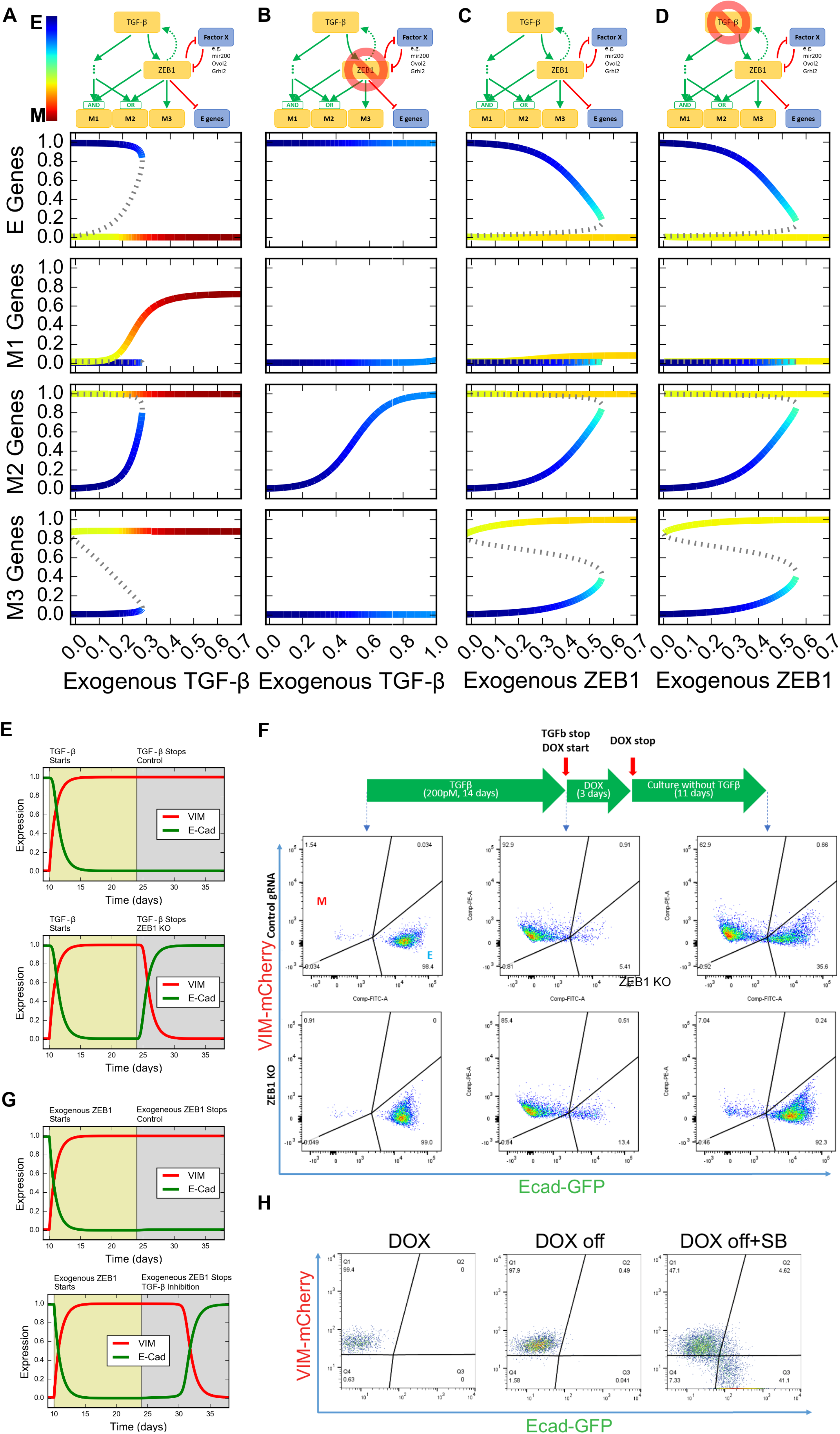
Mathematical modeling of EMT under control of TGF-β and ZEB1. **(A-D)**. Top diagrams: influence diagrams for gene regulatory networks under four conditions. Lower panels: bifurcation diagram for four types of genes (E, M1, M2 and M3) with respect to exogenous TGF-β and ZEB1 expression under two conditions. Solid curves represent stable steady states. Dashed curves represent unstable steady states. Color gradient represents the position in the EMT spectrum, which is calculated by adding the expression of the three M nodes and subtracting the expression of the E node. **(A)** Normal condition. Induced by TGF-β. **(B)** ZEB1 KO. Induced by TGF-β. **(C)** Normal condition. Induced by ZEB1. **(D)** TGF-β inhibited condition. Induced by ZEB1. **(E)** Simulation for expression of VIM and E-cad upon treatment and withdrawal of TGF-β. Top: control. Down: ZEB1 KO after TGF-β withdrawal. **(F)** Expression of VIM and E-cad analyzed by FACS upon treatment and withdrawal of TGF-β. Top: control. Bottom: ZEB1 knockout induced by DOX. Cells were treated with TGF-β for two weeks and then subject to TGF-β withdrawal, and (for the experiment group) to ZEB1 knockout. **(G)** Simulations for expression of VIM and E-cad upon treatment and withdrawal of exogenous ZEB1 expression followed by inhibition of TGF-β signaling. **(H)** Expression of VIM and E-cad analyzed by FACS upon induction and withdrawal of exogenous ZEB1 by DOX. Exogenous ZEB1 was induced for one week and then subject to the withdrawal of the induction signal for two weeks, and (for the experiment group) to the inhibition of TGF-β signal by SB431542 (SB) for the same period.

We performed bifurcation analysis with the model by varying two parameters: the strength of external TGF-β signal, which represents physiological inducer of EMT and the concentrations of the exogenous ZEB1 which was controlled experimentally in this study. With the increasing strength of external TGF-β, the production of E-genes was turned off, and the production of all types of M-genes was turned on (**Figure 6A**). The ‘flipping’ from E-on-M-off state to E-off-M-on state shows a reciprocal regulation of E- and M-genes under normal conditions. In addition, the model suggests that the TGF-β induced EMT is irreversible once the cells have committed to the complete M state, and this is consistent with earlier mathematical models and experimental observations ^7,56^. Note that in this study we define reversibility of EMT as the ability for the system to return to the E state upon the complete withdrawal of EMT signal TGF-β or ZEB1 (transition from red/yellow branch to blue branch with EMT signal decreasing to zero in **Figure 6A-D**). It is possible to examine the reversibility of EMT upon partial withdrawal of EMT signal but this property is more accurately described as hysteresis instead of (ir)reversibility ^57^. We next blocked the production of ZEB1 in the model and performed similar bifurcation analysis. In the absence of ZEB1, the TGF-β signaling only triggered the transition into a partial EMT state, in which only M2 was upregulated whereas the ZEB1-dependent M1 and M3 genes were not responsive (**Figure 6B**). In addition, the model predicts that TGF-β induced EMT becomes reversible upon the loss of ZEB1 (**Figure 6B and E**).

Exogenous expression of ZEB1 triggered reciprocal regulation of E-genes and some M-genes: with increasing ZEB1 production rate, E-genes were downregulated whereas M2 and M3 genes were upregulated (**Figure 6C**). However, exogenous ZEB1 did not activate M1 genes because of the absence of exogenous TGF-β. Nonetheless, the model suggests that ZEB1 can trigger irreversible EMT (**Figure 6C**). In addition, the model predicts that the inhibition of TGF-β can reduce the irreversibility (**Figure 6D and G**).

To validate the predictions in terms of the reversibility (**Figure 6E**), we first verified that TGF-β-induced EMT is irreversible in most WT cells at least for 10 days without continuous exposure to TGF-β using an EMT reporter system (See **Method, Figure 6F**). In contrast, the EMT phenotype induced by TGF-β was reversed by inducing ZEB1 deletion using an inducible genome editing system with DOX-inducible Cas9 and constitutive expression of ZEB1-targeting gRNA (**Figure 6F**). Furthermore, transient induction of ZEB1 expression triggered irreversible EMT (**Figure 6H**, left and middle panels), which is consistent with our modeling analysis. Interestingly, the irreversibility of the transition is even more robust than that of the TGF-β-induced EMT (**Figure 6H**, middle panel, **Figure 6F**, top right panel) 10 days after signal withdrawal. Treatment with TGF-β inhibitor upon withdrawal of exogenous ZEB1 expression caused a partial reversal of the EMT (**Figure 6H**, right panel). These results suggest that the endogenous ZEB1 with its feedback regulation is essential for maintenance of irreversible EMT phenotype in mammary epithelial cells as reported for other cell types ^38^. Together with the results shown in earlier sections, our data indicate that the irreversibility and the reciprocity of EMT are both regulated by ZEB1, and these two properties may be closely related.

## Discussion

EMT is involved in many biological processes, but a remarkable diversity of EMT phenotypes has been observed in both distinct and similar pathological conditions ^9,10,44,58^. This diversity may be attributed to the metastable states that exist between terminal E and M states ^2,4,5,7^. To describe these multiple states, a linear lineage progression model with coupled changes of E- and M-gene expression was widely used ^2,4,58^. In this study, we identified the key roles of ZEB1 in regulations of the coupling between E- and M-genes. In addition, our findings raised the possibility that genetic or microenvironmental perturbations on ZEB1 activity may result in decoupling of E and M phenotypes, which may contribute to the diverse ‘hybrid’ EMT populations observed previously ^58^. Furthermore, our analysis of the expression of EMT genes in response to ZEB1 and TGF-β not only indicates that M-genes exhibit a greater diversity of responses, but also suggests that they can be classified into no less than three sub-clusters. M-gene clusters have distinct patterns of expression in response to different perturbations of ZEB1 and TGF-β, and they can also be separated based on their functions, with M2 exhibiting development-related function, while M1 genes are involved in cell motility, growth and adhesion. Furthermore, M3 genes have a significant association with the prognosis of breast cancer, indicating that genes that are controlled by ZEB1 but not other TGF-β-dependent pathways likely play a protective role. Although ZEB1 was shown to promote tumor initiation and metastasis ^59^, the importance of the poised ZEB1 promoter status in cancer progression ^16^ indicates that there is a subset of ZEB1 induced genes that may be inversely correlated with tumorigenesis. This may account for the differential roles of multiple EMT states in progression of cancer ^60^. The existence of these functionally distinct sub-groups of M-genes further elucidates the connection between decoupling and hybrid EMT phenotypes because the disruption of normal regulation may not affect all processes involved in EMT equally. Our study focused on the diversity of M-genes instead of E-genes, because M-genes are more diverse than E-genes in terms of their responsiveness to EMT inducing factors (**Figure 1**). However, it is possible that E-genes exhibit significantly albeit more weakly divergent responses, and they may be diversely regulated by factors other than ZEB1 and TGF-β. Future work is warranted to test these possibilities.

Our model elucidates the key roles of ZEB1 in regulating the reciprocity of EMT. During EMT, ZEB1 is directly responsible for the downregulation of most E-genes and the upregulation of a group of M3 genes. Effectively, ZEB1 ensures the coupling between the loss of the E phenotypes and the gain of M phenotypes. Conversely, the absence of ZEB1 blocks the ability of the cells to transition to terminal M state, and it decouples the dynamics of E- and M-genes. In addition, a ZEB1-independent pathway that activates M1 genes is also required for the complete transition to the terminal M state. Taken together, our model provides a mechanistic view of the EMT transcriptional network controlled by ZEB1 and TGF-β at the transcriptomic level. Our results also demonstrated the importance of ZEB1 in controlling the reversibility of EMT. This is consistent with a very recent study which showed the essential roles of ZEB1-miR200 feedback loop in the hysteresis of EMT, and that the varied reversibility of EMT can influence the metastatic potentials of cancer cells ^57^. Our results further suggest that ZEB1 serves as the hub for coordinating the reciprocity of E- and M-genes, and this coupling is closely related to the variable reversibility of EMT. In our mathematical model, we took the simple assumption that ZEB1 forms a positive feedback (double-negative) loop with another E-gene. In fact, ZEB1 may form positive feedback (double-positive) loops with other M-genes, such as SNAIL or TWIST as well, and these feedback loops may also contribute to the irreversibility of EMT. With the positive feedback loops involving ZEB1, E- and M-genes are reciprocally regulated, and their dynamics are always inversely correlated when the extracellular signal is varied (**Supplementary Figure 15A**). This reciprocity is also essential for the irreversibility of the EMT. Without the feedback loops (e.g. loss of ZEB1), the reciprocity of E- and M-genes are compromised, and EMT becomes more reversible because the lack of feedback that requires the downregulation of E-genes (**Supplementary Figure 15B**).

Controversial findings have been reported for roles of EMT in cancer metastasis ^61–64^ Accumulating evidence supports that sequential induction of EMT and its reverse process mesenchymal-to-epithelial transition (MET) allows cancer cells enter into the systemic circulation and subsequently colonize at distant organs ^2,65^. On the other hand, other studies suggested that cancer metastasis is often caused by circulating clusters of epithelial tumor cells and that EMT is dispensable for this process ^62,63^. These conflicting findings suggest two modes of spreading processes of cancer cells; migratory and invasive properties of individual cancer cells that are related to EMT phenotypes and collective cell migration which occurs without losing epithelial integrity. The latter type of collective migration may not fit with the classical linear lineage progression definition of EMT. In fact, recent theoretical work suggests the importance of hybrid EMT states for circulating tumor cell clusters ^66^. Our findings suggest that E and M phenotypes are not necessarily regulated in a reciprocal manner and that EMT process can be flexibly reversed in these cells. It has been suggested that such plastic states are related to cancer stem cell phenotypes ^8,16,67^. Furthermore, several reports suggested that sequential activation of EMT and MET promotes reprograming or differentiation of cell lineages including iPS, neurons, or hepatocyte lineages ^68–70^. Our study provides a possible molecular basis for such plastic transition of cellular state and manipulating the balance between the two inhibitory networks may be useful to develop new treatment for diseases or novel cell conversion methods.

Previous mathematical models and experiments showed the existence of intermediate EMT states *in silico*, *in vitro* and *in vivo* ^4,5,7,58^. In contrast, the model and the experiment presented in this study focus on the diversity of M-genes in terms of their connectivity with TGF-β and ZEB1, as well as the role of ZEB1 in controlling reversibility and reciprocity of EMT. However, the model does not describe the intricate feedback loops in the EMT transcriptional network. These feedback loops were shown to be critical for the formation of intermediate cell states ^4,56,71^. Therefore, the current model has limited predictive power in terms of the detailed transitions involving intermediate EMT states. In particular, the bifurcation point shown in **Figure 6A-D** may be decomposed into several consecutive switches that govern the critical transitions of different E- or M-genes. Future work is needed to integrate different elements of the EMT control circuits into a unified model to analyze the system in a more comprehensive manner. In addition, previous models predicted discrete states between E and M phenotypes ^4,5,7,40,56^, whereas our model and another EMT model by Celià-Terrassa et al. ^57^ suggest that continuous EMT phenotypes may be observed with increasing EMT signals. Distinguishing these two scenarios requires future experiments. Nonetheless, our model and experiment demonstrated the existence of hybrid EMT state at which both E- and M-genes are upregulated. It is unclear whether the ‘intermediate’ states can be distinguished from the ‘hybrid’ states under physiological conditions, and whether the intermediate phenotypes observed *in vivo* are transitional cells ready to commit to the destination states, or those unable to make the complete transition due to genetic or environmental perturbations and likely to be reverted to the initial states. More modeling work and single-cell experiments are warranted to address these questions, and the insights into these problems will help elucidating the functional roles of the intermediate or hybrid EMT states ^72^.

Our study provides a holistic view of the roles of ZEB1 and its interplay with TGF-β signaling at transcriptomic level. We found the key roles of ZEB1 in regulating both the reciprocity and reversibility of EMT, and we identified three classes of mesenchymal genes that are controlled by three different types of regulatory circuits downstream of TGF-β and ZEB1. These results shed light into the complex molecular mechanisms for regulating EMT, and they are useful for the understanding of the diversity of EMT phenotypes observed in many physiological and pathological conditions.

## Materials and Methods

### Cell lines

MCF10A cells (ATCC) were grown in DMEM/F12(1:1) medium with 5% horse serum, epidermal growth factor (10 ng/mL), cholera toxin (100 ng/mL) and insulin (0.023 IU/mL). For TGF-β treatment, cells were incubated with titrated concentrations of human TGF-β1 protein (R&D systems) in the complete culture medium. The culture medium was replaced daily, and cells were passaged right before reaching full confluency.

### CRISPR/Cas9-mediated ZEB1 deletion

CRISPR/Cas9-mediated genome editing of ZEB1 locus was performed using lentiviral gRNA expression system with lentiGuide-Puro (a gift from Feng Zhang Addgene plasmid #52963) and lentiCas9-Blast (a gift from Feng Zhang Addgene plasmid #52962). For inducible Cas9 expression and genome editing, Lenti-X™ Tet-One™ Inducible Expression System (Clontech) was used. Production of lentiviruses was carried out as previously described ^73^. Two gRNA sequences were used to delete ZEB1 expression; TGAAGACAAACTGCATATTG (tgg: PAM sequence) and CAGACCAGACAGTGTTACCA (ggg: PAM sequence) and the following gRNA sequences were used as controls: ACCAGGATGGGCACCACCC and GGCCAAACGTGCCCTGACGG. For ZEB1 KO clones, two clones (clone2 and clone5) were established showing complete absence of ZEB1 protein upon TGF-β stimulation (**Supplementary Figure 2A**). As the morphology, proliferation rates and gene expression patterns are similar, we chose to use clone5 for the following studies. This clone contained homozygous 370-bp deletion in Intron1-Exon2 boundary which results in exon skipping and frameshift (**Supplementary Figure 1B**).

### Inducible expression of ZEB1 protein

Puromycin-resistant Tet-based inducible cDNA expression system (pSLIK-Puro) was engineered by replacing the hygromycin-resistant gene in pSLIK-Hygro vector (a gift from Iain Fraser, Addgene plasmid # 25737) with a puromycin-resistant gene obtained from lentiGuide-Puro by PCR amplification. Mouse Zeb1 cDNA was digested from the pHIV-ZsGreen-Zeb1 lentiviral construct ^73^ and cloned into the pSLIK-Puro vector. Production and infection of lentiviruse were carried out as described above. Induction of the transgene was performed by doxycycline (DOX) treatment at a concentration of 500 ng/mL.

### EMT reporter and flow cytometry

The EMT reporter was engineered by removing the puromycin-resistant gene from a TCGP-Puro lentiviral EMT reporter system which contains E-cad promoter-driven eGFP and VIM promoter-driven mCherry (a kind gift from Kiyotsugu Yoshikawa). The EMT reporter-expressing MCF10A clone was established by infection and FACS sorting of eGFP positive cells, followed by serial dilution cloning. Several clones were screened by flow cytometry and clones that showed most distinguishable FACS profiles before and after TGF-β treatment was selected for the downstream analyses. Flow cytometry was performed on a BD FACSAria equipped with FACS DiVa6.0 software operating. Cell clusters and doublets were electronically gated out. Positive and negative gates for eGFP and mCherry fluorescence were determined using untreated and ZEB1-induced MCF10A cells as controls.

### RT-PCR

Total RNA was isolated using the TRIzol Reagent (Invitrogen) followed by cleaning up and RNase-free DNaseI treatment using the RNeasy mini kit (QIAGEN). cDNA was prepared using Retroscript Kit (Applied Biosystems) according to manufacturer’s instructions. Real-time PCR was performed using using a StepOnePlus™ real-time PCR system (Thermo Fisher), with SYBR Premix Ex Taq™ II (Takara). Comparative analysis was performed between the genes of interest normalized by the house keeping genes *GAPDH* and *ACTB*. The primer sequences used in this study are described in **Supplementary Table 7**.

### Cap analysis of gene expression (CAGE)

CAGE libraries were prepared as previously described ^17^. Briefly, 3 μg of total RNA from each sample were subjected to reverse transcription, using SuperScript III Reverse Transcriptase (Thermo Fisher) with random primers. The 5-end cap structure was biotinylated by sequential oxidation with NaIO4 and biotinylation with biotin hydrazide (Vector Laboratories, USA). After RNase I treatment (Promega, USA), the biotinylated cap structure was captured with streptavidin-coated magnetic beads (Thermo Fisher). After ligation of 5′ and 3′ adaptors, second-strand cDNA was synthesized with DeepVent (exo–) DNA polymerase (New England BioLabs, USA). The double-stranded cDNA was treated with exonuclease I (New England BioLabs) and purified. The resulting CAGE libraries were sequenced using single-end reads of 50 bp on the Illumina HiSeq 2500 (Illumina, USA). The extracted CAGE tags were then mapped to the human genome (hg38). After filtering low-quality reads, mapped CAGE tags were counted regarding FANTOM5-CAT TSSs ^18^, providing a unit of CAGE tag start site. The tags per million (tpm) were calculated for each TSS peak and computed as gene expression levels for multiple TSS peaks associated with a single gene. Differentially expressed genes were identified using QL F-test implemented by EdgeR, followed by false discovery rate control with Benjamini-Hochberg method.

### Clustering EMT genes with a semi-supervised approach

For the unsupervised step, two approaches were used to cluster EMT genes based on the logFC of expression under the 8 contrast conditions described in **Table 2**. logFC data was row-scaled to adjust for the large difference in average expression across EMT genes. Both approaches were implemented in the R programming language. In the first approach, a matrix of distances between annotated E- and M-genes was calculated based on logFC expression data using the ‘dist’ function with the Euclidean method. The ‘hclust’ function was then used to generate the dendrogram seen in **Supplementary Figure 5**. In the second approach, we use the ‘som’ function, which is part of the kohonen package ^74^, to map EMT genes onto a 10×10 grid of hexagonal cells. SOM were performed using the ‘som’ function. The full data set was presented to the network 1000 times during the learning process and the learning rate was set to decline linearly from 0.05 to 0.01 over the course of the learning process. From the final grid, we selected nodes which predominantly (2:1) consisted of M-genes (this supervised method is essentially a k-nearest-neighbors algorithm). We then clustered these selected nodes using the hierarchical clustering method described above, replacing logFC data with the codebook vectors describing each node. The ‘cutree’ function was used to derive clusters from the resulting dendrogram and we selected four as optimal numbers of clusters using the within cluster sums of squares error using the elbow criterion (**Supplementary Figure 6**).

**Table 2.**
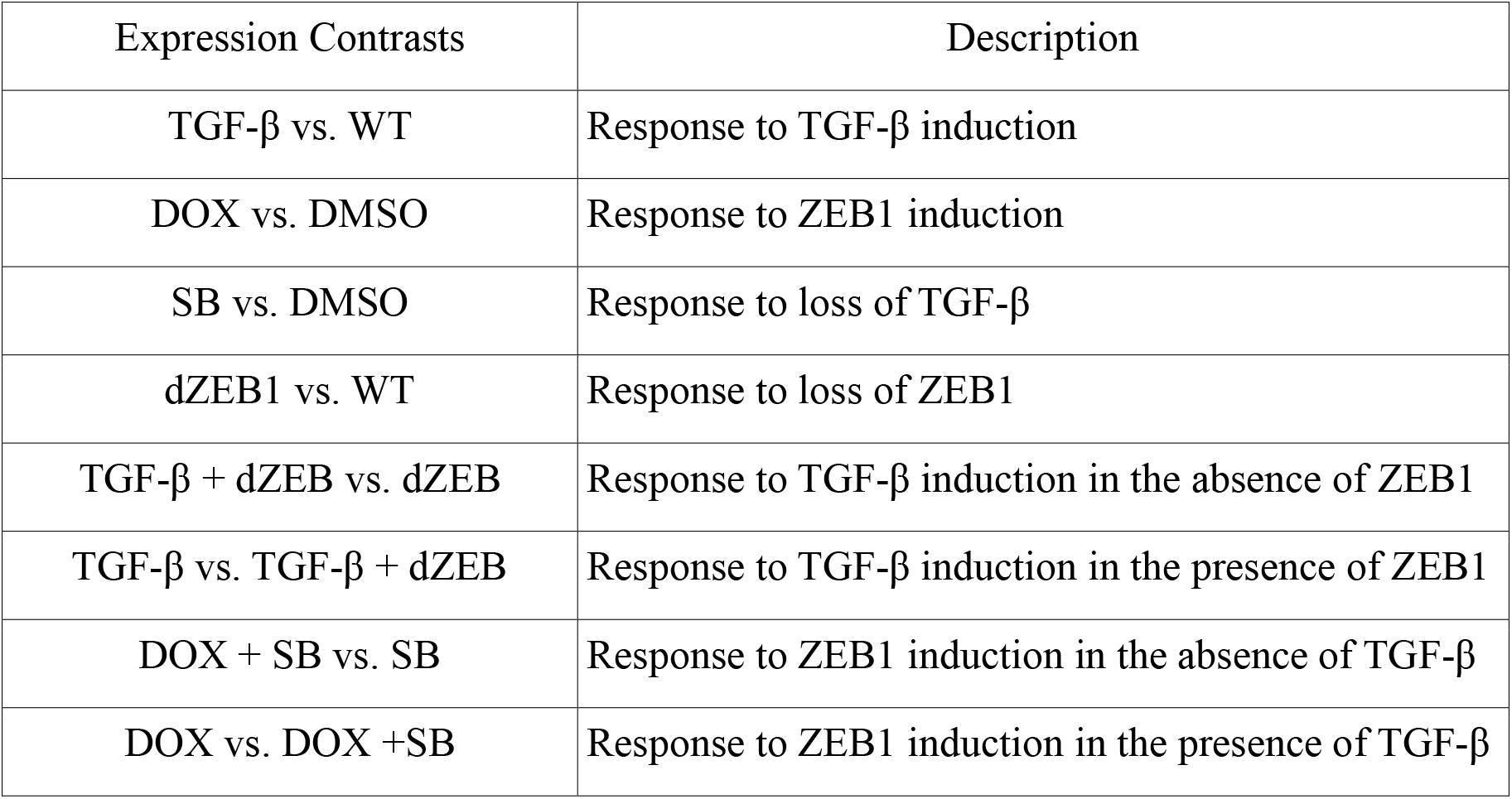
Contrast between expression conditions

| Expression Contrasts | Description |
| --- | --- |
| TGF-β vs. WT | Response to TGF-β induction |
| DOX vs. DMSO | Response to ZEB1 induction |
| SB vs. DMSO | Response to loss of TGF-β |
| dZEB1 vs. WT | Response to loss of ZEB1 |
| TGF-β + dZEB vs. dZEB | Response to TGF-β induction in the absence of ZEB1 |
| TGF-β vs. TGF-β + dZEB | Response to TGF-β induction in the presence of ZEB1 |
| DOX + SB vs. SB | Response to ZEB1 induction in the absence of TGF-β |
| DOX vs. DOX +SB | Response to ZEB1 induction in the presence of TGF-β |

**Table 3.** Size and membership of the three M-genes expression clusters

| Cluster # | Total Genes | M-genes | E-genes | Notable Genes |
| --- | --- | --- | --- | --- |
| 1 | 180 | 92 (48.9%) | 16 (6.8%) | SNAI1 |
| 2 | 80 | 39 (20.7%) | 1 (0.4%) | FN1, VIM |
| 3 | 77 | 31 (16.5%) | 4 (1.7%) | TWIST1, TWIST2 |

### Identifying enriched GO annotations in M-genes

GO annotation were obtained from Gene Ontology Consortium ^75,76^. The significance of enrichment of individual terms in each of the M-gene clusters was evaluated using Fisher’s Exact test and multiple test correction was implemented using the Benjamini-Hochberg correction.

### Survival analysis

Survival analysis was performed with KM-plotter ^46^. Patients of all breast cancer types were selected. Out of 735 EMT genes, probes for 720 genes were identified and analyzed. Medium survival months for high expression and low expression cohorts, as well as the p-values for the significance of their differences, were obtained from the website. We corrected the p-values using Benjamini-Hochberg procedure with false discovery rate (FDR) of 0.05. Genes without significant difference in survival months were discarded for subsequent analysis (assuming zero difference for these genes produced very similar results). Differences in survival months for genes in each M clusters were aggregated, and *t*-tests were performed to compare means of the differences to 0.

### Statistical tests and boxplots

Unless otherwise indicated, all *p* values were obtained with two-sided *t*-test assuming unequal variances. Other tests include two-sided Fisher’s Exact test (FET) for count data and two-sided Mann-Whitney U test for continuous numerical data with distributions far from normal. In all boxplots, center lines indicate median values, box heights indicate the inter-quartile range of data, whiskers extend 1.5 times this range on either side, and outliers are indicated by black dots.

### Mathematical modeling

We used a gene regulatory network that is simplified from earlier models ^4,5,7^. We incorporated the effector E-genes and three types of M-genes in the network, and their regulations by TGF-β and ZEB1 are based on the analysis from this study (**Figures 2 and 3**). To describe the system mathematically, we used a generic form of ordinary differential equations (ODEs) suitable for describing both gene expression and molecular interaction networks ^77–79^. Each ODE system in the model has the form:

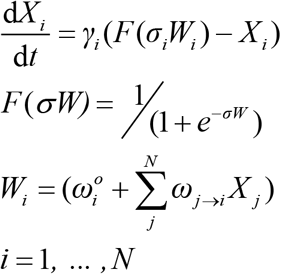

Here, *X_i_* is the activity or concentration of protein *i*. On a time scale 1/*γ_i_, X_i_*(*t*) relaxes toward a value determined by the sigmoidal function, *F*, which has a steepness set by *σ*. The basal value of *F*, in the absence of any influencing factors, is determined by 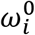. The coefficients *ω*_*j→i*_ determine the influence of protein *j* on protein *i. N* is the total number of proteins in the network. All variables and parameters are dimensionless. One time unit in the simulations corresponds to approximately 1 day.

To model the AND logic gate on M1 genes regulated by TGF-β and ZEB1, we assumed that the total influences of TGF-β (*ω_TGFβ→M1_TGFβ*) on M1 and ZEB1 (*ω_ZEB1→M1_ZEB*1) on M1 are both saturated at values less than 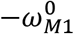, so that these signals do not activate M1 genes alone.

Steady state analysis was performed by varying the parameters representing exogenous TGF-β or ZEB1 production rate. The total value of the state variables representing the M-genes (M1, M2, M3) was used to quantify the M phenotype, and the value of the ‘E-genes’ was used to quantify the E phenotype. The difference between the phenotype was used to determine the position of the state in the EMT spectrum.

Parameter values were selected to fit to the observation that TGF-β induced EMT is irreversible under normal conditions. These values are listed in **Supplementary Table 8**.

### Imaging analysis

MCF10A cells were transduced with nuclear RFP-expressing lentivirus (LV-RFP, a gift from Elaine Fuchs, Addgene plasmid #26001) and treated under conditions listed in **Table 1**. Cell movement dynamics in 2D culture was monitored and recorded by IncuCyte® for 40 hours. Using binarized images, cells were identified and tracked by a Fiji package TrackMate. Cell trajectories longer than 24 frames (6 hours) were used for analysis. Movement analysis was performed using similar metrics described in an earlier study ^80^. Instantaneous velocity was computed as 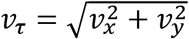 where *v_x_* = (*x_τ_* − *x*_*τ*−1_)/(*t_τ_* − *t*_*τ*−1_). Here, *x_τ_* is the *x* coordinate at time *τ*, and *t*_*τ*_ − *t*_*τ*−1_ is the time inverval between frames (15 min). The mean instantaneous velocity was calculated for each cell, and they were aggregated and compared across condtions. Scaled displacement was calculated with *SD* = |*x*(*t_end_*) − *x*(*t_start_*)|/(*t_end_* − *t_start_*), where *x*(*t_start_*) and *x*(*t_end_*) are the initial and final positional vectors of the trajectory respectively. Straightness index was calculated with

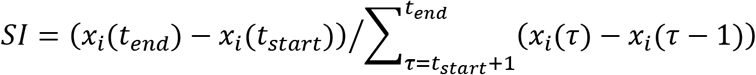

or the ratio of the distance between the initial and final positions for each cell to the integrated distance traveled. The mean number of nearest neighbors was computed for each cell in each frame by counting the other cells within a 30 μm search radius. This value was then divided by the totoal number of frames of each trajectory. These summary staticstics of the trajectories were compared between pairs of conditions listed in **Table 2**. Mann-Whitney U test was performed to obtain statisitcal significance.

## Data Availability

Sequencing data generated for this work have been deposited in the NCBI Gene Expression Omnibus (GEO) under accession number GSE124843. Computer code to reproduce the results of self-organizing maps, clustering and mathematical modeling is available upon request.

## Supplementary Materials

Supplementary Text 1. Regulation of M-gene clusters by other EMT regulators

Supplementary Text 2. Gene set enrichment analysis of M-gene clusters

Supplementary Text 3. Expression of M-gene clusters in mesenchymal-like cancer cell lines

Supplementary Figure 1. Establishment of ZEB1-deficient MCF10A cells.

Supplementary Figure 2. Role of ZEB1 in reciprocity of E and M phenotypes in EMT.

Supplementary Figure 3. Expression of EMT genes in the 8 combinations of TGF-β and ZEB1 treatments.

Supplementary Figure 4. Quantification of EMT gene expression in response to TGF-β and ZEB1 under different definitions of differential expression.

Supplementary Figure 5. Clustering E- and M-genes according to log fold-change of expression. Supplementary Figure 6. Elbow plot showing the changes of sum of squared errors as the number of clusters increases.

Supplementary Figure 7. Principal Component Analysis of CAGE expression data for M-genes.

Supplementary Figure 8. Comparison of expression patterns representative genes in four clusters using CAGE and RT-PCR.

Supplementary Figure 9. Expression of E-gene cluster under TGF-β and ZEB1 regulation. Supplementary Figure 10. Significance and confidence intervals for difference in odds of enrichment between M-gene groups across different regulatory data sets.

Supplementary Figure 11. Log fold-change of expression of each M-gene cluster in response to GSC overexpression.

Supplementary Figure 12. Expression of three cluster of M-genes in cancer cell lines.

Supplementary Figure 13. Positive correlations among M3 genes.

Supplementary Figure 14. Correlation between changes of mean gene expression and velocity of cells under 8 contrast conditions.

Supplementary Figure 15. Metaphoric illustration of relationship between reciprocity of E- and M-genes and irreversibility of EMT transition.

Supplementary Table 1. Overlapping EMT gene expression before and after annotating dbEMT genes.

Supplementary Table 2. List of EMT genes analyzed in this study and their clustering information.

Supplementary Table 3. Enrichment of EMT genes among gene upregulated by EMT regulators in Tauble et al.

Supplementary Table 4. List of GO Terms enriched in all epithelial genes.

Supplementary Table 5. List of GO Terms enriched in epithelial genes which are differentially expressed in response to both TGF-β and ZEB1.

Supplementary Table 6. Gene sets enriched in M-gene clusters.

Supplementary Table 7. List of primer sequences.

Supplementary Table 8. Parameter values for mathematical models.

## Acknowledgments

The authors would like to thank Ms. Hajime Nishimura and Ms. Mami Kishima for their technical assistance in preparing this article and Dr. Kiyotsugu Yoshikawa for kindly providing the TCGP reporter construct. The authors thank Ms. Esha Dutta for the assistance with survival analysis.

## Author Contributions

Conceived and designed the experiments: KW, TH. Performed the experiments and modeling: KW, NP, TH. Analyzed the data: KW, NP, SN, HS, TH. Wrote the manuscript: KW, NP, TH.

