## Supplemental files for "Combinatorial perturbation analysis reveals divergent regulations of mesenchymal genes during epithelial-to-mesenchymal transition"

**SUPPLEMENTARY INFORMATION**

SUPPLEMENTARY TEXTS…………………………………………………………………………………………………4

**Supplementary Text 1. Regulation of M-gene clusters by other EMT regulators….……..**4

**Supplementary Text 2. Gene set enrichment analysis of M-gene clusters……….…..…..**7

**Supplementary Text 3. Expression of M-gene clusters in mesenchymal-like cancer cell lines…………………………………………………………………………………………………….**8

SUPPLEMENTARY FIGURES………………………………………………………………………………………10

**Supplementary Figure 1.** Establishment of ZEB1-deficient MCF10A cells……………………10

**Supplementary Figure 2**. Role of ZEB1 in reciprocity of E and M phenotypes in EMT…..11

**Supplementary Figure 3.** Expression of EMT genes in the 8 combinations of TGF-β and ZEB1 treatments…………………………………………………………………………………………….13

**Supplementary Figure 4.** Quantification of EMT gene expression in response to TGF-β and ZEB1 under different definitions of differential expression…………………………………...14

**Supplementary Table 5.** List of GO Terms enriched in epithelial genes which are differentially expressed in response to both TGF-β and ZEB1……………………………………….49

**Supplementary Table 6.** Gene sets enriched in M-gene clusters………………………………….50

**Supplementary Table 7.** List of primer sequences…………………………………………………….53

**Supplementary Table 8.** Parameter values for mathematical models………………………….54

REFERENCES…………………………………………………………………………………………………………………55

### SUPPLEMENTARY TEXTS

#### Supplementary Text 1 - Regulation of M-gene clusters by other EMT regulators

To further investigate the regulation of M-gene clusters, we employed microarray expression data generated by Taube et al. [1] for four transcription factors identified as being upstream EMT inducers (TGF-β, Snail, Twist, and Gsc) as well as knockdown of E-cadherin (sh-Ecad). Taube et al. reported as list of 159 genes down-regulated under all conditions and 87 up-regulated under all conditions. We found that E-genes are significantly enriched among down-regulated genes (55 downregulated; Fisher's Exact Test, $p$ = 6.6e-52) and M-genes are significantly enriched among up-regulated genes (37 upregulated, Fisher's Exact Test, $p$ = 1.1e-36). However, because of the strict selection criteria, these genes are highly expressed across all condition and there are relatively few of them in each cluster (18, 15 and 4 in M1, M2, and M3 respectively). As such, we were unable to use these upregulated genes alone to distinguish between our M-gene clusters.

Therefore, we reanalyzed the microarray data produced by Taube et al. [1] Data relating to the Taube et al. study was retrieved from the Gene Expression Omnibus, accession numbers GSE9691 and GSE24202. These data were analyzed using the 'limma' and 'affy' packages available through Bioconductor [2]. Normalization was done using RMA and the significance of each differential expression model (treatment – control) was determined using empirical Bayes. We applied the same criteria used by Taube et al. [1] (magnitude of fold-change > 2) to determine differential expression in response to each treatment. Enrichment of E, M, M1, M2, and M3-genes were evaluated using Fisher's Exact test. Significance of enrichment differences between groups was determined using the log of the odds of ratio of enrichment, which is approximately normally distributed with a standard error of:

$$SE = \sqrt{\frac{1}{n_1} + \frac{1}{n_2} + \frac{1}{n_3} + \frac{1}{n_4}}$$

Where the n values are the counts of the contingency table used to determine enrichment.

Using this approach, we found that only E-genes are enriched among down-regulated genes for any factor, while both M-genes as a whole and each cluster of M-genes are enriched among in every set of up-regulated genes (**Supplementary Table 3**). To compare responsiveness across M-gene clusters, we looked at the difference in the log-odds of enrichment **(Supplementary Figure 10)** and found no evidence for differential enrichment for Snail, Twist, or E-cadherin knockdown. For TGF-β, we observe a non-significant reduction in the log-odds of enrichment for M3-genes, which we found to be significantly less responsive to TGF-β in our CAGE data set. However, we do not consider this a contradiction, given both differences methodology and the fact that, while not significant, the confidence interval of the difference of log-odds is biased in favor of reduced response to TGF-β. Conversely, there is evidence of differential response to Gsc between our M-genes clusters. There is a significant difference in the enrichment of Gsc up-regulated genes between M1 and M2 and a large but non-significant difference between M1 and M3, both of which indicate reduced enrichment of M1-genes among Gsc up-regulated genes. As such, it is not surprising that, when we considered the expression of all M-genes, we see a significant difference in the log fold-change of expression in response to Gsc between M1 and M2 genes (Mann-Whitney U-test, $p$ = 0.004) and between M1 and M3 genes (Mann-Whitney U-test, $p$ = 0.019), but not between M2 and M3 genes (Mann-Whitney U-test, $p$ = 0.5237) (**Supplementary Figure 11**). Gsc is notable in that Taube et al. found it was not up-regulated by any of the other EMT inducers used in their study, suggesting that Gsc regulates EMT in an independent fashion. As such, while these results do indicate that M2 and M3 genes may be regulated by additional EMT factors, it is in a way that is independent of the TGF-β/ZEB1 pathway explored in our study.

### Supplementary Text 2 – Gene set enrichment analysis of M-gene clusters

Gene set enrichment analysis was done for the genes in each M-gene clusters against the Molecular Signature Database maintained by the Broad Institute [3]. Specifically, we compare M-gene clusters to selected cancer related expression sets from the 'CGP: chemical and genetic perturbations' database. We also considered the 'CGN: cancer gene neighborhoods', 'CM: cancer modules' and 'C6: oncogenic signatures' sets, but chose not to use them for the following reasons: the CGN and CM database contained mostly computationally clustered gene sets without further annotation while C6 focused response to one specific gene or factor in a cancer context rather differential expression across broad cancer phenotypes. Using the described criteria, we found 18, 16, and 22 sets enriched in M1, M2, and M3-gene clusters respectively (**Supplementary Table 6**) for a total 31 unique enriched sets, the majority (16) of which are enriched in more than one cluster. Of the remaining 15 sets, 10 are unique to M3 genes. The gene sets enriched in M-gene clusters can be used to distinguish between specific subtypes of cancer: for example, M3 genes are enriched amongst those up-regulated in luminal A breast cancer, but down-regulated in luminal B. M3 genes are also enriched in those up-regulated in papillary thyroid carcinoma but down regulated in follicular carcinoma. Nevertheless, we should be careful not to extrapolate the annotation to the whole of any M-gene cluster as the size of the enriched groups vary widely (between 3 and 41 genes, or 4% and 28% of a cluster). Furthermore, some annotations are contradictory, such as both M1 and M3 being enriched for genes that are up-regulated and down-regulated in basal breast cancer. Instead, these results suggest that M-gene clusters contain sub-clusters of genes which are associated with a diversity cancer types, with M3 genes standing out as having both the greatest overall and most unique associations.

**Supplementary Text 3. Expression of M-gene clusters in mesenchymal-like cancer cell lines**

To examine the expression of M-gene clusters in cancer cells, we obtained RNA-sequencing data from the Cancer Cell Line Encyclopedia, which contains transcriptomic information of 1019 cancer cell lines [4]. We selected the cell lines with mesenchymal-like expression profile by calculating the mean M-genes and mean E-genes (the classification of E and M genes is based on our SOM followed by supervised learning. See **Methods**) expressions in all cell lines. Visual inspection of the expression profiles of the cells suggests that there is a large group of cancer cell lines that highly express M-genes but not E-genes (**Supplementary Figure 12A**). We used a simple threshold to select the mesenchymal-like cancer cell lines (mean M-gene $\log_2$(RPKM) > -1, mean E-gene $\log_2$(RPKM) < 1), and we obtained 416 such cell lines. We found that the mean expressions of the three M clusters are significantly different in these cell lines (**Supplementary Figure 12B**, p-values < $10^{-5}$). The mean expression of M2 gene cluster is higher than that of the other two clusters by more than 2-fold, and the mean expression of M3 cluster is higher than that M1 cluster by 30% (**Supplementary Figure 12B**).

Because we showed that the high expression of M3 genes are significantly associated with good prognosis, we asked if these genes are correlated in cancer settings. We calculated the Pearson correlation coefficients of all pairs of M3 genes (within-cluster correlations) in terms of their expression in the 416 cell lines (**Supplementary Figure 12C**, red), and we compared these correlations with those of M3 genes with M1 and M2 genes (between-cluster correlations) (**Supplementary Figure 12C**, gray). We found that the within-cluster correlations are significantly more positive than the between-cluster correlations (0.020 vs. -0.008, $p < 10^{-5}$). However, the within-cluster correlations exhibit a large degree of heterogeneity (**Supplementary Figure 12C**), reflecting the complexity of the regulations on these genes in cancer cells. Nonetheless, when we used a threshold ($r > 0.25$, above the maximal correlation in between-cluster gene pairs) to select genes that are involved in positive correlations. 73 out of 77 M3 genes were involved in the single cluster connected by edges based on the correlations (**Supplementary Figure 13**). Collectively, these results suggest that even though the regulatory circuits of M-gene clusters obtained in our study cannot be extrapolated to gene expressions in cancer cells directly, their differential patterns and within-cluster correlations are observed in cancer cells. Therefore, our study provides useful information in understanding M-genes in cancer cells.

### SUPPLEMENTARY FIGURES

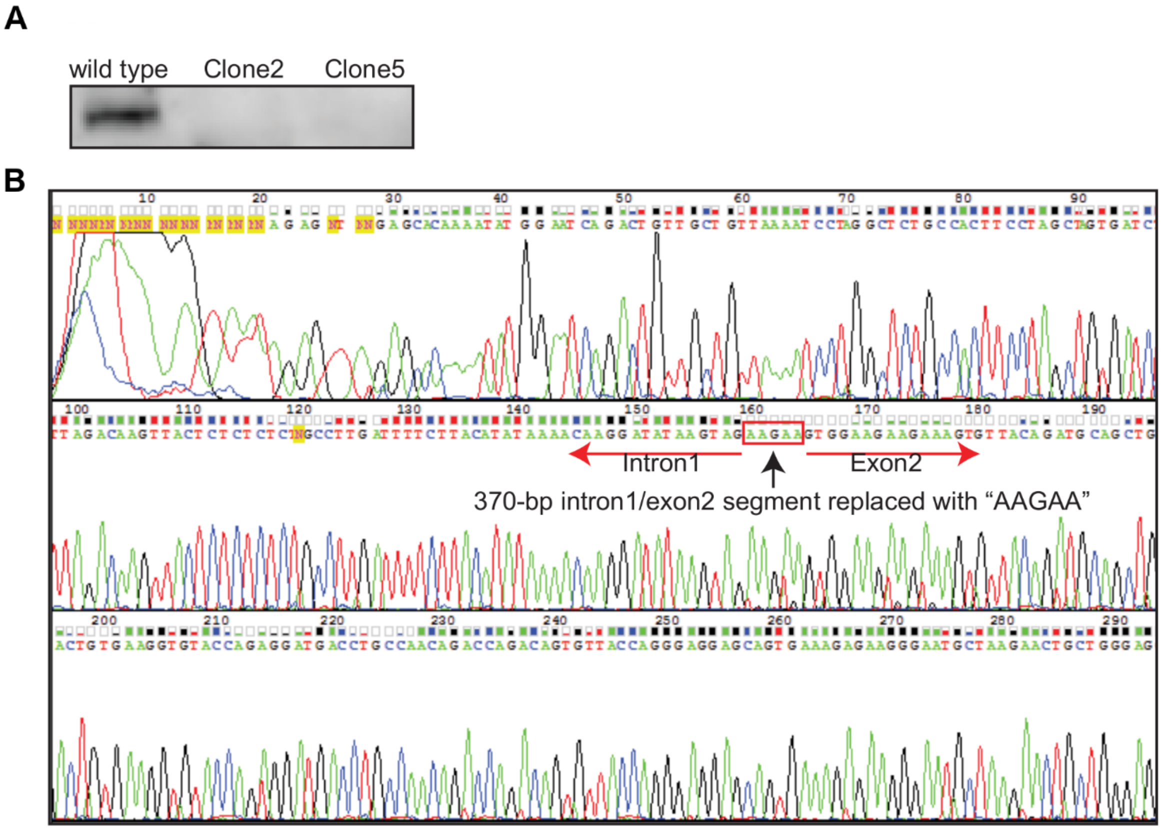

**Supplementary Figure 1. Establishment of ZEB1-deficient MCF10A cells. (A)** Western blotting for ZEB1 protein. Control MCF10A and ZEB1 KO clones were treated with TGF-β to induce endogenous ZEB1. **(B)** Sanger sequencing result at the Intron1-Exon2 boundary of ZEB1 gene in clone5.

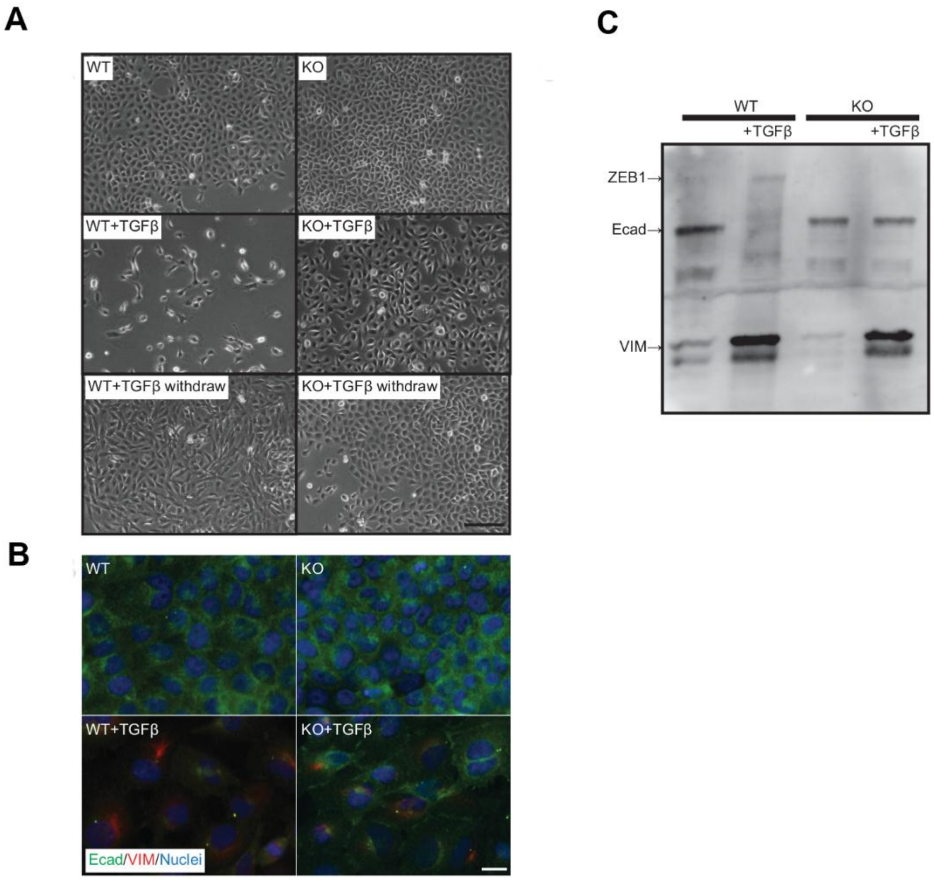

**Supplementary Figure 2. Role of ZEB1 in reciprocity of E and M phenotypes in EMT. (A) Morphological changes induced by TGFβ treatment in wild-type (WT) and ZEB1-deficient (KO) cells.** Cells were treated with 200 pM TGF-β recombinant protein for 2 weeks (+TGF-β) and then cultured without TGF-β for another 2 weeks (+TGF-β withdraw). Scale bar = 100 µm. **(A) Immunofluorescent analysis of E-cadherin (Ecad, green) and vimentin (VIM, red).** Cells were treated with 200 pM TGF-β recombinant protein for 2 weeks (+TGF-β). Nuclei were counterstained with DAPI. Scale bar = 20 µm. **(C) Western blot analysis for epithelial (E) and mesenchymal (M) marker proteins.** Protein samples were collected from cells after TGFβ treatment as described above. The blot was sequentially incubated with antibodies against the indicated proteins (ZEB1, Ecad, and VIM, respectively) and the signals were detected simultaneously.

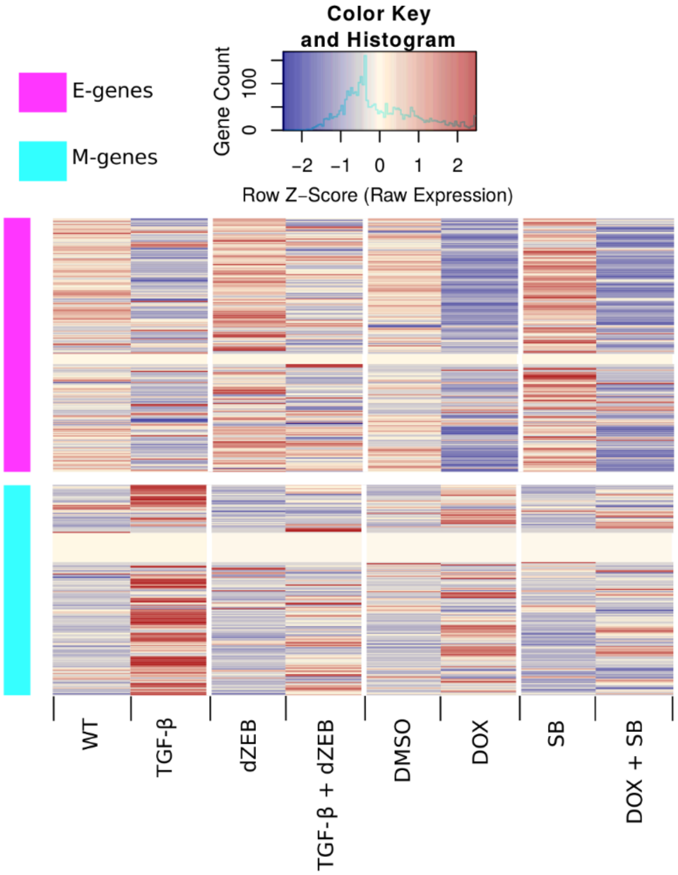

**Supplementary Figure 3. Expression of EMT genes in the 8 combinations of TGF-β and ZEB1 treatments** (see **Table 1**). Expression values were scaled across genes (rows) to adjust for different average expression between genes. Cells in the heatmap are colored based on the z-score of each expression value in its respective row, indicating whether the gene is expressed relatively higher (red) or lower (blue) in that condition than on average. E-genes (pink) and M-genes (light blue) were manually separated, then ordered using hierarchical clustering.

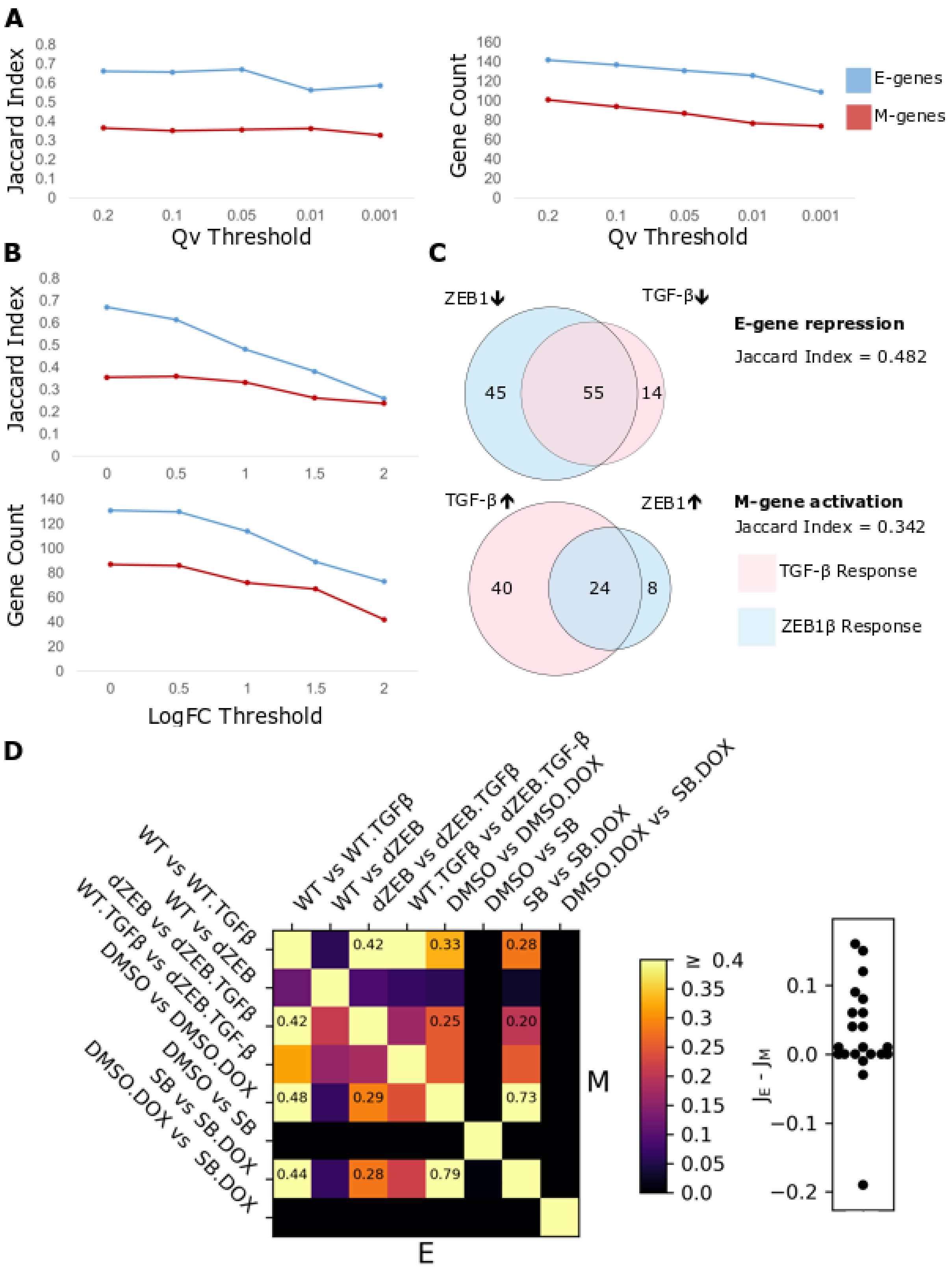
A
Jaccard Index
Qv Threshold
Gene Count
Qv Threshold
E-genes
M-genes
B
Jaccard Index
Gene Count
LogFC Threshold
C
ZEB1
TGF-β
E-gene repression
Jaccard Index = 0.482
45
55
14
TGF-β
ZEB1
M-gene activation
Jaccard Index = 0.342
40
24
8
TGF-β Response
ZEB1β Response
D
WT vs WT.TGFβ
WT vs dZEB
dZEB vs dZEB.TGFβ
WT.TGFβ vs dZEB.TGF-β
DMSO vs DMSO.DOX
DMSO vs SB
SB vs SB.DOX
DMSO.DOX vs SB.DOX
M
E
≥ 0.4
$J_E$ - $J_M$

**Supplementary Figure 4. Quantification of EMT gene expression in response to TGF-β and ZEB1 under different definitions of differential expression. (A)** (Left) Jaccard index for response to TGF-β and ZEB1 using different q-value thresholds and (Right) the total number of genes passing the threshold for E-(blue) and M-(red) genes. **(B)** (Top) Jaccard index for response to TGF-β and ZEB1 using different log fold-change thresholds with q-value cutoff of 0.05 and (Bottom) the total number of genes passing both thresholds for E-(blue) and M-(red) genes. **(C)** Venn diagrams showing the overlap in E- (top) and M-(bottom) genes which show significant expression differences in response to TGF-β or ZEB1 treatment relative to their respective control conditions and have an absolute log fold-change >1. TGF-β response is in pink while ZEB1 response is in light blue. The number E- and M-genes responding to TGF-β or ZEB1 uniquely as well as those responding to both are listed in the respective part of the Venn diagram. The type of response, activation or repression, is indicated by directional arrows (up-arrow: activation, down-arrow: repression). The overlap was quantified using the Jaccard index, which is the number of genes differentially expressed genes by both TGF-β and ZEB1 divided by the total number of differentially expressed gene. **(D)** Jaccard indices of E- and M-genes for each pair of conditions where only genes with an absolute log fold-change >1 are considered. Heatmap shows all the Jaccard indices. Lower triangular entries: E-genes. Upper triangular entries: M genes. The Jaccard index those cells labeled in Figure 1D are also labeled here for comparison. Swarm plot shows the differences between the Jaccard indices of E-genes and those of the M-genes for each of the 28 pairs of conditions ($p < 0.001$ for single value t-test with a null distribution centered at 0).

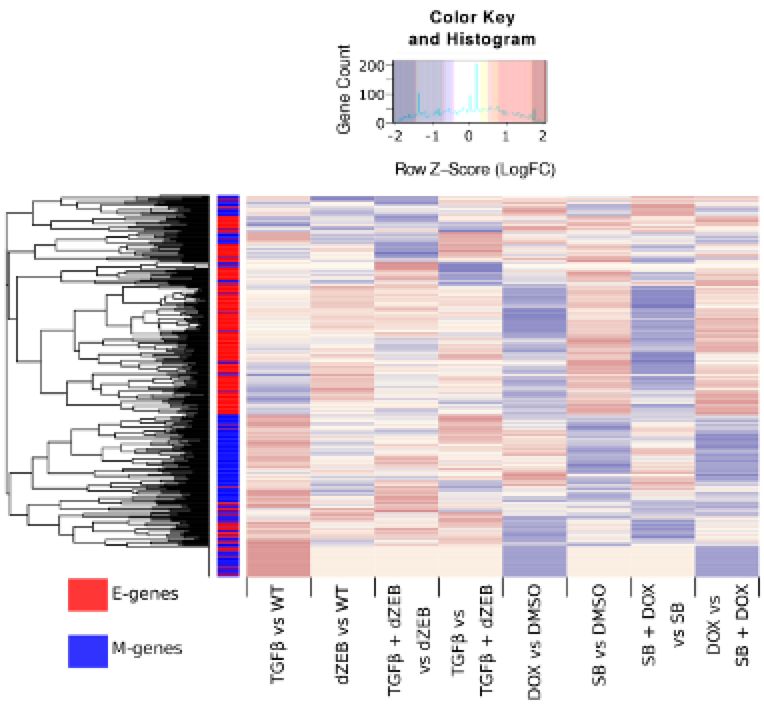

**Supplementary Figure 5**. **Clustering E- and M-genes according to log fold-change of expression.** Log fold-change of expression of EMT genes in the 8 contrasts conditions (see **Table 2**). Expression values were scaled across genes (rows) to adjust for different average expression between genes. Cells in the heatmap are colored based on the z-score of each expression value in its respective row, indicating whether the gene is expressed relatively higher (yellow) or lower (orange) in that condition than on average. The order of genes was determined by hierarchical clustering without accounting for annotation. E (red) or M (blue) gene annotation is indicated by the color of the left side bar.

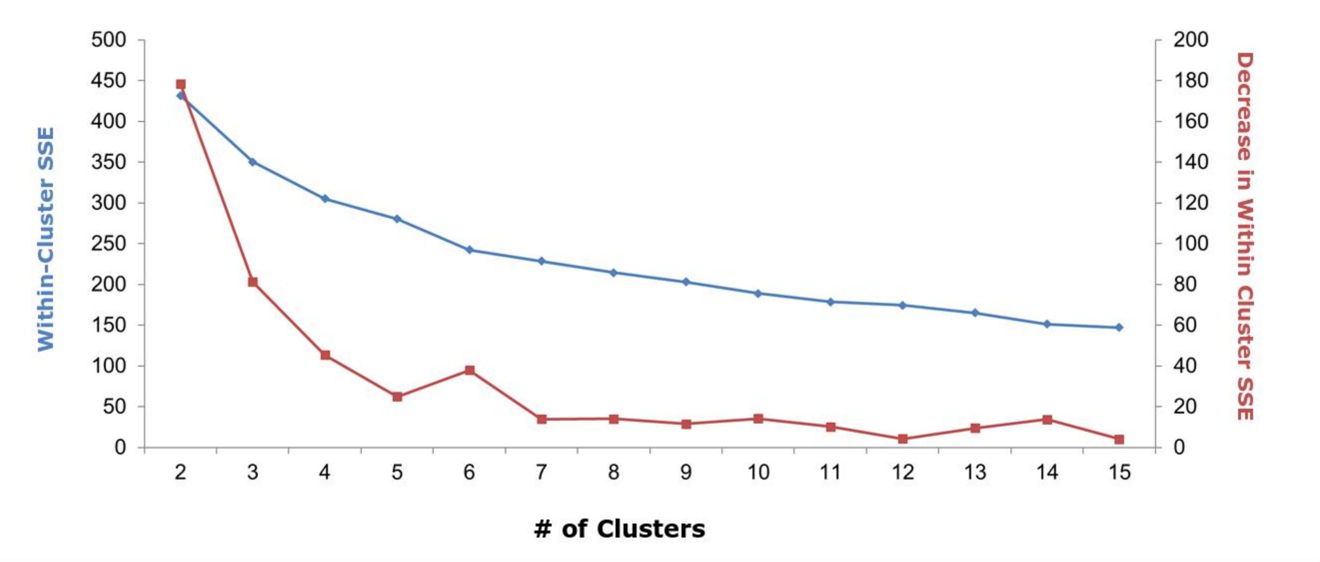

**Supplementary Figure 6**. **Elbow plot showing the changes of sum of squared errors as the number of clusters increases.** Hierarchical clustering was performed on the average log fold-change of expression of each of the 38 M-gene nodes. Within-Cluster standard squared errors were calculated with different choices of number of clusters for cutting the phylogenetic tree. Blue curve: sum of squared errors. Red curve: changes of Within-Cluster standard squared errors when the number of clusters changes from n-1 to n.

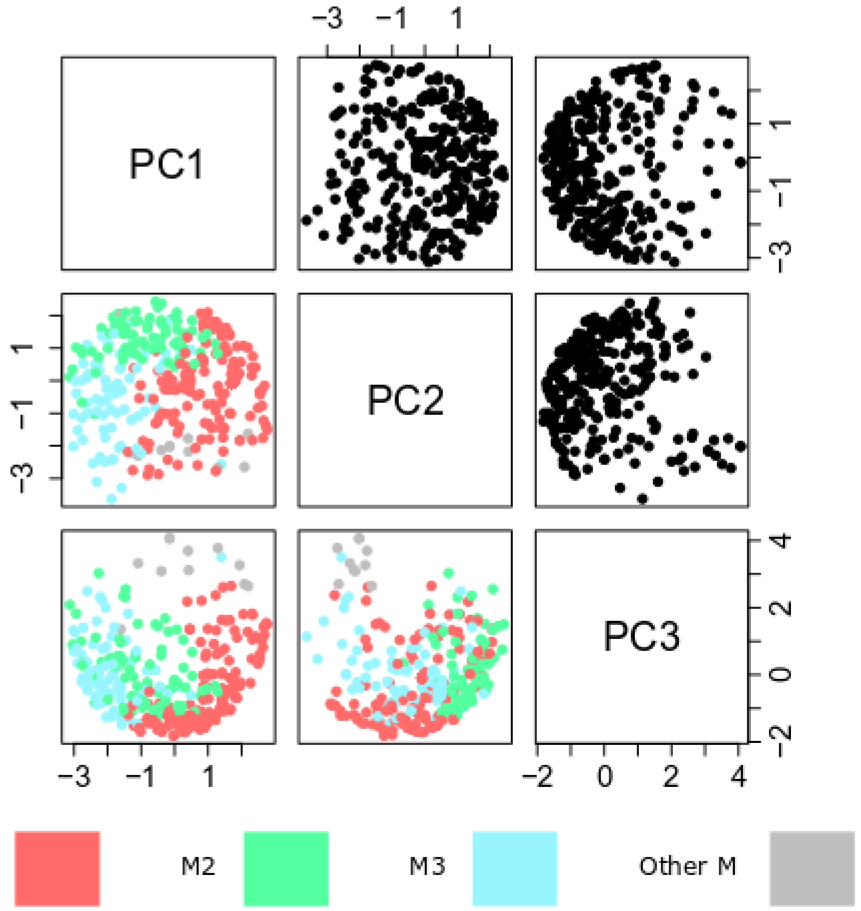

**Supplementary Figure 7. Principal Component Analysis of CAGE expression data for M-genes**. Lower triangle: M-genes plotted against pairs of the first three principal components of the normalized CAGE expressed data set labeled according to their cluster (M1 = red, M2 = green, M3 = blue, Other M = grey). Upper triangle: the same data points as those in the lower triangle without the cluster labels.

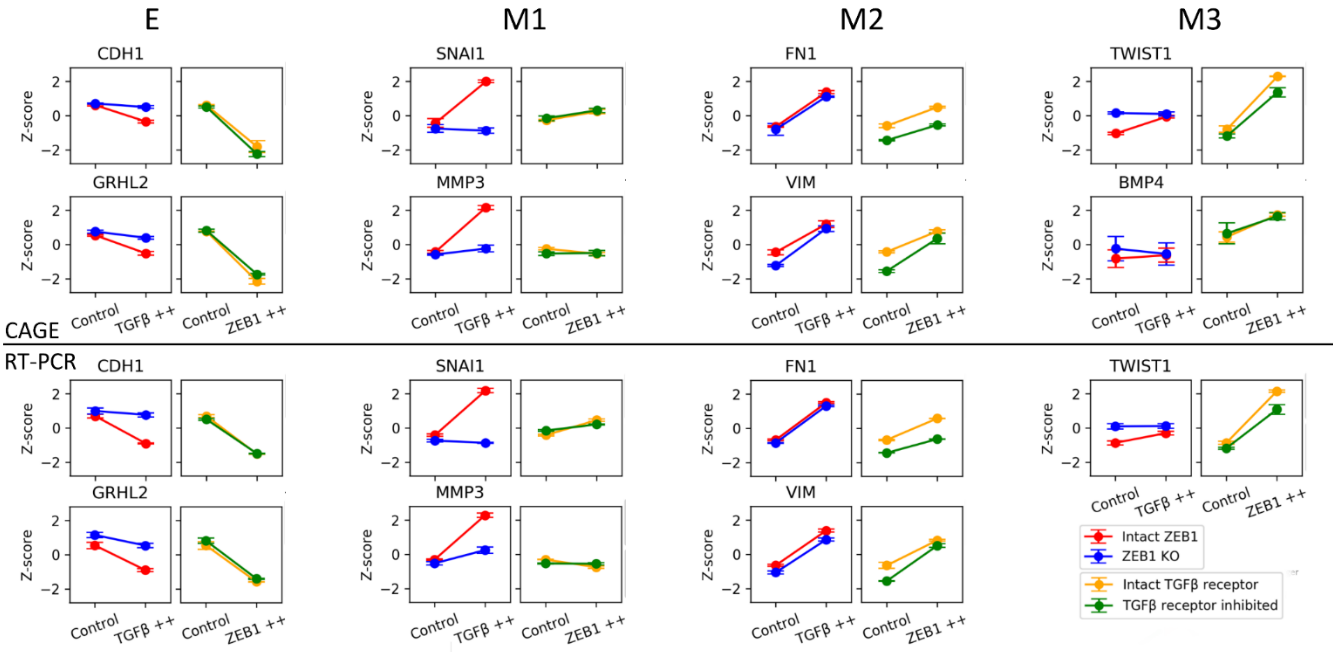

**Supplementary Figure 8. Comparison of expression patterns representative genes in four clusters using CAGE and RT-PCR.** Expression values of representative genes from each cluster were extracted from the CAGE data set. RT-PCR was performed to verify their expression. 2 replicates were used for each gene at each condition. Error bars indicate the standard error of the means. Z-scores were calculated by centering each of the eight data points to 0 and scale them to unit variance.

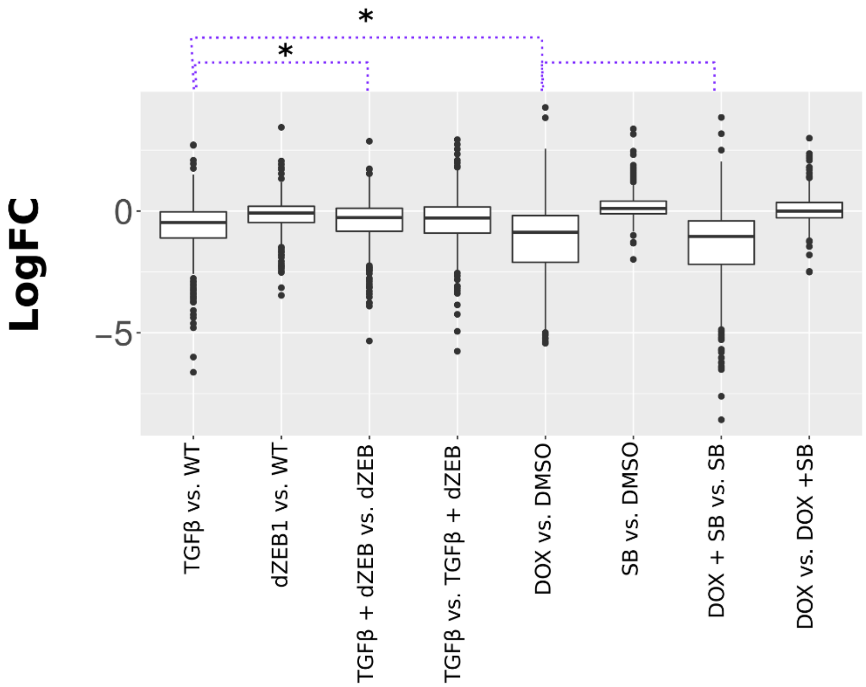

**Supplementary Figure 9. Expression of E-gene cluster under TGF-β and ZEB1 regulation.** Boxplot of Log-Fold expression of E-genes annotated by SOM clustering in the 8 combination of TGF-β and ZEB1 treatment. The middle region indicates the inter-quartile range of expression while whiskers extend 1.5 times this range on either side. Outliers are indicated by black dots. The purple dotted lines above the plot indicate comparisons between the expression under conditions, specifically, TGF-β vs. WT to DOX vs. DMSO, TGF-β vs. WT to TGF-β + dZEB vs. dZEB and DOX vs. DMSO to DOX + SB vs. SB. A * indicates that distribution of expression is significantly different based on the Mann-Whitney U-test. Note that E-genes have a similar response patter to M3 genes (**Figure 3C)** but the direction of response is reversed (repression in E-genes vs. activation in M3 genes).

| Data | Comparison | Δ Log Odds | P-value |
|---|---|---|---|
| TGF-β | M1 vs M2 | -0.458 | 0.161 |
| TGF-β | M1 vs M3 | 0.342 | 0.368 |
| TGF-β | M2 vs M3 | 0.800 | **0.053** |
| Twist | M1 vs M2 | -0.259 | 0.405 |
| Twist | M1 vs M3 | -0.158 | 0.622 |
| Twist | M2 vs M3 | 0.100 | 0.782 |
| Snail | M1 vs M2 | -0.400 | 0.186 |
| Snail | M1 vs M3 | 0.016 | 0.960 |
| Snail | M2 vs M3 | 0.416 | 0.251 |
| Gsc | M1 vs M2 | -1.114 | **0.0002** |
| Gsc | M1 vs M3 | -0.603 | **0.051** |
| Gsc | M2 vs M3 | 0.510 | 0.134 |
| sh-Ecad | M1 vs M2 | -0.260 | 0.411 |
| sh-Ecad | M1 vs M3 | -0.003 | 0.992 |
| sh-Ecad | M2 vs M3 | 0.256 | 0.0495 |

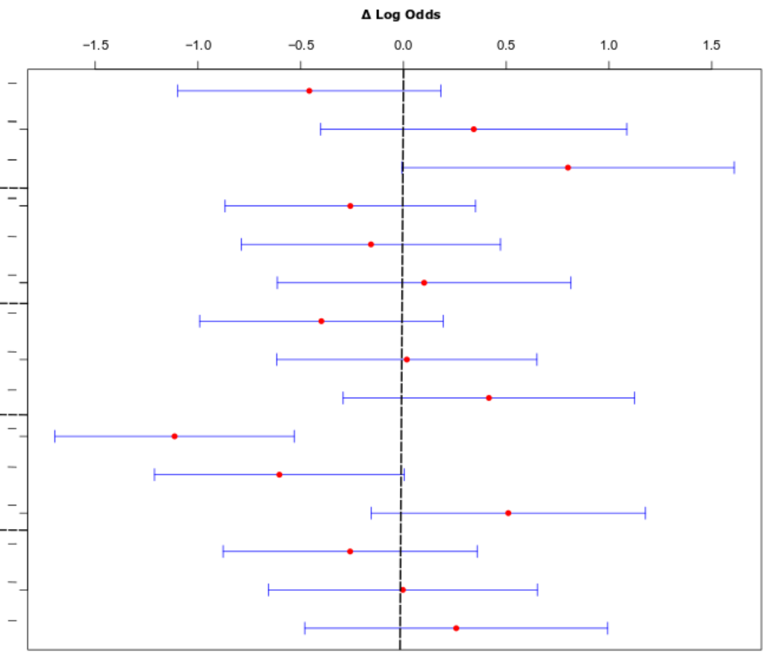

**Supplementary Figure 10. Significance and confidence intervals for difference in odds of enrichment between M-gene groups across different regulatory data sets**. Left: A table of values showing the significance of enrichment of M1, M2, and M3 genes among genes differentially expressed during the overexpression of TGF-β, Twist, Snail, Gsc or the knockdown of E-cadherin (sh-Ecad). For each comparison between M-gene groups, mean difference in odds is given followed by the two-tailed p-value. Right: 95% confidence intervals of each comparison, where the red dot indicates the average value and blue whiskers indicate the interval range. The dotted line is located at zero which represent no difference in the log of the odds ratio of enrichment between the compared M-gene clusters.

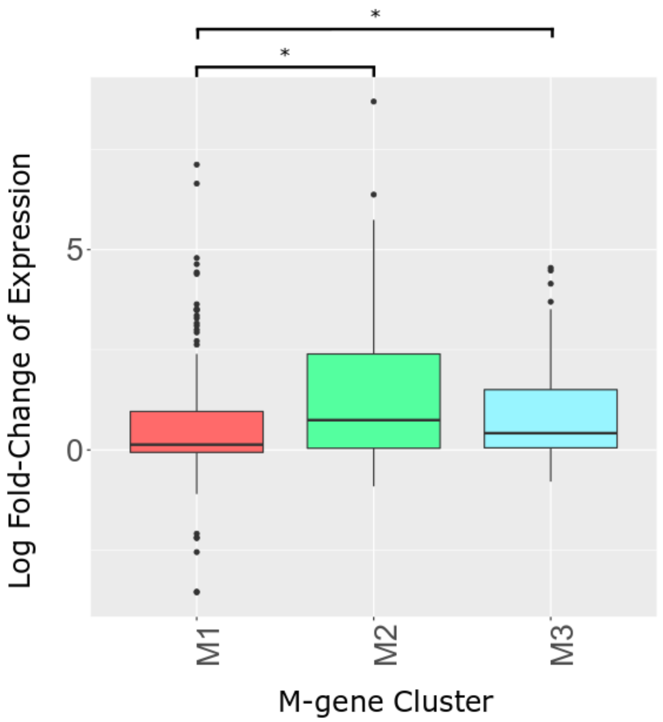

**Supplementary Figure 11. Log fold-change of expression of each M-gene cluster in response to GSC overexpression**. Boxplot of log fold-change of expression of M-genes cluster in response to the overexpression of Gsc in microarray data from Taube et al. [1]. The colored region (M1 = red, M2 = green, M3 = blue) indicates the inter-quartile range of expression while whiskers extend 1.5 times this range on either side. Outliers are indicated by black dots. The lines above the plot indicate comparisons between the expression under conditions, specifically, M1 vs M2 and M1 and M3. A * indicates that distribution of expression is significantly different based on the Mann-Whitney U-test at an alpha of 0.05.

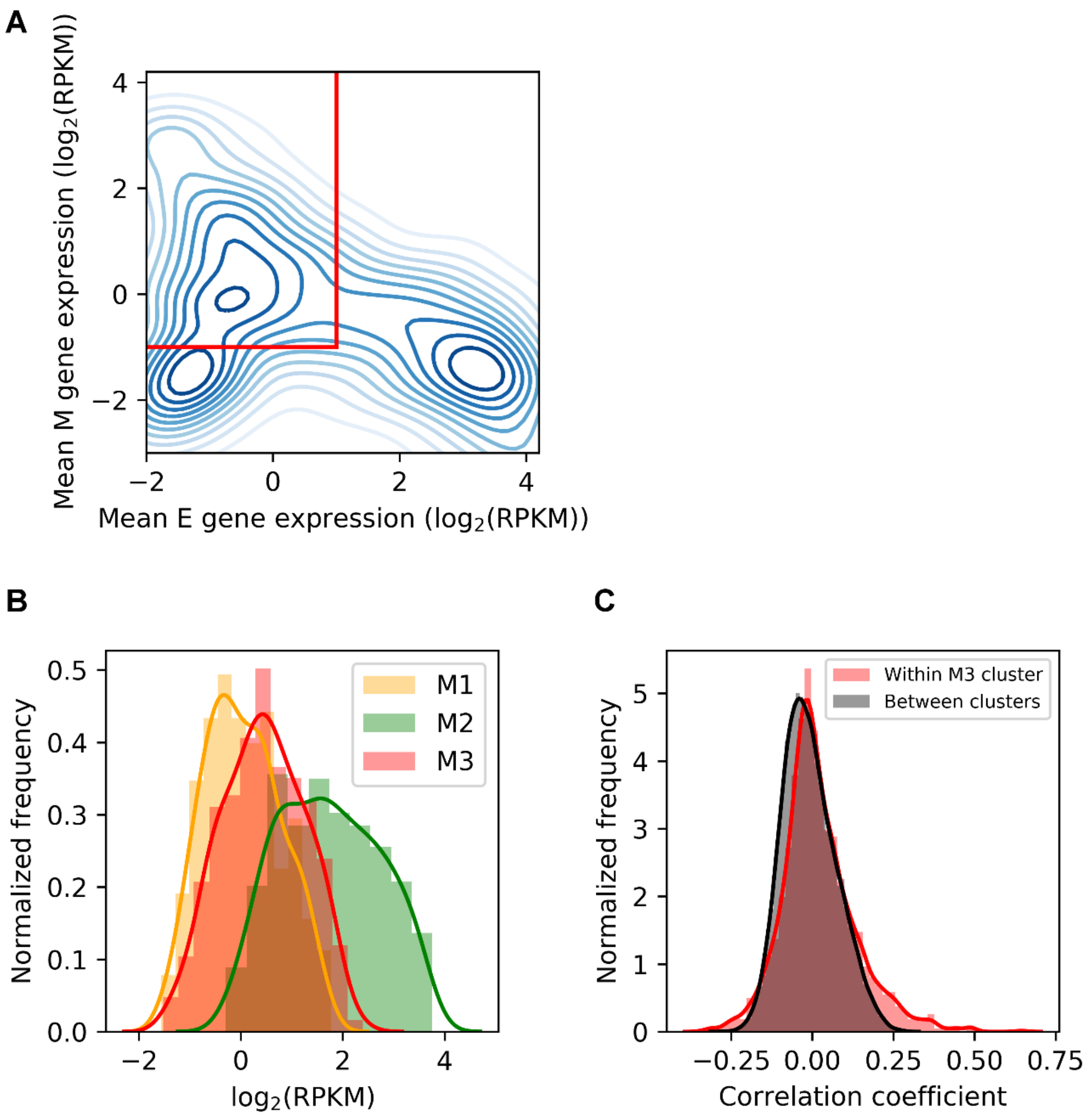

**Supplementary Figure 12. Expression of three cluster of M-genes in cancer cell lines. A.** Mean expressions of E-genes and M-genes in 1019 cancer cell lines [4]. Red lines indicate the thresholds for selecting mesenchymal-like cancer cells (mean M-gene expression $log_2$(RPKM) > -1, mean E-gene expression $log_2$(RPKM) < 1). Contour lines indicate densities of cells. **B.** Mean expressions of M-gene clusters in 416 mesenchymal-like cancer cells. **C.** Pearson correlations between pairs of M3 genes (red), and those between M3 genes and M1/M2 genes (gray). Histograms show the frequencies. Curves show kernel density estimates.

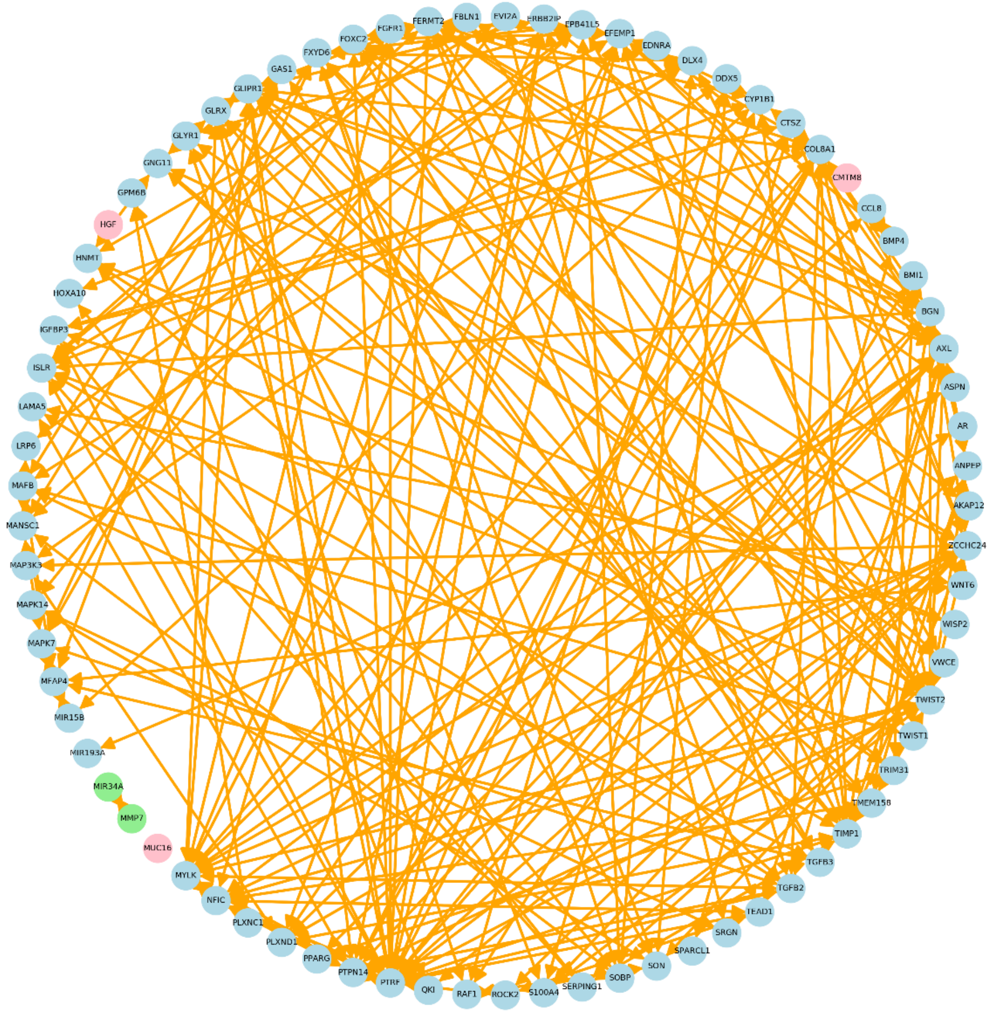

**Supplementary Figure 13. Correlation network of M3 genes.** Correlation based network of M3 genes obtained from the analysis of gene expression in 416 mesenchymal-like cancer cells (**Supplementary Text 3**). Correlation threshold (r > 0.25) was chosen based on distributions of correlation coefficients of within-cluster correlations and between-cluster correlations (**Supplementary Figure 13C**). The threshold is above the maximal correlation coefficient of the between-cluster correlations. Orange lines denote the association between genes using this threshold. Blue icons show 73 M3 genes forming a large cluster based on these associations. Green and pink icons show 4 M3 genes that are not in the cluster.

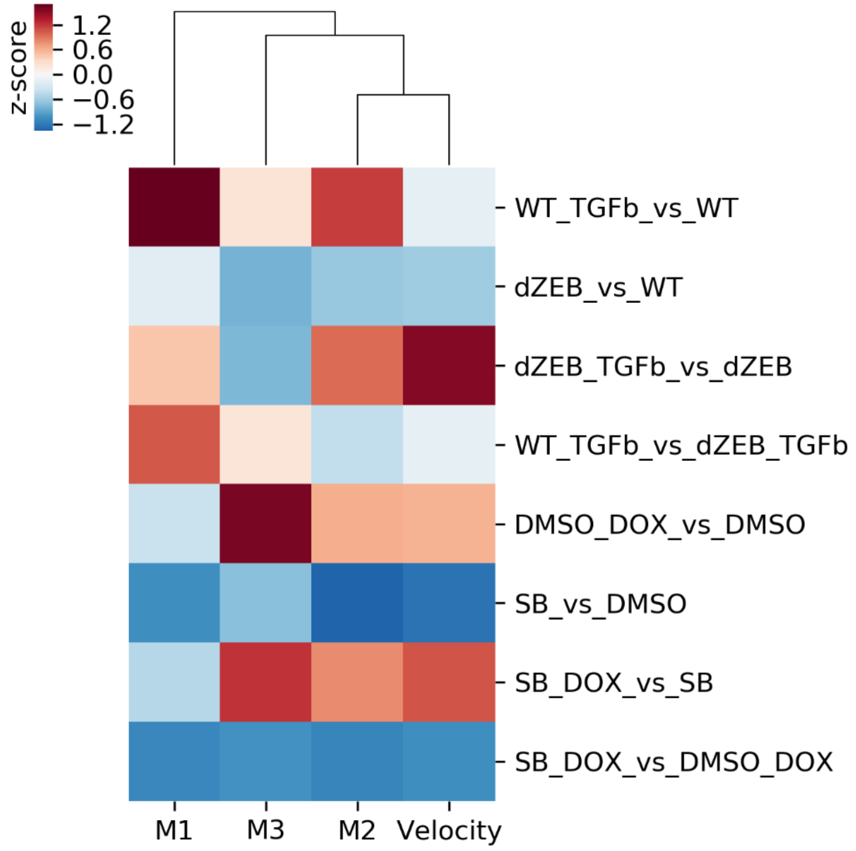

**Supplementary Figure 14. Correlation between changes of mean gene expression and velocity of cells under 8 contrast conditions.** The fold-changes of mean gene expression of three M-gene clusters under 8 contrast experimental conditions were clustered together with that of the velocity of cells under the same conditions. Column z-scores were computed to generate the heatmap

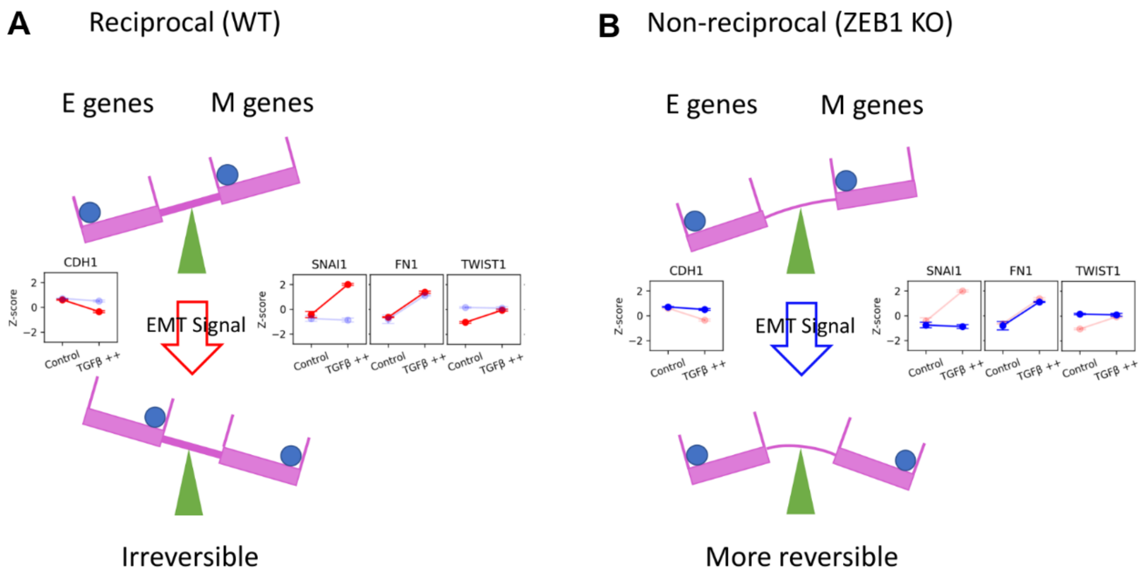

**Supplementary Figure 15. Metaphoric illustration of relationship between reciprocity of E- and M-genes and irreversibility of EMT transition. (A)** Under normal condition, E- and M-genes are reciprocally regulated and both balls representing the feedback loops involving E- and M-genes move upon the external force followed by the flip of the seesaw, which makes the EMT irreversible **(B)** Under ZEB1 knockout condition, reciprocity of E- and M-genes is compromised and some of the feedback loops are no longer functioning. EMT becomes less irreversible in this scenario.

**SUPPLEMENTARY TABLES**

**Supplementary Table 1.** Overlapping EMT gene expression before and after annotating dbEMT genes

| Overlap Condition | Class | Original Annotation | Updated Annotation |
|---|---|---|---|
| TGF-β and ZEB1 | E | 67.1% | 55.5% |
| | M | 35.6% | 30.1% |
| TGF-β and TGF-β w/o ZEB1 | E | 63.3% | 46.9% |
| | M | 46.1% | 47.2% |
| ZEB1 and ZEB1 w/o TGF-β | E | 93.6% | 77.3% |
| | M | 67.5% | 60.0% |

**Supplementary Table 2.** List of EMT genes analyzed in this study and their clustering information

| Gene | Cluster | Annotation | Source | P_values | LEC_months | HEC_months |
|---|---|---|---|---|---|---|
| ABCC3 | E | E | Tan et al., 2014 | 0.6026 | 228.85 | 216.66 |
| ABHD11 | E | E | Tan et al., 2014 | 0.0013 | 228.85 | 216.66 |
| ADAM17 | E | NA | dbEMT | 0.032 | 56.6 | 42 |
| ADAP1 | E | E | Tan et al., 2014 | 5.20E-08 | 39 | 64 |
| AGER | E | NA | dbEMT | 0.0002 | 42 | 59 |
| AGR2 | E | E | Tan et al., 2014 | 0.0171 | 29 | 40.3 |
| AIM1 | E | E | Tan et al., 2014 | 0.0301 | 55.43 | 42 |
| AKR1B10 | E | E | Tan et al., 2014 | 0.3266 | 228.85 | 191.21 |
| ALDH3B2 | E | E | Tan et al., 2014 | 0.0357 | 216.66 | 191.21 |
| ALOX5 | E | E | Tan et al., 2014 | 1.80E-06 | 184.04 | 216.66 |
| AMHR2 | E | NA | dbEMT | 1.60E-09 | 38 | 66 |
| ANK2 | E | M | Tan et al., 2014 | 2.90E-12 | 35.04 | 69.7 |
| ANK3 | E | E | Tan et al., 2014 | 1.60E-07 | 28.96 | 46.8 |
| ANXA1 | E | NA | dbEMT | 0.3191 | 53 | 45.6 |
| ANXA4 | E | E | Tan et al., 2014 | 0.6625 | 47.51 | 48 |
| ANXA9 | E | E | Tan et al., 2014 | 3.60E-05 | 228.85 | 216.66 |
| AP1M2 | E | E | Tan et al., 2014 | 0.2305 | 228.85 | 216.66 |
| AQP3 | E | E | Tan et al., 2014 | 0.0001 | 58 | 41.69 |
| ARAP2 | E | E | Tan et al., 2014 | 0.6035 | 50.4 | 46 |
| AREG | E | E | Tan et al., 2014 | 0.0024 | 42.3 | 56.4 |
| ARHGAP8 | E | E | Tan et al., 2014 | 8.90E-07 | 40.08 | 62 |
| ARHGDIB | E | E | Tan et al., 2014 | 5.40E-07 | 38.4 | 61 |
| ARHGEF5 | E | E | Tan et al., 2014 | 0.3123 | 228.85 | 216.66 |
| ATP2C2 | E | E | Tan et al., 2014 | 0.0659 | 52.83 | 44 |
| AURKA | E | NA | dbEMT | 1.00E-16 | 216.66 | 163.46 |
| AXIN1 | E | NA | dbEMT | 5.60E-06 | 191.21 | 216.66 |
| AZGP1 | E | E | Tan et al., 2014 | 7.10E-06 | 40.56 | 61 |
| B3GNT3 | E | E | Tan et al., 2014 | 0.6972 | 185.16 | 216.66 |
| BCAS1 | E | E | Tan et al., 2014 | 0.0125 | 216.66 | 228.85 |
| BCL2 | E | NA | dbEMT | 9.90E-11 | 191.21 | 216.66 |
| BCL2L1 | E | NA | dbEMT | 1.90E-05 | 228.85 | 216.66 |
| BIK | E | E | Tan et al., 2014 | 0.4743 | 48 | 48 |
| BLNK | E | E | Tan et al., 2014 | 0.0001 | 228.85 | 216.66 |
| BMP7 | E | NA | dbEMT | 0.0232 | 228.85 | 216.66 |
| BOP1 | E | NA | dbEMT | 0.0002 | 57.27 | 41.64 |
| BSPRY | E | E | Tan et al., 2014 | 6.90E-08 | 171.43 | 216.66 |
| C1orf106 | E | E | Tan et al., 2014 | 4.70E-11 | 228.85 | 184.04 |
| C1orf116 | E | E | Tan et al., 2014 | 0.0006 | 216.66 | 185.16 |
| C1orf54 | E | M | Tan et al., 2014 | 0.5834 | 216.66 | 228.85 |
| C4orf19 | E | E | Tan et al., 2014 | 0.126 | 38 | 33 |
| CAMK2N1 | E | E | Tan et al., 2014 | 0.0044 | 216.66 | 191.21 |
| CAPN1 | E | E | Tan et al., 2014 | 0.2362 | 228.85 | 216.66 |
| CAV1 | E | M | Tan et al., 2014 | 0.0117 | 216.66 | 228.85 |
| CBLC | E | E | Tan et al., 2014 | 0.2914 | 228.85 | 216.66 |

| | | | | | | |
|---|---|---|---|---|---|---|
| CBR1 | E | NA | dbEMT | 0.0593 | 53.04 | 44.4 |
| CD2AP | E | E | Tan et al., 2014 | 4.20E-06 | 216.66 | 191.21 |
| CD46 | E | E | Tan et al., 2014 | 0.0103 | 55.66 | 45 |
| CD9 | E | E | Tan et al., 2014 | 0.0101 | 216.66 | 228.85 |
| CDH1 | E | E | Tan et al., 2014 | 2.80E-05 | 59 | 40.56 |
| CDH3 | E | E | Tan et al., 2014 | 0.0108 | 216.66 | 228.85 |
| CDS1 | E | E | Tan et al., 2014 | 0.4644 | 48 | 48 |
| CEACAM1 | E | E | Tan et al., 2014 | 0.0119 | 228.85 | 191.21 |
| CEACAM5 | E | E | Tan et al., 2014 | 0.691 | 228.85 | 216.66 |
| CEACAM6 | E | E | Tan et al., 2014 | 0.0234 | 228.85 | 184.04 |
| CLDN1 | E | NA | dbEMT | 0.3356 | 39 | 30.72 |
| CLDN4 | E | E | Tan et al., 2014 | 0.1675 | 228.85 | 216.66 |
| CLDN7 | E | E | Tan et al., 2014 | 0.1046 | 51 | 46 |
| CLEC2B | E | M | Tan et al., 2014 | 1.40E-05 | 28.35 | 44 |
| CLU | E | NA | dbEMT | 1.20E-11 | 37.2 | 69 |
| CNKSR1 | E | E | Tan et al., 2014 | 2.20E-15 | 150.54 | 216.66 |
| COL14A1 | E | M | Tan et al., 2014 | 1.00E-06 | 28 | 46 |
| COMT | E | E | Tan et al., 2014 | 0.1553 | 228.85 | 216.66 |
| CORO2A | E | E | Tan et al., 2014 | 0.174 | 31.56 | 39 |
| CRISPLD2 | E | M | Tan et al., 2014 | 8.90E-05 | 185.16 | 216.66 |
| CSF2RB | E | M | Tan et al., 2014 | 2.20E-09 | 39 | 65.04 |
| CSK | E | NA | dbEMT | 0.2892 | 48 | 48 |
| CSRP2 | E | M | Tan et al., 2014 | 7.60E-07 | 216.66 | 228.85 |
| CTNNB1 | E | NA | dbEMT | 0.038 | 46 | 53.16 |
| CTNNBIP1 | E | NA | dbEMT | 0.0027 | 43 | 57.3 |
| CTNND1 | E | NA | dbEMT | 0.0494 | 44.22 | 53.1 |
| CTSH | E | E | Tan et al., 2014 | 0.0377 | 43 | 55.13 |
| CXADR | E | E | Tan et al., 2014 | 0.0011 | 42 | 28.35 |
| CXCL16 | E | NA | dbEMT | 0.0052 | 34 | 38.4 |
| CYP4F3 | E | E | Tan et al., 2014 | 4.60E-05 | 39.95 | 60 |
| DAPK1 | E | NA | dbEMT | 0.1356 | 48 | 48 |
| DDR1 | E | E | Tan et al., 2014 | 0.0529 | 216.66 | 228.85 |
| DENND2D | E | E | Tan et al., 2014 | 1.20E-14 | 46 | 53.16 |
| DHCR24 | E | E | Tan et al., 2014 | 4.00E-06 | 43 | 57.3 |
| DPYSL3 | E | M | Tan et al., 2014 | 0.0031 | 44.22 | 53.1 |
| DSC2 | E | E | Tan et al., 2014 | 0.0002 | 43 | 55.13 |
| DSG2 | E | E | Tan et al., 2014 | 5.50E-07 | 42 | 28.35 |
| DSP | E | E | Tan et al., 2014 | 0.4936 | 34 | 38.4 |
| DTX4 | E | E | Tan et al., 2014 | 0.42 | 39.95 | 60 |
| ECT2 | E | NA | dbEMT | 1.00E-16 | 53.16 | 44.22 |
| EGF | E | NA | dbEMT | 0.0003 | 44.17 | 53.56 |
| EGFR | E | NA | dbEMT | 0.0736 | 171.43 | 216.66 |
| EGR1 | E | NA | dbEMT | 2.90E-05 | 40 | 60.12 |
| EHF | E | E | Tan et al., 2014 | 0.3975 | 228.85 | 216.66 |
| EIF3I | E | NA | dbEMT | 0.0148 | 43 | 28.8 |
| EIF5A2 | E | NA | dbEMT | 0.0127 | 61.2 | 38.7 |
| ELF3 | E | E | Tan et al., 2014 | 0.0773 | 228.85 | 216.66 |
| ELL3 | E | NA | dbEMT | 1.50E-05 | 48 | 48 |
| ELMO3 | E | E | Tan et al., 2014 | 0.3213 | 216.66 | 171.43 |
| EPAS1 | E | NA | dbEMT | 0.0469 | 191.21 | 228.85 |

| | | | | | | |
|---|---|---|---|---|---|---|
| EPB41L4B | E | E | Tan et al., 2014 | 0.0232 | 35 | 37 |
| EPCAM | E | E | Tan et al., 2014 | 1.00E-16 | 185.16 | 216.66 |
| EPHA1 | E | E | Tan et al., 2014 | 7.80E-10 | 38.4 | 32.4 |
| EPN3 | E | E | Tan et al., 2014 | 0.0167 | 191.21 | 228.85 |
| EPS8L1 | E | E | Tan et al., 2014 | 0.0003 | 31.56 | 40 |
| EPS8L2 | E | E | Tan et al., 2014 | 0.0389 | 34 | 38.4 |
| ERBB2 | E | E | Tan et al., 2014 | 0.1804 | 62 | 40 |
| ERBB3 | E | E | Tan et al., 2014 | 0.0011 | 228.85 | 216.66 |
| ERMP1 | E | E | Tan et al., 2014 | 0.6614 | 44.22 | 55 |
| ESR1 | E | NA | dbEMT | 2.50E-13 | 30.72 | 40.44 |
| ESRP1 | E | E | Tan et al., 2014 | 4.40E-12 | 77.23 | 33.63 |
| ESRP2 | E | E | Tan et al., 2014 | 0.0006 | 191.21 | 216.66 |
| ETV1 | E | M | Tan et al., 2014 | 8.10E-09 | 216.66 | 185.16 |
| ETV4 | E | NA | dbEMT | 0.8505 | 228.85 | 216.66 |
| EVPL | E | E | Tan et al., 2014 | 0.1597 | 228.85 | 216.66 |
| EXPH5 | E | E | Tan et al., 2014 | 0.0065 | 228.85 | 216.66 |
| EZH2 | E | NA | dbEMT | 4.70E-13 | 27 | 43.93 |
| EZR | E | E | Tan et al., 2014 | 0.0081 | 36 | 36.9 |
| F11R | E | E | Tan et al., 2014 | 0.6525 | 184.04 | 216.66 |
| FA2H | E | E | Tan et al., 2014 | 0.0091 | 51.36 | 27.4 |
| FGF1 | E | NA | dbEMT | 4.70E-05 | 57 | 41.95 |
| FGFR3 | E | E | Tan et al., 2014 | 0.0077 | 228.85 | 216.66 |
| FHL2 | E | NA | dbEMT | 0.0004 | 228.85 | 216.66 |
| FLI1 | E | M | Tan et al., 2014 | 0.5315 | 228.85 | 216.66 |
| FLT1 | E | NA | dbEMT | 0.5276 | 228.85 | 216.66 |
| FOSL1 | E | NA | dbEMT | 0.6232 | 74 | 34.8 |
| FOXA1 | E | E | Tan et al., 2014 | 3.10E-08 | 55.99 | 44 |
| FOXM1 | E | NA | dbEMT | 1.00E-16 | 50 | 46.8 |
| FOXQ1 | E | NA | dbEMT | 0.657 | 55 | 43 |
| FSCN1 | E | NA | dbEMT | 0.0808 | 42 | 59 |
| FUT2 | E | E | Tan et al., 2014 | 2.40E-05 | 42 | 55.2 |
| FUT3 | E | E | Tan et al., 2014 | 0.1216 | 41.95 | 60 |
| FXYD3 | E | E | Tan et al., 2014 | 6.30E-06 | 228.85 | 191.21 |
| GALE | E | E | Tan et al., 2014 | 0.022 | 37.64 | 34.82 |
| GALNT3 | E | E | Tan et al., 2014 | 2.20E-12 | 191.21 | 216.66 |
| GALNT7 | E | E | Tan et al., 2014 | 0.0948 | 191.21 | 216.66 |
| GATA3 | E | NA | dbEMT | 1.00E-05 | 37.64 | 60 |
| GDF15 | E | E | Tan et al., 2014 | 0.5517 | 228.85 | 216.66 |
| GEMIN2 | E | NA | dbEMT | 6.00E-08 | 216.66 | 185.16 |
| GIMAP6 | E | M | Tan et al., 2014 | 0.0004 | 216.66 | 228.85 |
| GMDS | E | E | Tan et al., 2014 | 0.3653 | 52.54 | 41.69 |
| GMNN | E | NA | dbEMT | 1.00E-16 | 216.66 | 184.04 |
| GPR56 | E | E | Tan et al., 2014 | 9.80E-14 | 228.85 | 171.43 |
| GPRC5A | E | E | Tan et al., 2014 | 0.0007 | 216.66 | 191.21 |
| GPX2 | E | E | Tan et al., 2014 | 6.50E-05 | 228.85 | 216.66 |
| GRB7 | E | E | Tan et al., 2014 | 0.0069 | 228.85 | 216.66 |
| GRHL1 | E | E | Tan et al., 2014 | 0.882 | 37 | 34.8 |
| GRHL2 | E | E | Tan et al., 2014 | 0.0001 | 216.66 | 184.04 |
| GRHL3 | E | E | Tan et al., 2014 | 0.9686 | 34.8 | 37 |
| GSK3A | E | NA | dbEMT | 2.00E-08 | 191.21 | 216.66 |

| | | | | | | |
|---|---|---|---|---|---|---|
| GSK3B | E | NA | dbEMT | 0.3888 | 36.44 | 35 |
| HBEGF | E | NA | dbEMT | 9.00E-06 | 184.04 | 216.66 |
| HDGF | E | NA | dbEMT | 6.40E-07 | 228.85 | 216.66 |
| HDHD3 | E | E | Tan et al., 2014 | 0.0161 | 228.85 | 216.66 |
| HES1 | E | E | Tan et al., 2014 | 0.0128 | 184.04 | 216.66 |
| HMGB1 | E | NA | dbEMT | 0.8187 | 36.44 | 36 |
| HPGD | E | E | Tan et al., 2014 | 0.0009 | 43 | 57.6 |
| HPSE | E | NA | dbEMT | 1.70E-07 | 28.8 | 46.8 |
| HRAS | E | NA | dbEMT | 0.0409 | 53.28 | 44.22 |
| HS6ST2 | E | NA | dbEMT | 0.5385 | 36.96 | 35 |
| HSP90AA1 | E | NA | dbEMT | 5.90E-15 | 216.66 | 184.04 |
| HSPA4 | E | NA | dbEMT | 4.30E-07 | 216.66 | 185.16 |
| ICA1 | E | E | Tan et al., 2014 | 7.30E-11 | 228.85 | 216.66 |
| ID1 | E | NA | dbEMT | 0.0002 | 216.66 | 228.85 |
| ID2 | E | NA | dbEMT | 0.0757 | 55.13 | 43 |
| IDH1 | E | NA | dbEMT | 2.10E-06 | 61.05 | 39 |
| IDH2 | E | NA | dbEMT | 7.80E-14 | 216.66 | 184.04 |
| IFFO1 | E | M | Tan et al., 2014 | 2.50E-10 | 37.8 | 65.04 |
| IL18 | E | NA | dbEMT | 4.90E-05 | 216.66 | 228.85 |
| IL1B | E | NA | dbEMT | 1.40E-11 | 171.43 | 216.66 |
| IL1RN | E | E | Tan et al., 2014 | 0.0028 | 44 | 54.77 |
| IL20RA | E | E | Tan et al., 2014 | 2.50E-12 | 171.43 | 216.66 |
| IL6R | E | NA | dbEMT | 6.50E-05 | 30 | 43 |
| ILK | E | NA | dbEMT | 8.20E-05 | 228.85 | 216.66 |
| IRF6 | E | E | Tan et al., 2014 | 0.6955 | 228.85 | 216.66 |
| ITGA6 | E | NA | dbEMT | 0.654 | 51 | 46.68 |
| ITGB3 | E | NA | dbEMT | 0.4851 | 228.85 | 191.21 |
| ITGB4 | E | E | Tan et al., 2014 | 0.0022 | 228.85 | 216.66 |
| JAG1 | E | NA | dbEMT | 0.636 | 228.85 | 216.66 |
| JAG2 | E | NA | dbEMT | 2.50E-05 | 216.66 | 228.85 |
| JUP | E | E | Tan et al., 2014 | 0.2519 | 228.85 | 216.66 |
| KCNK1 | E | E | Tan et al., 2014 | 4.80E-08 | 228.85 | 216.66 |
| KL | E | NA | dbEMT | 2.70E-11 | 37 | 66 |
| KLF5 | E | E | Tan et al., 2014 | 8.70E-06 | 228.85 | 216.66 |
| KLF6 | E | NA | dbEMT | 0.7772 | 38 | 31.51 |
| KLF8 | E | M | Tan et al., 2014 | 0.0079 | 58 | 41.04 |
| KLK6 | E | E | Tan et al., 2014 | 0.0628 | 228.85 | 185.16 |
| KRAS | E | NA | dbEMT | 0.0003 | 191.21 | 228.85 |
| KRT15 | E | E | Tan et al., 2014 | 0.0044 | 45 | 55.9 |
| KRT18 | E | E | Tan et al., 2014 | 0.7277 | 46.8 | 50.3 |
| KRT19 | E | E | Tan et al., 2014 | 0.1144 | 48 | 48.16 |
| KRT7 | E | E | Tan et al., 2014 | 0.0029 | 228.85 | 216.66 |
| L1CAM | E | NA | dbEMT | 0.6474 | 191.21 | 216.66 |
| LAD1 | E | E | Tan et al., 2014 | 0.0237 | 216.66 | 184.04 |
| LAMA1 | E | NA | dbEMT | 0.5511 | 35 | 37.2 |
| LATS1 | E | NA | dbEMT | 1.60E-06 | 28.09 | 44 |
| LCN2 | E | E | Tan et al., 2014 | 0.0772 | 228.85 | 216.66 |
| LETMD1 | E | NA | dbEMT | 1.10E-16 | 35 | 75 |
| LIMA1 | E | NA | dbEMT | 1.10E-06 | 39 | 61.92 |
| LLGL2 | E | E | Tan et al., 2014 | 0.6439 | 51 | 47 |

| | | | | | | |
|---|---|---|---|---|---|---|
| LOXL2 | E | M | Tan et al., 2014 | 0.0004 | 228.85 | 216.66 |
| LOXL3 | E | NA | dbEMT | 0.0497 | 33 | 39 |
| LRRC1 | E | E | Tan et al., 2014 | 4.80E-05 | 216.66 | 191.21 |
| LSR | E | E | Tan et al., 2014 | 2.90E-11 | 69.6 | 35.98 |
| LY75 | E | E | Tan et al., 2014 | 0.0002 | 191.21 | 228.85 |
| LY96 | E | M | Tan et al., 2014 | 0.1804 | 216.66 | 191.21 |
| LYPD3 | E | NA | dbEMT | 0.0001 | 191.21 | 228.85 |
| MALL | E | E | Tan et al., 2014 | 0.7841 | 228.85 | 216.66 |
| MAP2K1 | E | NA | dbEMT | 0.8283 | 50 | 47 |
| MAP7 | E | E | Tan et al., 2014 | 0.002 | 216.66 | 191.21 |
| MAPK13 | E | E | Tan et al., 2014 | 0.0032 | 58 | 40.8 |
| MAPK3 | E | NA | dbEMT | 3.80E-05 | 228.85 | 216.66 |
| MAPK8 | E | NA | dbEMT | 1.00E-16 | 95 | 171.43 |
| MARVELD3 | E | NA | dbEMT | 0.0228 | 36.96 | 35 |
| MET | E | NA | dbEMT | 0.3528 | 47.4 | 49.45 |
| MGAT3 | E | NA | dbEMT | 0.085 | 45.37 | 53.36 |
| MGST2 | E | E | Tan et al., 2014 | 0.0001 | 216.66 | 228.85 |
| MIR200C | E | NA | dbEMT | n/a | n/a | n/a |
| MIR221 | E | NA | dbEMT | n/a | n/a | n/a |
| MIR222 | E | NA | dbEMT | n/a | n/a | n/a |
| MIR23A | E | NA | dbEMT | n/a | n/a | n/a |
| MIR30A | E | NA | dbEMT | n/a | n/a | n/a |
| MIR33A | E | NA | dbEMT | n/a | n/a | n/a |
| MKL2 | E | NA | dbEMT | 0.0032 | 35.04 | 68.4 |
| MPZL2 | E | E | Tan et al., 2014 | 3.80E-05 | 39 | 30.42 |
| MST1R | E | E | Tan et al., 2014 | 1.00E-16 | 191.21 | 216.66 |
| MSX2 | E | NA | dbEMT | 0.0228 | 41.72 | 57 |
| MTA1 | E | NA | dbEMT | 0.3528 | 191.21 | 216.66 |
| MTA3 | E | NA | dbEMT | 0.085 | 27 | 49 |
| MTOR | E | NA | dbEMT | 0.0001 | 42.96 | 57 |
| MTUS1 | E | E | Tan et al., 2014 | 2.20E-11 | 39 | 61 |
| MUC1 | E | E | Tan et al., 2014 | 0.7243 | 228.85 | 216.66 |
| MUC2 | E | NA | dbEMT | 5.10E-06 | 216.66 | 228.85 |
| MYC | E | NA | dbEMT | 0.0003 | 228.85 | 216.66 |
| MYH10 | E | M | Tan et al., 2014 | 2.70E-06 | 228.85 | 191.21 |
| MYO1D | E | E | Tan et al., 2014 | 5.30E-09 | 191.21 | 216.66 |
| MYO5C | E | E | Tan et al., 2014 | 3.90E-05 | 185.16 | 216.66 |
| MYO6 | E | E | Tan et al., 2014 | 5.00E-07 | 191.21 | 216.66 |
| NDRG1 | E | NA | dbEMT | 0.0019 | 216.66 | 184.04 |
| NFKB1 | E | NA | dbEMT | 1.20E-08 | 191.21 | 216.66 |
| NOTCH1 | E | NA | dbEMT | 0.0493 | 191.21 | 216.66 |
| NOTCH2 | E | NA | dbEMT | 0.2878 | 44.17 | 55 |
| NQO1 | E | E | Tan et al., 2014 | 0.207 | 216.66 | 185.16 |
| NRP2 | E | NA | dbEMT | 1.30E-14 | 37 | 64.73 |
| NUMB | E | NA | dbEMT | 0.0047 | 54 | 43 |
| OAS1 | E | E | Tan et al., 2014 | 2.20E-09 | 216.66 | 191.21 |
| OCLN | E | E | Tan et al., 2014 | 3.30E-16 | 25.2 | 50.4 |
| OR7E14P | E | E | Tan et al., 2014 | 0.9091 | 29 | 42 |
| OVOL1 | E | E | Tan et al., 2014 | 0.0049 | 184.04 | 216.66 |
| OVOL2 | E | E | Tan et al., 2014 | 4.70E-06 | 31.51 | 43 |

| | | | | | | |
|---|---|---|---|---|---|---|
| PAG1 | E | NA | dbEMT | 9.00E-06 | 228.85 | 216.66 |
| PAK1 | E | NA | dbEMT | 0.4208 | 42 | 57 |
| PARD3 | E | NA | dbEMT | 0.5676 | 228.85 | 216.66 |
| PARP1 | E | NA | dbEMT | 6.60E-09 | 216.66 | 184.04 |
| PCMT1 | E | NA | dbEMT | 0.0009 | 51 | 46.8 |
| PDGFB | E | NA | dbEMT | 5.20E-06 | 52.9 | 45 |
| PDPN | E | NA | dbEMT | 1.00E-04 | 38 | 33 |
| PERP | E | E | Tan et al., 2014 | 2.10E-09 | 35 | 36.9 |
| PHLDA1 | E | NA | dbEMT | 0.0107 | 228.85 | 173.2 |
| PHLDA2 | E | E | Tan et al., 2014 | 0.0569 | 216.66 | 185.16 |
| PIN1 | E | NA | dbEMT | 2.30E-05 | 228.85 | 216.66 |
| PKP3 | E | E | Tan et al., 2014 | 0.4644 | 228.85 | 216.66 |
| PLAUR | E | NA | dbEMT | 0.1181 | 185.16 | 216.66 |
| PLLP | E | E | Tan et al., 2014 | 0.0983 | 48.96 | 48 |
| PLS1 | E | E | Tan et al., 2014 | 0.4374 | 33 | 38 |
| POF1B | E | E | Tan et al., 2014 | 4.00E-10 | 228.85 | 216.66 |
| PPAP2C | E | E | Tan et al., 2014 | 0.0007 | 228.85 | 216.66 |
| PPFIBP2 | E | E | Tan et al., 2014 | 0.1244 | 228.85 | 216.66 |
| PPL | E | E | Tan et al., 2014 | 0.7174 | 216.66 | 228.85 |
| PRDX1 | E | NA | dbEMT | 7.10E-13 | 32 | 39 |
| PRKCE | E | NA | dbEMT | 0.5707 | 52.32 | 44.22 |
| PROM1 | E | NA | dbEMT | 0.1483 | 228.85 | 216.66 |
| PRR15L | E | E | Tan et al., 2014 | n/a | n/a | n/a |
| PRSS8 | E | E | Tan et al., 2014 | 0.1483 | 52.83 | 45.84 |
| PSCA | E | E | Tan et al., 2014 | 3.50E-06 | 185.16 | 216.66 |
| PTHLH | E | NA | dbEMT | 1.10E-06 | 216.66 | 191.21 |
| PTK2 | E | NA | dbEMT | 0.5338 | 29 | 45.6 |
| PTK6 | E | E | Tan et al., 2014 | 0.375 | 216.66 | 228.85 |
| PTPRC | E | M | Tan et al., 2014 | 0.1231 | 228.85 | 216.66 |
| PTPRF | E | E | Tan et al., 2014 | 0.0629 | 43.7 | 57.6 |
| PTPRZ1 | E | NA | dbEMT | 3.60E-05 | 191.21 | 216.66 |
| PYCARD | E | E | Tan et al., 2014 | 0.0006 | 28.96 | 42 |
| RAB11FIP1 | E | E | Tan et al., 2014 | 3.30E-06 | 44 | 55.13 |
| RAB20 | E | E | Tan et al., 2014 | 0.0004 | 228.85 | 185.16 |
| RAB25 | E | E | Tan et al., 2014 | 0.0198 | 164.3 | 228.85 |
| RABGAP1L | E | E | Tan et al., 2014 | 0.0005 | 228.85 | 216.66 |
| RAPGEFL1 | E | E | Tan et al., 2014 | 1.10E-05 | 51.36 | 46.49 |
| RBM47 | E | E | Tan et al., 2014 | 0.044 | 228.85 | 216.66 |
| RGS3 | E | NA | dbEMT | 0.0006 | 216.66 | 228.85 |
| RHOA | E | NA | dbEMT | 0.5855 | 49 | 47 |
| RHOD | E | E | Tan et al., 2014 | 3.70E-08 | 48 | 48.16 |
| RNF128 | E | E | Tan et al., 2014 | 2.90E-06 | 216.66 | 161.29 |
| S100A14 | E | E | Tan et al., 2014 | 0.4268 | 38 | 32 |
| S100P | E | E | Tan et al., 2014 | 0.5239 | 191.21 | 216.66 |
| SCEL | E | E | Tan et al., 2014 | 0.4037 | 228.85 | 216.66 |
| SCNN1A | E | E | Tan et al., 2014 | 0.6697 | 57 | 42 |
| SCRIB | E | NA | dbEMT | 0.374 | 44.5 | 52.54 |
| SDC1 | E | E | Tan et al., 2014 | 2.60E-13 | 39 | 31.18 |
| SDC4 | E | E | Tan et al., 2014 | 0.4125 | 228.85 | 185.16 |
| SEMA7A | E | NA | dbEMT | 8.20E-09 | 45.6 | 53.36 |

| | | | | | | |
|---|---|---|---|---|---|---|
| SERINC5 | E | E | Tan et al., 2014 | n/a | n/a | n/a |
| SFN | E | E | Tan et al., 2014 | 0.0329 | 52.83 | 45.84 |
| SFRP1 | E | M | Tan et al., 2014 | 0.0353 | 185.16 | 216.66 |
| SFTPC | E | NA | dbEMT | 1.70E-14 | 216.66 | 191.21 |
| SH2D3A | E | E | Tan et al., 2014 | 4.90E-07 | 29 | 45.6 |
| SH3YL1 | E | E | Tan et al., 2014 | 0.0022 | 216.66 | 228.85 |
| SHANK2 | E | E | Tan et al., 2014 | 0.6314 | 228.85 | 216.66 |
| SIM2 | E | NA | dbEMT | 1.30E-05 | 43.7 | 57.6 |
| SLC16A5 | E | E | Tan et al., 2014 | 4.00E-05 | 191.21 | 216.66 |
| SLC22A18 | E | E | Tan et al., 2014 | 0.0991 | 28.96 | 42 |
| SLC2A3 | E | M | Tan et al., 2014 | 0.616 | 44 | 55.13 |
| SLC35A3 | E | E | Tan et al., 2014 | 0.0562 | 228.85 | 185.16 |
| SLC37A1 | E | E | Tan et al., 2014 | 0.1087 | 164.3 | 228.85 |
| SLC39A6 | E | NA | dbEMT | 9.90E-08 | 228.85 | 216.66 |
| SLC44A4 | E | E | Tan et al., 2014 | 4.00E-09 | 51.36 | 46.49 |
| SLC9A3R1 | E | E | Tan et al., 2014 | 0.0028 | 228.85 | 216.66 |
| SLIT2 | E | M | Tan et al., 2014 | 1.10E-15 | 216.66 | 228.85 |
| SLPI | E | E | Tan et al., 2014 | 6.00E-08 | 49 | 47 |
| SMAD2 | E | NA | dbEMT | 0.5183 | 48 | 48.16 |
| SMAD3 | E | NA | dbEMT | 0.0069 | 216.66 | 161.29 |
| SMAD9 | E | NA | dbEMT | 2.20E-12 | 38 | 32 |
| SNW1 | E | NA | dbEMT | 0.0002 | 191.21 | 216.66 |
| SORD | E | E | Tan et al., 2014 | 0.0029 | 228.85 | 216.66 |
| SORL1 | E | E | Tan et al., 2014 | 8.50E-08 | 57 | 42 |
| SOX9 | E | NA | dbEMT | 0.0369 | 44.5 | 52.54 |
| SP1 | E | NA | dbEMT | 5.20E-08 | 39 | 31.18 |
| SPAG1 | E | E | Tan et al., 2014 | 0.471 | 228.85 | 185.16 |
| SPDEF | E | E | Tan et al., 2014 | 3.40E-07 | 185.16 | 216.66 |
| SPINT1 | E | E | Tan et al., 2014 | 0.7666 | 58.52 | 40.37 |
| SPINT2 | E | E | Tan et al., 2014 | 7.00E-08 | 228.85 | 216.66 |
| SPP1 | E | NA | dbEMT | 4.90E-11 | 27 | 45.6 |
| SPRR2A | E | NA | dbEMT | n/a | n/a | n/a |
| SSH3 | E | E | Tan et al., 2014 | 0.0329 | 216.66 | 228.85 |
| ST14 | E | E | Tan et al., 2014 | 0.0353 | 44 | 54 |
| STAP2 | E | E | Tan et al., 2014 | 1.70E-14 | 191.21 | 216.66 |
| STK11 | E | NA | dbEMT | 4.90E-07 | 44.5 | 54 |
| STYK1 | E | E | Tan et al., 2014 | 0.0022 | 32 | 39 |
| SYNGR2 | E | E | Tan et al., 2014 | 0.6314 | 27.3 | 46 |
| TACSTD2 | E | E | Tan et al., 2014 | 1.30E-05 | 228.85 | 216.66 |
| TBX2 | E | NA | dbEMT | 4.00E-05 | 173.2 | 216.66 |
| TBX20 | E | NA | dbEMT | 0.0991 | 25.08 | 58.15 |
| TCF21 | E | NA | dbEMT | 0.616 | 191.21 | 216.66 |
| TCF7 | E | NA | dbEMT | 0.0562 | 51 | 46 |
| TDGF1 | E | NA | dbEMT | 0.1087 | 228.85 | 191.21 |
| TFCP2 | E | NA | dbEMT | 9.90E-08 | 27 | 57 |
| TFF1 | E | E | Tan et al., 2014 | 4.00E-09 | 31 | 40.44 |
| TGFA | E | E | Tan et al., 2014 | 0.0028 | 44.4 | 54.77 |
| TGFB1 | E | NA | dbEMT | 1.10E-15 | 228.85 | 216.66 |
| TGFBR3 | E | NA | dbEMT | 6.00E-08 | 228.85 | 216.66 |
| THBD | E | NA | dbEMT | 0.5183 | 228.85 | 216.66 |

| | | | | | | |
|---|---|---|---|---|---|---|
| TIAM1 | E | NA | dbEMT | 0.0069 | 45 | 52.3 |
| TJP2 | E | E | Tan et al., 2014 | 2.20E-12 | 228.85 | 216.66 |
| TJP3 | E | E | Tan et al., 2014 | 0.0002 | 216.66 | 191.21 |
| TMC5 | E | E | Tan et al., 2014 | 0.0029 | 184.04 | 216.66 |
| TMC6 | E | E | Tan et al., 2014 | 8.50E-08 | 37.25 | 69.6 |
| TMEM30B | E | E | Tan et al., 2014 | 0.0369 | 228.85 | 216.66 |
| TMPRSS4 | E | E | Tan et al., 2014 | 5.20E-08 | 216.66 | 228.85 |
| TNF | E | NA | dbEMT | 0.471 | 37.2 | 72.2 |
| TNFSF13 | E | E | Tan et al., 2014 | 3.40E-07 | 31 | 39 |
| TOB1 | E | E | Tan et al., 2014 | 0.7666 | 34 | 37.2 |
| TOM1L1 | E | E | Tan et al., 2014 | 7.00E-08 | 228.85 | 216.66 |
| TP63 | E | NA | dbEMT | 4.90E-11 | 228.85 | 216.66 |
| TP73 | E | NA | dbEMT | 0.4266 | 60 | 43 |
| TPD52 | E | E | Tan et al., 2014 | 0.5736 | 216.66 | 191.21 |
| TRPM4 | E | E | Tan et al., 2014 | 1.00E-12 | 171.43 | 216.66 |
| TSPAN1 | E | E | Tan et al., 2014 | 0.0289 | 228.85 | 185.16 |
| TSPAN13 | E | E | Tan et al., 2014 | 0.9798 | 228.85 | 216.66 |
| TSPAN15 | E | E | Tan et al., 2014 | 0.6097 | 36 | 72 |
| TTC39A | E | E | Tan et al., 2014 | 0.3555 | 51.36 | 45.27 |
| TUBB6 | E | M | Tan et al., 2014 | 1.90E-05 | 228.85 | 216.66 |
| TUFT1 | E | E | Tan et al., 2014 | 0.0042 | 216.66 | 185.16 |
| TXNIP | E | NA | dbEMT | 1.30E-13 | 171.43 | 216.66 |
| UCHL1 | E | M | Tan et al., 2014 | 1.50E-05 | 228.85 | 216.66 |
| UGT1A1 | E | E | Tan et al., 2014 | 0.5665 | 66 | 37 |
| VAMP8 | E | E | Tan et al., 2014 | 1.00E-13 | 216.66 | 191.21 |
| VAV3 | E | E | Tan et al., 2014 | 0.4425 | 43 | 58 |
| VDR | E | NA | dbEMT | 5.40E-05 | 216.66 | 228.85 |
| VEGFA | E | NA | dbEMT | 0.0183 | 44.5 | 56.6 |
| VGLL1 | E | E | Tan et al., 2014 | 2.80E-12 | 58.65 | 41 |
| VSNL1 | E | NA | dbEMT | 0.5664 | 163.46 | 216.66 |
| VTN | E | NA | dbEMT | 1.80E-07 | 35 | 37 |
| WISP3 | E | NA | dbEMT | 3.10E-06 | 216.66 | 171.43 |
| WT1 | E | NA | dbEMT | 0.0057 | 76.02 | 35.09 |
| XBP1 | E | E | Tan et al., 2014 | 0.0149 | 50 | 47 |
| YAP1 | E | NA | dbEMT | 0.017 | 57.6 | 27 |
| YBX1 | E | NA | dbEMT | 0.0004 | 216.66 | 228.85 |
| YWHAZ | E | NA | dbEMT | 1.00E-15 | 191.21 | 216.66 |
| YY1 | E | NA | dbEMT | 0.3429 | 60 | 40.44 |
| ZBTB33 | E | NA | dbEMT | 1.00E-16 | 184.04 | 216.66 |
| ZFYVE9 | E | NA | dbEMT | 6.70E-16 | 216.66 | 191.21 |
| ZNF165 | E | E | Tan et al., 2014 | 0.1806 | 50 | 46.32 |
| ZNF217 | E | NA | dbEMT | 3.20E-12 | 184.04 | 216.66 |
| ZYX | E | NA | dbEMT | 1.90E-07 | 228.85 | 191.21 |
| AKAP2 | M1 | M | Tan et al., 2014 | 0.001 | 36.9 | 35.09 |
| AKT1 | M1 | NA | dbEMT | 6.20E-05 | 27 | 48 |
| AKT3 | M1 | M | Tan et al., 2014 | 1.40E-10 | 216.66 | 228.85 |
| ANGPTL2 | M1 | M | Tan et al., 2014 | 0.0003 | 228.85 | 216.66 |
| AP1S2 | M1 | M | Tan et al., 2014 | 0.7127 | 27 | 49.2 |
| ARHGAP32 | M1 | E | Tan et al., 2014 | 2.40E-05 | 216.66 | 228.85 |
| ATM | M1 | NA | dbEMT | 0.3896 | 228.85 | 216.66 |

| | | | | | | |
|---|---|---|---|---|---|---|
| ATP1B1 | M1 | E | Tan et al., 2014 | 0.9761 | 57.3 | 40.56 |
| AXIN2 | M1 | NA | dbEMT | 2.10E-07 | 228.85 | 185.16 |
| BIRC2 | M1 | NA | dbEMT | 0.0062 | 228.85 | 216.66 |
| BMP2 | M1 | NA | dbEMT | 1.00E-07 | 31 | 47.28 |
| CALD1 | M1 | M | Tan et al., 2014 | 5.50E-08 | 185.16 | 216.66 |
| CAMK1D | M1 | NA | dbEMT | 0.5616 | 37 | 34.8 |
| CCL2 | M1 | M | Tan et al., 2014 | 0.0199 | 40 | 60.6 |
| CD163 | M1 | M | Tan et al., 2014 | 0.0056 | 40.44 | 60 |
| CD274 | M1 | NA | dbEMT | 0.5148 | 216.66 | 228.85 |
| CD44 | M1 | NA | dbEMT | 0.8939 | 216.66 | 191.21 |
| CDH11 | M1 | M | Tan et al., 2014 | 7.20E-08 | 61.64 | 40 |
| CDK14 | M1 | M | Tan et al., 2014 | n/a | n/a | n/a |
| CDX2 | M1 | NA | dbEMT | 3.90E-09 | 61 | 39 |
| CEACAM7 | M1 | E | Tan et al., 2014 | 0.5302 | 228.85 | 216.66 |
| CHN1 | M1 | M | Tan et al., 2014 | 6.30E-05 | 228.85 | 216.66 |
| CHRDL1 | M1 | M | Tan et al., 2014 | 6.50E-06 | 228.85 | 216.66 |
| CLDN3 | M1 | E | Tan et al., 2014 | 0.0517 | 184.04 | 216.66 |
| CLIC4 | M1 | M | Tan et al., 2014 | 2.40E-08 | 216.66 | 228.85 |
| COL15A1 | M1 | M | Tan et al., 2014 | 4.60E-06 | 47.04 | 50.86 |
| COL6A1 | M1 | M | Tan et al., 2014 | 0.0004 | 42 | 56.58 |
| COL6A2 | M1 | M | Tan et al., 2014 | 0.0254 | 45 | 52.9 |
| COL8A2 | M1 | NA | dbEMT | 0.0002 | 216.66 | 228.85 |
| COLEC12 | M1 | M | Tan et al., 2014 | 0.0038 | 191.21 | 216.66 |
| CRYAB | M1 | M | Tan et al., 2014 | 0.0013 | 46.68 | 52.2 |
| CTSK | M1 | M | Tan et al., 2014 | 0.0021 | 228.85 | 216.66 |
| CUX1 | M1 | NA | dbEMT | 0.3705 | 48.99 | 47.88 |
| CXCL12 | M1 | M | Tan et al., 2014 | 0.003 | 47.08 | 48 |
| CXCL13 | M1 | M | Tan et al., 2014 | 0.1049 | 36 | 36.04 |
| CXCL5 | M1 | NA | dbEMT | 6.10E-09 | 38 | 71 |
| CXCR4 | M1 | M | Tan et al., 2014 | 0.1052 | 216.66 | 228.85 |
| CYB561 | M1 | E | Tan et al., 2014 | 0.1408 | 55.99 | 43.56 |
| CYR61 | M1 | NA | dbEMT | 0.0028 | 40.37 | 58.65 |
| DDR2 | M1 | M | Tan et al., 2014 | 0.2749 | 35.09 | 36.44 |
| DENND5A | M1 | M | Tan et al., 2014 | 0.0474 | 228.85 | 216.66 |
| DPT | M1 | M | Tan et al., 2014 | 0.5559 | 216.66 | 228.85 |
| DSE | M1 | M | Tan et al., 2014 | 1.30E-13 | 216.66 | 228.85 |
| ECM2 | M1 | M | Tan et al., 2014 | 2.80E-05 | 228.85 | 216.66 |
| EDN1 | M1 | NA | dbEMT | 0.0269 | 41 | 59 |
| EFEMP2 | M1 | M | Tan et al., 2014 | 1.10E-05 | 37 | 68.48 |
| ELF5 | M1 | E | Tan et al., 2014 | 0.0274 | 47 | 48.16 |
| ENPP2 | M1 | M | Tan et al., 2014 | 0.0094 | 228.85 | 216.66 |
| EPO | M1 | NA | dbEMT | 0.1522 | 37 | 64.73 |
| F13A1 | M1 | M | Tan et al., 2014 | 5.40E-08 | 228.85 | 191.21 |
| FAM174B | M1 | E | Tan et al., 2014 | 0.0414 | 184.04 | 216.66 |
| FAP | M1 | M | Tan et al., 2014 | 1.50E-05 | 30.03 | 44 |
| FBLN5 | M1 | NA | dbEMT | 9.00E-11 | 216.66 | 191.21 |
| FBP1 | M1 | E | Tan et al., 2014 | 0.5099 | 37.8 | 64 |
| FGF2 | M1 | NA | dbEMT | 9.10E-06 | 185.16 | 216.66 |
| FGFR2 | M1 | NA | dbEMT | 1.80E-07 | 191.21 | 216.66 |
| FGL2 | M1 | M | Tan et al., 2014 | 9.50E-06 | 184.04 | 216.66 |

| | | | | | | |
|---|---|---|---|---|---|---|
| FHL1 | M1 | M | Tan et al., 2014 | 5.70E-07 | 41.04 | 60 |
| FLRT2 | M1 | M | Tan et al., 2014 | 4.90E-06 | 185.16 | 216.66 |
| FOXC1 | M1 | NA | dbEMT | 0.1473 | 216.66 | 228.85 |
| FSCN2 | M1 | NA | dbEMT | 2.40E-08 | 39 | 31.8 |
| GAB2 | M1 | NA | dbEMT | 0.0281 | 66 | 38 |
| GEM | M1 | M | Tan et al., 2014 | 0.0195 | 228.85 | 216.66 |
| GFPT2 | M1 | M | Tan et al., 2014 | 1.90E-07 | 35 | 37.52 |
| GIMAP4 | M1 | M | Tan et al., 2014 | 6.00E-05 | 43.2 | 58.15 |
| GLIPR2 | M1 | NA | dbEMT | 3.90E-05 | 228.85 | 216.66 |
| GREM1 | M1 | M | Tan et al., 2014 | 4.20E-05 | 216.66 | 228.85 |
| GRIN1 | M1 | NA | dbEMT | 0.4171 | 228.85 | 216.66 |
| GSC | M1 | M | Tan et al., 2014 | 6.90E-07 | 31.56 | 40.53 |
| GSN | M1 | NA | dbEMT | 0.0001 | 52.3 | 45.01 |
| GUCY1B3 | M1 | M | Tan et al., 2014 | 0.7886 | 28.8 | 50 |
| GZMK | M1 | M | Tan et al., 2014 | 3.20E-05 | 216.66 | 228.85 |
| HAS2 | M1 | NA | dbEMT | 0.4407 | 32.03 | 40.44 |
| HMGA2 | M1 | NA | dbEMT | 6.50E-08 | 37.2 | 33.64 |
| HMOX1 | M1 | NA | dbEMT | 0.0792 | 216.66 | 191.21 |
| HNF4A | M1 | NA | dbEMT | 0.0247 | 38 | 62.36 |
| HOXB7 | M1 | NA | dbEMT | 0.0652 | 38.4 | 68.4 |
| HOXB9 | M1 | NA | dbEMT | 6.40E-10 | 51 | 46 |
| HS3ST3B1 | M1 | NA | dbEMT | 0.1448 | 68.75 | 37.64 |
| HSPB2 | M1 | NA | dbEMT | 0.0025 | 42 | 30.72 |
| IGF1 | M1 | M | Tan et al., 2014 | 0.361 | 38 | 69.6 |
| IL10RA | M1 | M | Tan et al., 2014 | 0.2438 | 44.6 | 54.96 |
| ITGA5 | M1 | NA | dbEMT | 4.20E-07 | 27.96 | 48 |
| ITGB1 | M1 | NA | dbEMT | 5.30E-10 | 38 | 61 |
| ITGB6 | M1 | E | Tan et al., 2014 | 0.4375 | 228.85 | 191.21 |
| ITM2A | M1 | M | Tan et al., 2014 | 6.00E-09 | 216.66 | 191.21 |
| JAK2 | M1 | NA | dbEMT | 0.0937 | 228.85 | 216.66 |
| JAM2 | M1 | M | Tan et al., 2014 | 4.20E-12 | 228.85 | 216.66 |
| JAM3 | M1 | M | Tan et al., 2014 | 0.0846 | 40.3 | 65.04 |
| KCNH1 | M1 | NA | dbEMT | 4.50E-08 | 55.66 | 43.56 |
| KCNJ8 | M1 | M | Tan et al., 2014 | 1.20E-06 | 185.16 | 228.85 |
| KDELC1 | M1 | M | Tan et al., 2014 | 0.0176 | 45 | 53.04 |
| KIAA1462 | M1 | M | Tan et al., 2014 | 0.0024 | 228.85 | 191.21 |
| KIT | M1 | NA | dbEMT | 6.60E-05 | 60.3 | 42 |
| KRT8 | M1 | E | Tan et al., 2014 | 0.0053 | 37 | 66 |
| LEF1 | M1 | NA | dbEMT | 3.00E-07 | 105.6 | 171.43 |
| LEP | M1 | NA | dbEMT | 0.0011 | 216.66 | 173.2 |
| LEPRE1 | M1 | M | Tan et al., 2014 | 4.80E-09 | 228.85 | 191.21 |
| LGALS1 | M1 | M | Tan et al., 2014 | 0.1714 | 47.28 | 49 |
| LHFP | M1 | M | Tan et al., 2014 | 0.5578 | 42.05 | 56 |
| LIMS1 | M1 | NA | dbEMT | 9.10E-05 | 31.2 | 42 |
| MAGED1 | M1 | NA | dbEMT | 9.00E-10 | 43 | 61 |
| MAP1B | M1 | M | Tan et al., 2014 | 3.90E-13 | 191.21 | 216.66 |
| MDK | M1 | NA | dbEMT | 3.40E-08 | 49.02 | 48 |
| MEOX2 | M1 | M | Tan et al., 2014 | 0.9971 | 228.85 | 191.21 |
| MIR194-1 | M1 | NA | dbEMT | n/a | n/a | n/a |
| MIR34C | M1 | NA | dbEMT | 0.8392 | 57 | 43 |

| | | | | | | |
|---|---|---|---|---|---|---|
| MIR9-1 | M1 | NA | dbEMT | n/a | n/a | n/a |
| MKL1 | M1 | NA | dbEMT | 0.05 | 56 | 40.8 |
| MLPH | M1 | E | Tan et al., 2014 | 0.0011 | 216.66 | 185.16 |
| MMP14 | M1 | NA | dbEMT | 1.90E-06 | 228.85 | 216.66 |
| MMP3 | M1 | NA | dbEMT | 6.60E-09 | 228.85 | 191.21 |
| MMP9 | M1 | NA | dbEMT | 0.9385 | 35 | 36.96 |
| MSN | M1 | M | Tan et al., 2014 | 0.8334 | 228.85 | 216.66 |
| MUC4 | M1 | NA | dbEMT | 0.0065 | 41.69 | 58.52 |
| MXRA7 | M1 | M | Tan et al., 2014 | 0.0448 | 228.85 | 216.66 |
| MYCN | M1 | NA | dbEMT | 0.3122 | 228.85 | 216.66 |
| MYH14 | M1 | E | Tan et al., 2014 | 0.4864 | 228.85 | 216.66 |
| MYL9 | M1 | M | Tan et al., 2014 | 1.30E-06 | 228.85 | 216.66 |
| NAP1L3 | M1 | M | Tan et al., 2014 | 0.9801 | 228.85 | 216.66 |
| NUAK1 | M1 | M | Tan et al., 2014 | 0.0218 | 216.66 | 184.04 |
| OLFML2B | M1 | M | Tan et al., 2014 | 0.0002 | 47 | 50 |
| PAX2 | M1 | NA | dbEMT | 0.6865 | 44 | 53.04 |
| PCOLCE | M1 | M | Tan et al., 2014 | 0.0134 | 216.66 | 228.85 |
| PDZRN3 | M1 | M | Tan et al., 2014 | 0.0088 | 216.66 | 191.21 |
| PIK3CA | M1 | NA | dbEMT | 0.0486 | 52.3 | 46.49 |
| PLEKHO1 | M1 | M | Tan et al., 2014 | 0.0106 | 45 | 53.16 |
| PLN | M1 | M | Tan et al., 2014 | 2.00E-07 | 228.85 | 185.16 |
| PMP22 | M1 | M | Tan et al., 2014 | 0.2577 | 216.66 | 228.85 |
| POPDC3 | M1 | M | Tan et al., 2014 | 0.0073 | 32 | 39 |
| POSTN | M1 | NA | dbEMT | 0.4699 | 46 | 50 |
| PRKCA | M1 | NA | dbEMT | 0.0135 | 228.85 | 185.16 |
| PTGIS | M1 | M | Tan et al., 2014 | 0.3289 | 30 | 44.4 |
| RAC1 | M1 | NA | dbEMT | 0.0056 | 42 | 61 |
| RARRES2 | M1 | M | Tan et al., 2014 | 0.0004 | 216.66 | 228.85 |
| RDX | M1 | NA | dbEMT | 0.1448 | 185.16 | 216.66 |
| RECK | M1 | M | Tan et al., 2014 | 0.0155 | 228.85 | 216.66 |
| RNF111 | M1 | NA | dbEMT | 0.1137 | 44.61 | 52.3 |
| ROCK1 | M1 | NA | dbEMT | 0.4053 | 216.66 | 191.21 |
| RUNX3 | M1 | NA | dbEMT | 0.3587 | 39 | 63.6 |
| SACS | M1 | M | Tan et al., 2014 | 8.80E-06 | 228.85 | 216.66 |
| SDC2 | M1 | M | Tan et al., 2014 | 1.10E-05 | 39 | 32.4 |
| SERPINE1 | M1 | M | Tan et al., 2014 | 0.0335 | 228.85 | 216.66 |
| SERPINF1 | M1 | M | Tan et al., 2014 | 0.0005 | 216.66 | 191.21 |
| SFRP4 | M1 | M | Tan et al., 2014 | 0.003 | 228.85 | 216.66 |
| SH2B3 | M1 | M | Tan et al., 2014 | 0.029 | 228.85 | 191.21 |
| SHH | M1 | NA | dbEMT | 2.30E-05 | 216.66 | 185.16 |
| SIX1 | M1 | NA | dbEMT | 1.30E-08 | 171.43 | 216.66 |
| SMAD7 | M1 | NA | dbEMT | 4.20E-08 | 228.85 | 216.66 |
| SNAI1 | M1 | M | Tan et al., 2014 | 0.0042 | 216.66 | 191.21 |
| SPARC | M1 | M | Tan et al., 2014 | 0.1208 | 184.04 | 216.66 |
| SRC | M1 | NA | dbEMT | 0.0673 | 37.35 | 72.49 |
| SRPX | M1 | M | Tan et al., 2014 | 0.0101 | 49 | 47.51 |
| STAT5A | M1 | NA | dbEMT | 3.30E-08 | 44 | 57 |
| STAT5B | M1 | NA | dbEMT | 0.9245 | 38.57 | 60 |
| STON1 | M1 | M | Tan et al., 2014 | 1.00E-16 | 228.85 | 216.66 |
| SYNE1 | M1 | M | Tan et al., 2014 | 1.00E-16 | 38 | 32 |

| | | | | | | |
|---|---|---|---|---|---|---|
| SYT11 | M1 | M | Tan et al., 2014 | 0.0444 | 40.08 | 61.05 |
| TAGLN | M1 | M | Tan et al., 2014 | 1.80E-12 | 171.43 | 216.66 |
| TCF3 | M1 | NA | dbEMT | 1.20E-13 | 216.66 | 191.21 |
| TFF3 | M1 | E | Tan et al., 2014 | 0.8548 | 39 | 32.82 |
| TGFB1I1 | M1 | M | Tan et al., 2014 | 0.0044 | 228.85 | 216.66 |
| TGFBR1 | M1 | NA | dbEMT | 0.0005 | 216.66 | 191.21 |
| TGM2 | M1 | NA | dbEMT | 0.973 | 216.66 | 228.85 |
| TM4SF5 | M1 | NA | dbEMT | 0.7866 | 228.85 | 185.16 |
| TMEFF1 | M1 | M | Tan et al., 2014 | 5.70E-06 | 191.21 | 228.85 |
| TMPRSS2 | M1 | E | Tan et al., 2014 | 2.80E-09 | 216.66 | 191.21 |
| TNC | M1 | M | Tan et al., 2014 | 0.4698 | 228.85 | 191.21 |
| TNS1 | M1 | M | Tan et al., 2014 | 0.1884 | 50.3 | 47.4 |
| TOX3 | M1 | E | Tan et al., 2014 | 0.0972 | 216.66 | 191.21 |
| TPM2 | M1 | M | Tan et al., 2014 | 0.0012 | 228.85 | 216.66 |
| TRPC1 | M1 | M | Tan et al., 2014 | 0.176 | 216.66 | 228.85 |
| TSPAN8 | M1 | E | Tan et al., 2014 | 1.10E-07 | 228.85 | 185.16 |
| TUBA1A | M1 | M | Tan et al., 2014 | 0.0045 | 228.85 | 216.66 |
| VCAM1 | M1 | M | Tan et al., 2014 | 0.0004 | 228.85 | 191.21 |
| VSIG4 | M1 | M | Tan et al., 2014 | 2.40E-07 | 26 | 45 |
| WIPF1 | M1 | M | Tan et al., 2014 | 0.7863 | 24.07 | 56.4 |
| WNT1 | M1 | NA | dbEMT | 0.1527 | 184.04 | 216.66 |
| WNT3A | M1 | NA | dbEMT | n/a | n/a | n/a |
| WWTR1 | M1 | M | Tan et al., 2014 | 0.8466 | 228.85 | 216.66 |
| ACVR1 | M2 | NA | dbEMT | 9.70E-05 | 228.85 | 216.66 |
| BICC1 | M2 | M | Tan et al., 2014 | 0.0017 | 228.85 | 216.66 |
| BNC2 | M2 | M | Tan et al., 2014 | 0.2897 | 49.2 | 47 |
| BRAF | M2 | NA | dbEMT | 1.00E-10 | 60 | 39.95 |
| C1R | M2 | M | Tan et al., 2014 | 9.20E-09 | 228.85 | 216.66 |
| C1S | M2 | M | Tan et al., 2014 | 1.30E-13 | 228.85 | 216.66 |
| CDH13 | M2 | NA | dbEMT | 1.20E-06 | 228.85 | 191.21 |
| CDH2 | M2 | M | Tan et al., 2014 | 0.6864 | 43 | 55.43 |
| CDKN1B | M2 | NA | dbEMT | 0.011 | 228.85 | 216.66 |
| CDKN2A | M2 | NA | dbEMT | 0.0142 | 43 | 56 |
| CEP170 | M2 | M | Tan et al., 2014 | 0.3789 | 216.66 | 228.85 |
| COL5A2 | M2 | M | Tan et al., 2014 | 0.0009 | 29 | 41.95 |
| CTGF | M2 | NA | dbEMT | 0.218 | 191.21 | 216.66 |
| DAB2 | M2 | NA | dbEMT | 0.7456 | 228.85 | 216.66 |
| DCN | M2 | M | Tan et al., 2014 | 0.0188 | 216.66 | 191.21 |
| DNAJB6 | M2 | NA | dbEMT | 0.005 | 29 | 46 |
| EMP3 | M2 | M | Tan et al., 2014 | 0.022 | 44 | 54.7 |
| ENG | M2 | NA | dbEMT | 0.0007 | 38 | 68.48 |
| ERF | M2 | NA | dbEMT | 9.10E-05 | 228.85 | 216.66 |
| FBN1 | M2 | M | Tan et al., 2014 | 2.60E-05 | 216.66 | 191.21 |
| FN1 | M2 | M | Tan et al., 2014 | 9.00E-05 | 228.85 | 185.16 |
| FSTL1 | M2 | M | Tan et al., 2014 | 0.0951 | 50 | 47.28 |
| FYN | M2 | M | Tan et al., 2014 | 0.0694 | 57 | 40 |
| GIPC2 | M2 | NA | dbEMT | 0.0005 | 228.85 | 216.66 |
| GJA1 | M2 | M | Tan et al., 2014 | 2.30E-05 | 228.85 | 216.66 |
| HDAC6 | M2 | NA | dbEMT | 0.0142 | 46 | 50.86 |
| HEG1 | M2 | M | Tan et al., 2014 | 5.40E-11 | 48 | 48 |

| | | | | | | |
|---|---|---|---|---|---|---|
| HIF1A | M2 | NA | dbEMT | 0.0338 | 40.44 | 60 |
| HSPB1 | M2 | NA | dbEMT | 6.30E-13 | 51 | 47 |
| IGF1R | M2 | NA | dbEMT | 2.20E-09 | 191.21 | 216.66 |
| IGFBP5 | M2 | M | Tan et al., 2014 | 0.8651 | 52.32 | 46 |
| KDM3A | M2 | NA | dbEMT | 0.0746 | 43 | 57 |
| KLF4 | M2 | NA | dbEMT | 2.20E-05 | 228.85 | 216.66 |
| LOX | M2 | M | Tan et al., 2014 | 0.0225 | 216.66 | 191.21 |
| MAF | M2 | M | Tan et al., 2014 | 0.7329 | 191.21 | 216.66 |
| MAP3K4 | M2 | NA | dbEMT | 0.6733 | 34 | 37.52 |
| MAP3K7 | M2 | NA | dbEMT | 3.50E-07 | 50 | 47 |
| MCAM | M2 | NA | dbEMT | 0.4734 | 185.16 | 216.66 |
| MIR21 | M2 | NA | dbEMT | 0.4656 | 33.36 | 76 |
| MMP13 | M2 | NA | dbEMT | 0.9199 | 40.56 | 58 |
| MMP2 | M2 | M | Tan et al., 2014 | 0.0012 | 44.61 | 53.1 |
| MOXD1 | M2 | M | Tan et al., 2014 | 6.20E-07 | 228.85 | 216.66 |
| MPDZ | M2 | M | Tan et al., 2014 | 0.0239 | 31.51 | 38 |
| NCSTN | M2 | NA | dbEMT | 0.0007 | 216.66 | 191.21 |
| NOG | M2 | NA | dbEMT | 0.2196 | 45 | 55.9 |
| NR3C1 | M2 | M | Tan et al., 2014 | 0.4995 | 42 | 60 |
| OLFML3 | M2 | M | Tan et al., 2014 | 3.40E-08 | 216.66 | 228.85 |
| PDE4A | M2 | NA | dbEMT | 1.00E-16 | 216.66 | 228.85 |
| PDGFC | M2 | M | Tan et al., 2014 | 0.0002 | 228.85 | 216.66 |
| PLXNB2 | M2 | E | Tan et al., 2014 | 0.1499 | 28 | 46.8 |
| POU5F1 | M2 | NA | dbEMT | 0.0007 | 216.66 | 228.85 |
| PTEN | M2 | NA | dbEMT | 0.6247 | 173.2 | 216.66 |
| PTGDS | M2 | M | Tan et al., 2014 | 1.40E-07 | 50 | 46.68 |
| PTGS2 | M2 | NA | dbEMT | 0.0019 | 26 | 48 |
| PTN | M2 | NA | dbEMT | 0.0008 | 51 | 46 |
| PTX3 | M2 | M | Tan et al., 2014 | 0.3181 | 31.51 | 42 |
| RGS2 | M2 | M | Tan et al., 2014 | 0.6106 | 228.85 | 216.66 |
| ROR2 | M2 | NA | dbEMT | 0.0039 | 48 | 48 |
| RUNX1T1 | M2 | M | Tan et al., 2014 | 1.30E-07 | 37.64 | 34 |
| SAMSN1 | M2 | M | Tan et al., 2014 | 0.8113 | 43.56 | 58 |
| SEMA3E | M2 | NA | dbEMT | 6.60E-08 | 25 | 49.2 |
| SEMA4C | M2 | NA | dbEMT | 0.932 | 228.85 | 185.16 |
| SMAD4 | M2 | NA | dbEMT | 4.10E-08 | 184.04 | 216.66 |
| SNAI2 | M2 | M | Tan et al., 2014 | 0.2975 | 228.85 | 216.66 |
| SOAT1 | M2 | M | Tan et al., 2014 | 0.0003 | 228.85 | 216.66 |
| SPOCK1 | M2 | M | Tan et al., 2014 | 0.9936 | 228.85 | 216.66 |
| SRF | M2 | NA | dbEMT | 0.6437 | 28.85 | 216.66 |
| STAT3 | M2 | NA | dbEMT | 0.1408 | 115 | 171.43 |
| SYNM | M2 | M | Tan et al., 2014 | 0.001 | 35 | 37.64 |
| TAB1 | M2 | NA | dbEMT | n/a | n/a | n/a |
| TBX3 | M2 | NA | dbEMT | 1.80E-10 | 228.85 | 216.66 |
| TCF4 | M2 | M | Tan et al., 2014 | 0.9205 | 29 | 43 |
| TP53 | M2 | NA | dbEMT | 0.0005 | 216.66 | 191.21 |
| TRPS1 | M2 | NA | dbEMT | 0.2083 | 28.96 | 44 |
| VCAN | M2 | M | Tan et al., 2014 | 0.2395 | 228.85 | 216.66 |
| VIM | M2 | M | Tan et al., 2014 | 0.0949 | 185.16 | 216.66 |
| WNT5A | M2 | NA | dbEMT | 0.433 | 228.85 | 216.66 |

| | | | | | | |
|---|---|---|---|---|---|---|
| ZEB1 | M2 | M | Tan et al., 2014 | 1.10E-06 | 216.66 | 228.85 |
| ZEB2 | M2 | M | Tan et al., 2014 | 0.0284 | 44 | 54 |
| ZFPM2 | M2 | M | Tan et al., 2014 | 0.0002 | 216.66 | 228.85 |
| AKAP12 | M3 | M | Tan et al., 2014 | 0.0004 | 34.8 | 37 |
| ANPEP | M3 | NA | dbEMT | 0.0793 | 228.85 | 216.66 |
| AR | M3 | NA | dbEMT | 4.20E-05 | 44.6 | 53.04 |
| ASPN | M3 | M | Tan et al., 2014 | 0.2608 | 228.85 | 191.21 |
| AXL | M3 | M | Tan et al., 2014 | 0.0004 | 27 | 45.6 |
| BGN | M3 | M | Tan et al., 2014 | 0.721 | 43.7 | 56.04 |
| BMI1 | M3 | NA | dbEMT | 0.7753 | 228.85 | 216.66 |
| BMP4 | M3 | NA | dbEMT | 0.058 | 43 | 56.58 |
| CCL8 | M3 | M | Tan et al., 2014 | 0.0025 | 23.82 | 61 |
| CMTM8 | M3 | NA | dbEMT | 0.9132 | 216.66 | 191.21 |
| COL8A1 | M3 | NA | dbEMT | 5.40E-06 | 42.96 | 55.43 |
| CTSZ | M3 | NA | dbEMT | 0.0819 | 228.85 | 216.66 |
| CYP1B1 | M3 | M | Tan et al., 2014 | 0.563 | 54.77 | 44 |
| DDX5 | M3 | NA | dbEMT | 3.30E-05 | 28.8 | 43.7 |
| DLX4 | M3 | NA | dbEMT | 0.0059 | 228.85 | 216.66 |
| EDNRA | M3 | NA | dbEMT | 0.7264 | 228.85 | 191.21 |
| EFEMP1 | M3 | M | Tan et al., 2014 | 0.013 | 48 | 48 |
| EPB41L5 | M3 | NA | dbEMT | 1.00E-16 | 147 | 171.43 |
| ERBB2IP | M3 | NA | dbEMT | 0.0018 | 50 | 47 |
| EVI2A | M3 | M | Tan et al., 2014 | 0.0026 | 26 | 49.2 |
| FBLN1 | M3 | M | Tan et al., 2014 | 3.60E-05 | 216.66 | 228.85 |
| FERMT2 | M3 | M | Tan et al., 2014 | 0.0285 | 216.66 | 228.85 |
| FGFR1 | M3 | NA | dbEMT | 0.0002 | 228.85 | 216.66 |
| FOXC2 | M3 | E | Tan et al., 2014 | 0.0427 | 30 | 40.71 |
| FXYD6 | M3 | M | Tan et al., 2014 | 0.0003 | 228.85 | 216.66 |
| GAS1 | M3 | M | Tan et al., 2014 | 0.3391 | 228.85 | 216.66 |
| GLIPR1 | M3 | M | Tan et al., 2014 | 1.80E-05 | 184.04 | 216.66 |
| GLRX | M3 | NA | dbEMT | 0.9264 | 228.85 | 216.66 |
| GLYR1 | M3 | M | Tan et al., 2014 | 1.30E-09 | 31.18 | 40.44 |
| GNG11 | M3 | M | Tan et al., 2014 | 0.0043 | 228.85 | 191.21 |
| GPM6B | M3 | M | Tan et al., 2014 | 0.2889 | 48 | 48 |
| HGF | M3 | NA | dbEMT | 0.0058 | 30 | 40.8 |
| HNMT | M3 | E | Tan et al., 2014 | 0.0082 | 53.1 | 44.61 |
| HOXA10 | M3 | NA | dbEMT | 0.038 | 42 | 59 |
| IGFBP3 | M3 | NA | dbEMT | 0.0446 | 38 | 65.1 |
| ISLR | M3 | M | Tan et al., 2014 | 0.0464 | 52.54 | 44.22 |
| LAMA5 | M3 | NA | dbEMT | 0.0002 | 191.21 | 228.85 |
| LRP6 | M3 | NA | dbEMT | 0.006 | 36 | 36.04 |
| MAFB | M3 | M | Tan et al., 2014 | 0.1534 | 29 | 42 |
| MANSC1 | M3 | E | Tan et al., 2014 | 0.6974 | 228.85 | 216.66 |
| MAP3K3 | M3 | NA | dbEMT | 0.0029 | 44 | 54 |
| MAPK14 | M3 | NA | dbEMT | 0.0918 | 40.3 | 57.3 |
| MAPK7 | M3 | NA | dbEMT | 0.0005 | 37 | 34.8 |
| MFAP4 | M3 | M | Tan et al., 2014 | 5.80E-09 | 43 | 58 |
| MIR15B | M3 | NA | dbEMT | n/a | n/a | n/a |
| MIR193A | M3 | NA | dbEMT | n/a | n/a | n/a |
| MIR34A | M3 | NA | dbEMT | n/a | n/a | n/a |

| | | | | | | |
|---|---|---|---|---|---|---|
| MMP7 | M3 | NA | dbEMT | 0.096 | 129 | 171.43 |
| MUC16 | M3 | NA | dbEMT | 0.9594 | 47 | 50 |
| MYLK | M3 | M | Tan et al., 2014 | 0.7165 | 54 | 43 |
| NFIC | M3 | NA | dbEMT | 0.0253 | 216.66 | 228.85 |
| PLXNC1 | M3 | M | Tan et al., 2014 | 0.2577 | 184.04 | 216.66 |
| PLXND1 | M3 | NA | dbEMT | 0.0492 | 35 | 36.9 |
| PPARG | M3 | NA | dbEMT | 0.0007 | 51 | 45.84 |
| PTPN14 | M3 | NA | dbEMT | 0.4434 | 35.98 | 71 |
| PTRF | M3 | M | Tan et al., 2014 | 0.0004 | 216.66 | 228.85 |
| QKI | M3 | M | Tan et al., 2014 | 2.00E-05 | 29 | 42 |
| RAF1 | M3 | NA | dbEMT | 0.2984 | 228.85 | 216.66 |
| ROCK2 | M3 | NA | dbEMT | 0.0786 | 228.85 | 216.66 |
| S100A4 | M3 | NA | dbEMT | 0.0437 | 216.66 | 228.85 |
| SERPING1 | M3 | M | Tan et al., 2014 | 1.30E-08 | 35 | 36.9 |
| SOBP | M3 | M | Tan et al., 2014 | 0.6293 | 51 | 45.84 |
| SON | M3 | NA | dbEMT | 0.6585 | 35.98 | 71 |
| SPARCL1 | M3 | M | Tan et al., 2014 | 1.00E-13 | 216.66 | 228.85 |
| SRGN | M3 | M | Tan et al., 2014 | 0.2041 | 29 | 42 |
| TEAD1 | M3 | NA | dbEMT | 0.0005 | 228.85 | 216.66 |
| TGFB2 | M3 | NA | dbEMT | 0.3875 | 228.85 | 216.66 |
| TGFB3 | M3 | NA | dbEMT | 1.50E-11 | 228.85 | 216.66 |
| TIMP1 | M3 | NA | dbEMT | 0.0414 | 228.85 | 216.66 |
| TMEM158 | M3 | M | Tan et al., 2014 | 0.0014 | 57.27 | 41.64 |
| TRIM31 | M3 | E | Tan et al., 2014 | 3.40E-06 | 191.21 | 216.66 |
| TWIST1 | M3 | M | Tan et al., 2014 | 0.7013 | 52 | 46 |
| TWIST2 | M3 | M | Tan et al., 2014 | 0.0003 | 30 | 43 |
| VWCE | M3 | NA | dbEMT | 0.1218 | 34.82 | 37 |
| WISP2 | M3 | NA | dbEMT | 0.0002 | 40.8 | 58 |
| WNT6 | M3 | NA | dbEMT | 0.0979 | 228.85 | 216.66 |
| ZCCHC24 | M3 | M | Tan et al., 2014 | 7.40E-12 | 36 | 71.04 |
| BAG2 | Other M | M | Tan et al., 2014 | 0.6068 | 52.2 | 45.9 |
| CSNK2B | Other M | NA | dbEMT | 1.20E-09 | 65.04 | 37.35 |
| CTBP1 | Other M | NA | dbEMT | 0.5356 | 191.21 | 216.66 |
| HMGB3 | Other M | NA | dbEMT | 2.60E-08 | 48 | 27.4 |
| KHDRBS1 | Other M | NA | dbEMT | 0.0008 | 58.52 | 41.65 |
| LEFTY1 | Other M | NA | dbEMT | 1.60E-08 | 228.85 | 216.66 |
| MAPK1 | Other M | NA | dbEMT | 0.1309 | 35.98 | 36.9 |
| MS4A4A | Other M | M | Tan et al., 2014 | 0.496 | 36 | 35.98 |
| MS4A6A | Other M | M | Tan et al., 2014 | 0.0004 | 216.66 | 228.85 |
| MTDH | Other M | NA | dbEMT | 3.50E-10 | 216.66 | 184.04 |
| NF1 | Other M | NA | dbEMT | 3.00E-11 | 191.21 | 216.66 |

**Supplementary Table 3** Enrichment of EMT genes among gene upregulated by EMT regulators in Tauble et al.

| **Down Regulated Genes (FC < 2)** | | | | | |
|---|---|---|---|---|---|
| | TGF-β | Twist | Snail | Gsc | sh-Ecad |
| E | 78 (*p* = 2.1e-61) | 110 (*p* = 4.8e-80) | 116 (*p* = 1.1e-77) | 114 (*p* = 1.9e-55) | 148 (*p* = 1.9e-94) |
| M | 3 (*p* = 0.47) | 7 (*p* = 0.67) | 6 (*p* = 0.47) | 8 (*p* = 0.41) | 14 (*p* = 0.78) |
| **Up Regulated Genes (FC > 2)** | | | | | |
| | TGF-β | Twist | Snail | Gsc | sh-Ecad |
| E | 7 (*p* = 1) | 12 (*p* = 0.77) | 12 (*p* = 1) | 21 (*p* = 0.82) | 4 (*p* = 1) |
| M | 70 (*p* = 1.1e-51) | 88 (*p* = 7.7e-52) | 94 (*p* = 1.5e-61) | 103 (*p* = 1.3e-45) | 82 (*p* = 2.8e-48) |
| M1 | 35 (*p* = 2.8e-25) | 43 (*p* = 1.9e-24) | 46 (*p* = 6.1e-29) | 40 (*p* = 1.3e-13) | 41 (*p* = 9.7e-24) |
| M2 | 23 (*p* = 6.1e-21) | 24 (*p* = 1.9e-16) | 28 (*p* = 5.5e-22) | 37 (*p* = 1.1e-24) | 23 (*p* = 4.5e-16) |
| M3 | 12 (*p* = 2.4e-8) | 21 (*p* = 6.4e-14) | 20 (*p* = 1.7e-13) | 26 (*p* = 4.9e-14) | 18 (*p* = 2.1e-11) |

**Supplementary Table 4.** List of GO Terms enriched in all epithelial genes

| GO | P.adjust | Short Description |
|---|---|---|
| GO:0016324 | 2.05E-08 | apical plasma membrane |
| GO:0016328 | 3.61E-08 | lateral plasma membrane |
| GO:0005911 | 7.14E-08 | cell-cell junction |
| GO:0016323 | 9.88E-08 | basolateral plasma membrane |
| GO:0070268 | 1.95E-07 | cornification |
| GO:0007219 | 1.95E-07 | Notch signaling pathway |
| GO:0030057 | 4.83E-07 | desmosome |
| GO:0098609 | 2.15E-06 | cell-cell adhesion |
| GO:0030216 | 3.08E-06 | keratinocyte differentiation |
| GO:0005923 | 4.08E-06 | bicellular tight junction |
| GO:0016477 | 4.08E-06 | cell migration |
| GO:0071944 | 9.66E-06 | cell periphery |
| GO:0007179 | 1.77E-05 | transforming growth factor beta receptor signaling pathway |
| GO:0008013 | 2.55E-05 | beta-catenin binding |
| GO:0001934 | 4.79E-05 | positive regulation of protein phosphorylation |
| GO:0001533 | 5.06E-05 | cornified envelope |
| GO:0050679 | 5.06E-05 | positive regulation of epithelial cell proliferation |
| GO:0043406 | 5.66E-05 | positive regulation of MAP kinase activity |
| GO:0090303 | 5.66E-05 | positive regulation of wound healing |
| GO:0002934 | 6.03E-05 | desmosome organization |
| GO:0051897 | 8.28E-05 | positive regulation of protein kinase B signaling |
| GO:0060749 | 8.28E-05 | mammary gland alveolus development |
| GO:0005913 | 8.41E-05 | cell-cell adherens junction |
| GO:0007173 | 0.000113066 | epidermal growth factor receptor signaling pathway |
| GO:0016327 | 0.000113066 | apicolateral plasma membrane |
| GO:0050839 | 0.000152801 | cell adhesion molecule binding |
| GO:0005829 | 0.000156763 | cytosol |
| GO:0038132 | 0.000209644 | neuregulin binding |
| GO:0086083 | 0.000209644 | cell adhesive protein binding involved in bundle of His cell-Purkinje myocyte communication |
| GO:0038128 | 0.000283371 | ERBB2 signaling pathway |
| GO:0061436 | 0.000304381 | establishment of skin barrier |
| GO:0098911 | 0.000308222 | regulation of ventricular cardiac muscle cell action potential |
| GO:0000165 | 0.000315781 | MAPK cascade |
| GO:0043235 | 0.000409511 | receptor complex |
| GO:0008544 | 0.000455941 | epidermis development |
| GO:0071773 | 0.000497706 | cellular response to BMP stimulus |
| GO:0070830 | 0.000497706 | bicellular tight junction assembly |
| GO:0005154 | 0.000497706 | epidermal growth factor receptor binding |

| | | |
|---|---|---|
| GO:0086073 | 0.000501582 | bundle of His cell-Purkinje myocyte adhesion involved in cell communication |
| GO:0014704 | 0.000552585 | intercalated disc |
| GO:0009925 | 0.000583926 | basal plasma membrane |
| GO:0031528 | 0.000643276 | microvillus membrane |
| GO:0005887 | 0.001074063 | integral component of plasma membrane |
| GO:1900740 | 0.001076022 | positive regulation of protein insertion into mitochondrial membrane involved in apoptotic signaling pathway |
| GO:0005198 | 0.001308149 | structural molecule activity |
| GO:0060672 | 0.001534535 | epithelial cell morphogenesis involved in placental branching |
| GO:0032496 | 0.001561501 | response to lipopolysaccharide |
| GO:0048709 | 0.001580009 | oligodendrocyte differentiation |
| GO:0032091 | 0.001580009 | negative regulation of protein binding |
| GO:0032147 | 0.001880503 | activation of protein kinase activity |
| GO:0070372 | 0.001880503 | regulation of ERK1 and ERK2 cascade |
| GO:0070371 | 0.001880503 | ERK1 and ERK2 cascade |
| GO:0045121 | 0.001962464 | membrane raft |
| GO:0008134 | 0.002462225 | transcription factor binding |
| GO:0071437 | 0.002820133 | invadopodium |
| GO:0030054 | 0.002820133 | cell junction |
| GO:0045747 | 0.003147601 | positive regulation of Notch signaling pathway |
| GO:0031424 | 0.003147601 | keratinization |
| GO:0042127 | 0.003362169 | regulation of cell population proliferation |
| GO:0030674 | 0.003536732 | protein binding, bridging |
| GO:0005200 | 0.00376746 | structural constituent of cytoskeleton |
| GO:0098641 | 0.003807955 | cadherin binding involved in cell-cell adhesion |
| GO:0023019 | 0.003807955 | signal transduction involved in regulation of gene expression |
| GO:0045216 | 0.003807955 | cell-cell junction organization |
| GO:0043547 | 0.004558012 | positive regulation of GTPase activity |
| GO:0005112 | 0.00490046 | Notch binding |
| GO:0030855 | 0.005509904 | epithelial cell differentiation |
| GO:0046934 | 0.006037726 | phosphatidylinositol-4,5-bisphosphate 3-kinase activity |
| GO:0051496 | 0.006466229 | positive regulation of stress fiber assembly |
| GO:0048754 | 0.007617998 | branching morphogenesis of an epithelial tube |
| GO:0001228 | 0.007830327 | DNA-binding transcription activator activity, RNA polymerase II-specific |
| GO:0034332 | 0.008195817 | adherens junction organization |
| GO:0086091 | 0.008195817 | regulation of heart rate by cardiac conduction |
| GO:2000146 | 0.008590302 | negative regulation of cell motility |
| GO:0019899 | 0.008836847 | enzyme binding |
| GO:0090263 | 0.008841438 | positive regulation of canonical Wnt signaling pathway |
| GO:0010468 | 0.008841438 | regulation of gene expression |
| GO:0042104 | 0.008841438 | positive regulation of activated T cell proliferation |

| | | |
|---|---|---|
| GO:0031982 | 0.009001695 | vesicle |
| GO:0071504 | 0.009449414 | cellular response to heparin |
| GO:0030618 | 0.009449414 | transforming growth factor beta receptor, pathway-specific cytoplasmic mediator activity |
| GO:0009967 | 0.010621008 | positive regulation of signal transduction |
| GO:0034236 | 0.011122653 | protein kinase A catalytic subunit binding |
| GO:0030154 | 0.012686211 | cell differentiation |
| GO:0051384 | 0.012972101 | response to glucocorticoid |
| GO:0010469 | 0.013970766 | regulation of signaling receptor activity |
| GO:0030307 | 0.014460919 | positive regulation of cell growth |
| GO:0051017 | 0.014460919 | actin filament bundle assembly |
| GO:0045668 | 0.014460919 | negative regulation of osteoblast differentiation |
| GO:0044319 | 0.014460919 | wound healing, spreading of cells |
| GO:0060487 | 0.016323064 | lung epithelial cell differentiation |
| GO:1902949 | 0.016323064 | positive regulation of tau-protein kinase activity |
| GO:0060056 | 0.016323064 | mammary gland involution |
| GO:0045840 | 0.016644153 | positive regulation of mitotic nuclear division |
| GO:2000145 | 0.016644153 | regulation of cell motility |
| GO:0005882 | 0.017306919 | intermediate filament |
| GO:0030949 | 0.017306919 | positive regulation of vascular endothelial growth factor receptor signaling pathway |
| GO:0001658 | 0.017306919 | branching involved in ureteric bud morphogenesis |
| GO:0022408 | 0.017306919 | negative regulation of cell-cell adhesion |
| GO:0060441 | 0.017306919 | epithelial tube branching involved in lung morphogenesis |
| GO:2000060 | 0.017306919 | positive regulation of ubiquitin-dependent protein catabolic process |
| GO:0035326 | 0.017306919 | enhancer binding |
| GO:0048471 | 0.017395121 | perinuclear region of cytoplasm |
| GO:0072659 | 0.018286409 | protein localization to plasma membrane |
| GO:0004888 | 0.018646687 | transmembrane signaling receptor activity |
| GO:0051781 | 0.021713298 | positive regulation of cell division |
| GO:0044212 | 0.021713298 | transcription regulatory region DNA binding |
| GO:0007229 | 0.023968789 | integrin-mediated signaling pathway |
| GO:0071141 | 0.023968789 | SMAD protein complex |
| GO:0035655 | 0.023968789 | interleukin-18-mediated signaling pathway |
| GO:0050731 | 0.023968789 | positive regulation of peptidyl-tyrosine phosphorylation |
| GO:0060687 | 0.023968789 | regulation of branching involved in prostate gland morphogenesis |
| GO:0005138 | 0.023968789 | interleukin-6 receptor binding |
| GO:0030056 | 0.023968789 | hemidesmosome |
| GO:0045617 | 0.023968789 | negative regulation of keratinocyte differentiation |
| GO:0061384 | 0.023968789 | heart trabecula morphogenesis |
| GO:0043231 | 0.023981011 | intracellular membrane-bounded organelle |
| GO:0007166 | 0.024190272 | cell surface receptor signaling pathway |

| | | |
|---|---|---|
| GO:0042531 | 0.024911995 | positive regulation of tyrosine phosphorylation of STAT protein |
| GO:0030676 | 0.025334535 | Rac guanyl-nucleotide exchange factor activity |
| GO:2000249 | 0.025334535 | regulation of actin cytoskeleton reorganization |
| GO:0008584 | 0.025892253 | male gonad development |
| GO:0007565 | 0.025892253 | female pregnancy |
| GO:0046777 | 0.027322274 | protein autophosphorylation |
| GO:0048557 | 0.030290213 | embryonic digestive tract morphogenesis |
| GO:0006970 | 0.030290213 | response to osmotic stress |
| GO:0031663 | 0.030290213 | lipopolysaccharide-mediated signaling pathway |
| GO:0005044 | 0.030290213 | scavenger receptor activity |
| GO:0030279 | 0.030290213 | negative regulation of ossification |
| GO:0000187 | 0.032035464 | activation of MAPK activity |
| GO:0071144 | 0.032035464 | heteromeric SMAD protein complex |
| GO:0061302 | 0.032035464 | smooth muscle cell-matrix adhesion |
| GO:0038092 | 0.032035464 | nodal signaling pathway |
| GO:0043588 | 0.032035464 | skin development |
| GO:0004450 | 0.032035464 | isocitrate dehydrogenase (NADP+) activity |
| GO:0006097 | 0.032035464 | glyoxylate cycle |
| GO:0048340 | 0.032035464 | paraxial mesoderm morphogenesis |
| GO:0070878 | 0.032035464 | primary miRNA binding |
| GO:0010482 | 0.032035464 | regulation of epidermal cell division |
| GO:0062043 | 0.032035464 | positive regulation of cardiac epithelial to mesenchymal transition |
| GO:0001773 | 0.032035464 | myeloid dendritic cell activation |
| GO:0071639 | 0.032035464 | positive regulation of monocyte chemotactic protein-1 production |
| GO:0003169 | 0.032035464 | coronary vein morphogenesis |
| GO:0022612 | 0.032035464 | gland morphogenesis |
| GO:0005172 | 0.032035464 | vascular endothelial growth factor receptor binding |
| GO:0071506 | 0.032035464 | cellular response to mycophenolic acid |
| GO:0005088 | 0.032963162 | Ras guanyl-nucleotide exchange factor activity |
| GO:0007266 | 0.033093313 | Rho protein signal transduction |
| GO:0043034 | 0.033813585 | costamere |
| GO:0008283 | 0.035449084 | cell proliferation |
| GO:0032570 | 0.035449084 | response to progesterone |
| GO:0046332 | 0.036079675 | SMAD binding |
| GO:0032755 | 0.039660936 | positive regulation of interleukin-6 production |
| GO:0005070 | 0.039660936 | SH3/SH2 adaptor activity |
| GO:0045104 | 0.039843233 | intermediate filament cytoskeleton organization |
| GO:0033627 | 0.039843233 | cell adhesion mediated by integrin |
| GO:0001889 | 0.040235914 | liver development |
| GO:0005794 | 0.040713061 | Golgi apparatus |
| GO:0045741 | 0.041122468 | positive regulation of epidermal growth factor-activated receptor activity |
| GO:0035313 | 0.041122468 | wound healing, spreading of epidermal cells |

| | | |
|---|---|---|
| GO:0060253 | 0.041122468 | negative regulation of glial cell proliferation |
| GO:0032872 | 0.041122468 | regulation of stress-activated MAPK cascade |
| GO:0032835 | 0.041122468 | glomerulus development |
| GO:0033689 | 0.041122468 | negative regulation of osteoblast proliferation |
| GO:0004714 | 0.041122468 | transmembrane receptor protein tyrosine kinase activity |
| GO:0005912 | 0.041122468 | adherens junction |
| GO:0048469 | 0.042765817 | cell maturation |
| GO:0001843 | 0.044774075 | neural tube closure |
| GO:0043407 | 0.04804062 | negative regulation of MAP kinase activity |

**Supplementary Table 5.** List of GO Terms enriched in epithelial genes which are differentially expressed in response to both TGF-β and ZEB1.

| GO | P.adjust | Short Description |
|---|---|---|
| GO:0070062 | 2.82E-08 | extracellular exosome |
| GO:0005886 | 0.00021 | plasma membrane |
| GO:0070268 | 0.00038 | cornification |
| GO:0060672 | 0.000546 | epithelial cell morphogenesis involved in placental branching |
| GO:0045296 | 0.000617 | cadherin binding |
| GO:0050839 | 0.000617 | cell adhesion molecule binding |
| GO:0098609 | 0.000617 | cell-cell adhesion |
| GO:0002934 | 0.000968 | desmosome organization |
| GO:0016323 | 0.000968 | basolateral plasma membrane |
| GO:0005923 | 0.003392 | bicellular tight junction |
| GO:0098641 | 0.003531 | cadherin binding involved in cell-cell adhesion |
| GO:0030057 | 0.010324 | desmosome |
| GO:0005913 | 0.010391 | cell-cell adherens junction |
| GO:0016328 | 0.010391 | lateral plasma membrane |
| GO:0005911 | 0.011385 | cell-cell junction |
| GO:0005154 | 0.021427 | epidermal growth factor receptor binding |
| GO:0031424 | 0.031024 | keratinization |
| GO:0034332 | 0.031024 | adherens junction organization |
| GO:0001843 | 0.03743 | neural tube closure |
| GO:0072659 | 0.040151 | protein localization to plasma membrane |

**Supplementary Table 6.** Gene sets enriched in M-gene clusters

| M-gene Cluster | q-value | GSEA Group Description |
|---|---|---|
| M1 | 3.96E-55 | Genes up-regulated in invasive ductal carcinoma (IDC) relative to ductal carcinoma in situ (DCIS, non-invasive). |
| M1 | 3.04E-43 | Genes specifically up-regulated in Cluster IIb of urothelial cell carcinom (UCC) tumors. |
| M1 | 1.44E-30 | Genes down-regulated in luminal-like breast cancer cell lines compared to the mesenchymal-like ones. |
| M1 | 5.57E-29 | Genes down-regulated in the luminal B subtype of breast cancer. |
| M1 | 1.17E-28 | Genes down-regulated in prostate cancer samples. |
| M1 | 5.77E-26 | Genes down-regulated in the Kras2LA mouse lung cancer model with mutated KRAS [GeneID=3845]. |
| M1 | 4.41E-21 | Genes up-regulated in the normal-like subtype of breast cancer. |
| M1 | 1.68E-17 | Genes down-regulated in papillary thyroid carcinoma (PTC) compared to normal tissue. |
| M1 | 2.38E-17 | Genes from 'subtype S1' signature of hepatocellular carcinoma (HCC): aberrant activation of the WNT signaling pathway. |
| M1 | 4.98E-17 | Invasiveness signature resulting from cancer cell/microenvironment interaction. |
| M1 | 3.73E-14 | Up-regulated genes distinguishing between two subtypes of gastric cancer: advanced (AGC) and early (EGC). |
| M1 | 1.12E-11 | Genes whose expression in suboptimally debulked ovarian tumors is associated with survival prognosis. |
| M1 | 5.72E-11 | Genes up-regulated in pancreatic ductal adenocarcinoma (PDAC) identified in a meta analysis across four independent studies. |
| M1 | 5.23E-10 | Genes up-regulated in basal subtype of breast cancer samles. |
| M1 | 1.47E-09 | Genes down-regulated in luminal-like breast cancer cell lines compared to the basal-like ones. |
| M1 | 1.68E-09 | Genes down-regulated in basal subtype of breast cancer samles. |
| M1 | 3.04E-09 | Cluster 4: selected stromal genes clustered together across breast cancer samples. |
| M1 | 4.53E-09 | Genes up-regulated in atypical ductal hyperplastic tissues from patients with (ADHC) breast cancer vs those without the cancer (ADH). |
| M2 | 9.18E-34 | Genes up-regulated in invasive ductal carcinoma (IDC) relative to ductal carcinoma in situ (DCIS, non-invasive). |
| M2 | 4.09E-24 | Genes down-regulated in luminal-like breast cancer cell lines compared to the mesenchymal-like ones. |
| M2 | 1.05E-13 | Genes specifically up-regulated in Cluster IIb of urothelial cell carcinom (UCC) tumors. |
| M2 | 1.05E-13 | Up-regulated genes distinguishing between two subtypes of gastric cancer: advanced (AGC) and early (EGC). |
| M2 | 6.19E-13 | Invasiveness signature resulting from cancer cell/microenvironment interaction. |

| | | |
|---|---|---|
| M2 | 6.19E-13 | Genes down-regulated in luminal-like breast cancer cell lines compared to the basal-like ones. |
| M2 | 2.43E-10 | Genes up-regulated in pancreatic ductal adenocarcinoma (PDAC) identified in a meta analysis across four independent studies. |
| M2 | 3.49E-10 | Genes down-regulated in mucinous ovarian carcinoma tumors of low malignant potential (LMP) compared to normal ovarian surface epithelium tissue. |
| M2 | 5.29E-09 | Genes down-regulated in prostate cancer samples. |
| M2 | 3.13E-08 | Genes from 'subtype S1' signature of hepatocellular carcinoma (HCC): aberrant activation of the WNT signaling pathway. |
| M2 | 2.58E-07 | Genes down-regulated in the luminal B subtype of breast cancer. |
| M2 | 6.14E-07 | Genes up-regulated in the normal-like subtype of breast cancer. |
| M2 | 1.05E-06 | Genes whose expression in suboptimally debulked ovarian tumors is associated with survival prognosis. |
| M2 | 1.96E-06 | Genes up-regulated in serrated vs conventional colorectal carcinoma (CRC) samples. |
| M2 | 2.05E-06 | Genes whose DNA was methylated both in primary tumors and across a panel of cancer cell lines. |
| M2 | 2.81E-06 | Genes down-regulated in the Kras2LA mouse lung cancer model with mutated KRAS [GeneID=3845]. |
| M3 | 1.68E-25 | Genes up-regulated in invasive ductal carcinoma (IDC) relative to ductal carcinoma in situ (DCIS, non-invasive). |
| M3 | 2.69E-15 | Genes specifically up-regulated in Cluster IIb of urothelial cell carcinom (UCC) tumors. |
| M3 | 4.03E-12 | Up-regulated genes distinguishing between two subtypes of gastric cancer: advanced (AGC) and early (EGC). |
| M3 | 7.93E-12 | Genes down-regulated in the luminal B subtype of breast cancer. |
| M3 | 9.19E-12 | Genes down-regulated in luminal-like breast cancer cell lines compared to the mesenchymal-like ones. |
| M3 | 2.67E-09 | Genes down-regulated in medullary breast cancer (MBC) relative to ductal breast cancer (DBD). |
| M3 | 2.77E-08 | Genes down-regulated in the Kras2LA mouse lung cancer model with mutated KRAS [GeneID=3845]. |
| M3 | 6.46E-08 | Genes down-regulated in prostate cancer samples. |
| M3 | 1.96E-07 | Genes down-regulated in basal subtype of breast cancer samles. |
| M3 | 1.96E-07 | Genes up-regulated in prostate cancer samples from African-American patients compared to those from the European-American patients. |
| M3 | 6.73E-07 | Genes up-regulated in the normal-like subtype of breast cancer. |
| M3 | 2.90E-06 | Genes down-regulated in mucinous ovarian carcinoma tumors of low malignant potential (LMP) compared to normal ovarian surface epithelium tissue. |
| M3 | 6.84E-06 | Genes up-regulated in basal subtype of breast cancer samles. |
| M3 | 1.40E-05 | Early prostate development genes (up-regulated at 48 hr dihydrotestosterone [PubChem=10635]) which are also up-regulated in high grade prostatic intraepithelial neoplasia (PIN) vs invasive cancer. |
| M3 | 1.47E-05 | Genes up-regulated in circulating endothelial cells (CEC) from cancer patients compared to those from healthy donors. |

| | | |
|---|---|---|
| M3 | 1.56E-05 | Genes up-regulated in the luminal A subtype of breast cancer. |
| M3 | 1.97E-05 | Top 200 marker genes down-regulated in the 'CTNNB1' subclass of hepatocellular carcinoma (HCC); characterized by activated CTNNB1 [GeneID=1499]. |
| M3 | 2.76E-05 | Genes down-regulated in prostate cancer vs benign prostate tissue, based on a meta-analysis of five gene expression profiling studies. |
| M3 | 3.16E-05 | Genes up-regulated in papillary thyroid carcinoma (PTC) compared to normal tissue. |
| M3 | 3.90E-05 | Genes from the brain cancer stem (cancer stem cell, CSC) signature. |
| M3 | 1.07E-04 | Genes down-regulated in oncocytic follicular carcinoma (FTC) vs mitochondrial-rich papillary carcinoma (PTC) types of thyroid cancer. |
| M3 | 1.39E-04 | Invasiveness signature resulting from cancer cell/microenvironment interaction. |

**Supplementary Table 7.** List of primer sequences

| Target gene | Forward primer | Reverse primer |
|---|---|---|
| *CDH1* | AAAGGCCCATTTCCTAAAAACCT | TGCGTTCTCTATCCAGAGGCT |
| *SNAI1* | TCGGAAGCCTAACTACAGCGA | AGATGAGCATTGGCAGCGAG |
| *FN1* | AGGAAGCCGAGGTTTTAACTG | AGGACGCTCATAAGTGTCACC |
| *TWIST1* | GTCCGCAGTCTTACGAGGAG | GCTTGAGGGTCTGAATCTTGCT |
| *GRHL2* | GGGGAAGAGCAACGAGTGG | GGGGAAGAGCAACGAGTGG |
| *MMP3* | CTGGACTCCGACACTCTGGA | CAGGAAAGGTTCTGAAGTGACC |
| *VIM* | GACGCCATCAACACCGAGTT | CTTTGTCGTTGGTTAGCTGGT |
| *GAPDH* | GAAGGTGAAGGTCGGAGT | GAAGATGGTGATGGGATTTC |
| *ACTB* | CTTCTACAATGAGCTGCGTG | GGGTGTTGAAGGTCTCAAAC |

**Supplementary Table 8.** Parameter values for mathematical models

| Parameter | Description | Value |
|---|---|---|
| $\gamma_{\mathrm{TGF\beta}}$ | Timescale of TGF-β | 1 |
| $\gamma_{\mathrm{ZEB1}}$ | Timescale of ZEB1 | 1 |
| $\gamma_{\mathrm{X}}$ | Timescale of factor X (e.g. OVOL2, miR200) | 1 |
| $\gamma_{\mathrm{M1}}$ | Timescale of M1 genes | 1 |
| $\gamma_{\mathrm{M2}}$ | Timescale of M2 genes | 1 |
| $\gamma_{\mathrm{M3}}$ | Timescale of M3 genes | 1 |
| $\gamma_{\mathrm{E}}$ | Timescale of E genes | 1 |
| $\sigma_{\mathrm{TGF\beta}}$ | Steepness of sigmoidal function of TGF-β | 10 |
| $\sigma_{\mathrm{ZEB1}}$ | Steepness of sigmoidal function of ZEB1 | 10 |
| $\sigma_{\mathrm{X}}$ | Steepness of sigmoidal function of factor X | 10 |
| $\sigma_{\mathrm{M1}}$ | Steepness of sigmoidal function of M1 genes | 10 |
| $\sigma_{\mathrm{M2}}$ | Steepness of sigmoidal function of M2 genes | 10 |
| $\sigma_{\mathrm{M3}}$ | Steepness of sigmoidal function of M3 genes | 10 |
| $\sigma_{\mathrm{E}}$ | Steepness of sigmoidal function of E genes | 10 |
| $w^0_{\mathrm{TGF\beta}}$ | Basal production rate of TGF-β | -1 |
| $w^0_{\mathrm{ZEB1}}$ | Basal production rate of ZEB1 | 0.2 |
| $w^0_{\mathrm{X}}$ | Basal production rate of factor X | 0.5 |
| $w^0_{\mathrm{M1}}$ | Basal production rate of M1 genes | -0.5 |
| $w^0_{\mathrm{M2}}$ | Basal production rate of M2 genes | -0.5 |
| $w^0_{\mathrm{M3}}$ | Basal production rate of M3 genes | -0.5 |
| $w^0_{\mathrm{E}}$ | Basal production rate of E genes | 0.5 |
| $\omega_{\mathrm{TGF\beta \to ZEB1}}$ | Activation strength of TGF-β on ZEB1 | 1 |
| $\omega_{\mathrm{ZEB1 \to TGF\beta}}$ | Activation strength of ZEB1 on TGF-β | 1 |
| $\omega_{\mathrm{ZEB1 \to X}}$ | Inhibition strength of ZEB1 on factor X | -0.6 |
| $\omega_{\mathrm{X \to ZEB1}}$ | Inhibition strength of X on factor ZEB1 | -0.6 |
| $\omega_{\mathrm{ZEB1 \to E}}$ | Inhibition strength of ZEB1 on E genes | -1 |
| $\omega_{\mathrm{TGF\beta \to M1}}$ | Activation strength of TGF-β on M1 genes | 0.5* |
| $\omega_{\mathrm{ZEB1 \to M1}}$ | Activation strength of ZEB1 on M1 genes | 0.1** |
| $\omega_{\mathrm{TGF\beta \to M2}}$ | Activation strength of TGF-β on M2 genes | 1 |
| $\omega_{\mathrm{ZEB1 \to M2}}$ | Activation strength of ZEB1 on M2 genes | 1 |
| $\omega_{\mathrm{ZEB1 \to M3}}$ | Activation strength of ZEB1 on M3 genes | 0.7 |

* Total activation is saturated at 0.4    ** Total activation is saturated at 0.1
